# Data Analyses and new Findings Indicate a Primordial Neurotropic Pathogen Evolved into Infectious Causes of Several CNS Neurodegenerative Diseases

**DOI:** 10.1101/232074

**Authors:** William J. Todd, Lidiya Dubytska, Peter J. Mottram, Xiaochu Wu, Yuliya Y. Sokolova

**Affiliations:** Department of Pathobiological Sciences, Louisiana State University School of Veterinary Medicine, Baton Rouge, LA 70803, USA.; Department of Biology and Chemistry, Southern University, Baton Rouge, LA 70813, USA.; Department of Comparative Biomedical Sciences, Louisiana State University School of Veterinary Medicine, Baton Rouge, LA 70803, USA.; Institute of Cytology Russian Academy of Sciences St. Petersburg Russia.

**Keywords:** Prions, MS, MSA, Lewy Body, VLP, SMCA, Spiroplasma, Microbial Diversity

## Abstract

The extraordinary genetic flexibility of microorganisms enables their evolution into diverse forms expressing unanticipated structures and functions. Typically, they evolve in response to selective pressures of challenging niches, enabling their evolution and survival in extreme environments wherein life forms were not thought to exist. Approaching the problem of persistent neurodegenerative CNS infections as a challenging niche for pathogen evolution led to uncovering microorganisms which expand concepts of microbial diversity. These organisms are proposed as hybrid pathogens. They express two separate sets of structures and functions: viruslike properties when intracellular, and yet also reproduce as unique prokaryotes when outside the host. Their recovery opens new opportunities to comprehend the remarkable diversity of pathogens and elucidate etiologies of unresolved CNS neurodegenerative infections. Cells infected with these agents produce virus-like particles, inclusions and cytopathic effects consistent with biopsy studies of multiple sclerosis (MS), the α-synucleinopathies, and the transmissible spongiform encephalopathies (TSE) or prion diseases. The principle agents described were recovered from sheep with scrapie and are available via the Biodefense and Emerging Infections Research Resources Repository. Comparative studies with SMCA, a tick isolate inducing neurodegeneration in lab animal models, are included as supportive evidence.

## Introduction

The variety of mechanisms used by microorganisms to rapidly alter their genetics enables their successful survival and procreation. This range of genetic manipulations permits timely alterations of their structures and functions essential for survival in atypical and changing environments. Examples include colonizing challenging environments where organisms previously were not thought to exist, including near deep sea hot volcanic vents (1), kilometers within Antarctic ice (2), acidic mines (3), as stomach pathogens (4, 5) and even within clouds (6). In causing diseases, they have evolved mechanisms to infect most cells tissues and organs of animals and humans. Though the CNS is a host environment protected and perplexing for any pathogen, microorganism possess the genetic ability to evolve into agents capable of causing CNS neurodegeneration.

Success as tenacious CNS pathogens requires the means to enter the host and the CNS without alerting antibody or inflammatory immune responses, then express forms and mechanisms for slow propagation over decades, prior to initiating a pathogenic mechanism essential to exit its old host for the chance to enter a new one, completing a life cycle. Yet once mammals evolved, it is likely too late for pathogens to then develop the skill sets required for each phase of a CNS infectious cycle. The hypothesis is that an ancient infectious microorganism evolved along with the evolution of the CNS. Once this prototype neurotropic pathogen succeeded, then evolving further to expand its host range and cause diverse diseases by using different pathogenic mechanisms was straight forward. If correct, then pathogens causing different CNS infections would express similar structures, functions and cell pathologies. Until cures are available for any CNS neurodegenerative infection, this hypothesis that a similar group of atypical pathogens have evolved as causes of different diseases should be considered.

The approaches for testing perceptions of CNS pathogens include analyzing CNS biopsy data, reviewing historical studies of scrapie affected sheep as the prototype, and learning principles of pathogen isolation and identification from prior successful scientists. These tactics led to recovering unusual microorganisms from prion positive sheep with scrapie. They are also relevant to understanding the five most published theories of infectious CNS neurodegeneration which are the slow viruses, VLP, TVS, spiroplasma, and prion theories. Furthermore, these microorganisms’ structures and functions, as shown in lab cultures, are comparable to biopsy studies of patients with infectious forms of CNS neurodegeneration, and new findings of SMCA show similar structures and cell pathologies. All examples shown are available for conformation and further research.

## Material and Methods

### Host Cells

For isolations, bovine cornea endothelial (BCE) cells were used as host cells (BCE, ATCC CRL-2048). These cells were grown in Gibco DMEM medium + GlutaMAX with 4.5 g/l glucose. Additional additives to 1% of: GlutaMAX 100X; MEM NEAA; sodium pyruvate 100 mM; HEPES 1M; and to 10% FBS. To limit contamination for isolations Vancomycin HCl (MP Biomedicals 195540) were added at10 μg/ml initially, subsequently reduced to 2.5 μg/ml. After several months of passages the Vancomycin was removed. All flasks were incubated at 37° C and 5% CO2. The cells were passed using trypsin (0.25% Gibco 25200-056-100mL) to detach cells from the flask.

### Isolations

To infect cultured BCE cells, frozen eyes from four different prion positive scrapie sheep, case numbers 296, 301, 5061, and 6000 were purchased from the University of Idaho Caine Veterinary Teaching Center. The frozen eyes as received in sealed bags were thawed in a 37° C water bath. Small tissue pieces were surgically removed and added to 60% confluent BCE monolayers containing Vancomycin. The sham infected control was eye tissues from a normal sheep processed in parallel. Following months of study little difference was found between infected, uninfected and sham infected BCE despite multiple passages. However remarkable differences were found only in cells occasionally released from the monolayers. Only BCE incubated with tissues from sheep with scrapie yielded the microorganisms described. Once established as persistent infections the agents can be transferred to uninfected BCE by adding aliquots of infected tissue culture fluids.

### Cell Free Growth

After a month and up to a year, in culture, forms resembling spiroplasma bacteria were detected. Three of the four isolates were adapted to cell free growth in broth by following Tully’s procedures (36). Equal volumes (1:2) of TCF from 296, 302, and 5061 BCE cultures were mixed with SP-4 spiroplasma medium (Teknova) and initially incubated at 37° C, to eliminate possible environmental spiroplasma contaminants which cannot grow at 37° C. They were passed blind every two to three weeks by mixing (1:2) with SP-4. Once established, as detected by dark field (DF) microscopy, they were passed 1:3 every one to two weeks and incubated at 31° C for improved growth. Later they could be passed twice per week. To identify the end products of growth independent of host cells, pooled samples from weekly passages of the three examples were kept at 31°C for a month until fine granules settled to the bottom of the flask, and others could be concentrated by centrifugation at 1000 X g for ten min.

### Microscopy

Infected BCE cultures were used for comparing the fine structural features of scrapie isolates with those of suckling mouse cataract agent (SMCA). Identical microscopy methods were applied to each.

### Light microscopy

Ten μl aliquots of culture medium were deposited on glass slides, covered using one inch coverslips and sealed. The samples were examined using lenses of 63 and 100X under oil immersion. Examinations were by dark field, phase contrast and fluorescence using either DAPI or Hoechst 33342.

### Electron microscopy

For negative staining 10 μl of culture, one week to 10 days post passage, were typically placed on 100 mesh formvar-coated copper grids. After two minutes the fluid was removed by wicking. For staining 10 μl of either phosphotungstic acid 1-4% or ammonium molybdate at 2-4% were applied for one minute and removed by wicking and examined by TEM.

### TEM

For TEM of thin sections pellets of BCE or cell-free cultures were washed in PBS and fixed in 2% paraformaldehyde and 1.25% glutaraldehyde in 0.1M cacodilate buffer with 5% sucrose for one to two hours. The pellets were usually embedded in 3% agarose, cut into small cubes, washed in buffer and post fixed in 1% osmium tetroxide for 30 minutes to one hour. After rinsing in water, the samples were incubated in 2% uranyl acetate in sodium acetate buffer, pH 3.5, for two hours, washed in water, dehydrated in a graded serious of ethanol, propylene oxide and embedded in epon-araldite. Thin sections were stained with uranyl acetate and lead citrate and examined in a JEOL JEM 1011 microscope. For TEM and SEM studies of granular ‘hedgehog’ forms 250 to 300 mls 6-8 weeks post passage were concentrated by centrifugation as 3700 RFC for 90 minutes prior to processing.

### SEM

For SEM of cell-free cultures one to two weeks old five ml of culture fluid were filtered through 0.22 μm pore size polycarbonate filters. The filters were gently rinsed in PBS and fixed using standard SEM methods. Following dehydration in ethanol and critical point processing using CO_2_ the filters were mounted on stubs and examined by SEM.

### Assay for α-synuclein and prion aggregations

Cultures of BCE alone, infected with SMCA and separately with the scrapie isolates were concentrated, fixed in formalin. They were paraffin embedded, sectioned and attached to glass slides by Angelina Demming of the National Hansen’s Disease Program at the LSU SVM. The slides, cleared of paraffin, were provided to histologist Ji-Ming Feng (CBS Dept. LSU SVM) for α-synuclein and prion assays. To confirm the α-synuclein aggregations were not a quirk of the BCE cell line, aliquots of BCE persistently infected with either scrapie isolates, SMCA or uninfected controls were separately added to N2a cultures. After weeks of incubation whole cells were formalin fixed and also provided to Ji-Ming Feng for the respective prion and α-synuclein amyloid assays (7, 8, and 9). Of interest, corneal endothelia cells express α-synuclein (10).

### SMCA Lewy body type inclusions

These inclusions were found by mimicking a persistent infection. Persistently infected monolayers were maintained without passage by replacing the TCF every 5-7 days for months. Over time most cells became granular, reduced in size and piled together forming rings of cells. Cells central to the rings were comparatively clear and flat. From these central cells evidence of the double membrane Lewy body was found. They formed within large open nuclei, rather than in the cytoplasm.

## PCR of the spiroplasma Hsp60 sequence

Next, 1.8 ml of cell-free culture were concentrated by centrifugation at 14000 G for 30 minutes. DNA was isolated by Roche High Pure Template Preparations Kit (Roche Applied Science).

The PCR probes to amplify spiroplasma Hsp60 were designed and applied by Lidiya Dubytska, PBS, using the Hsp60 sequence incorporated into US Patent 7,888,039 B2: 2011 “Assay for Transmissible Spongiform Encephalopathies” by Frank O. Bastian. The primers used were:

Hsp60 F T gat att gct ggg gac ggt act ac

Hsp60 R at cac ttc ttc aac agc tgc ctt agt cga G

## 3. Results

(see attachment)

## 4. Discussion

Today, prions “proteinaceous infectious particles” are widely accepted as the infectious cause of TSE, with scrapie affecting sheep (the prototype) and CJD (the human equivalent). Yet doubts remain that a protein could evolve into a pathogen, an infectious cause of disease. Organisms with nucleic acids encode proteins. If a protein mutated to become lethal it would kill its own origin. Proteins can alter protein conformations but without nucleic acids their total numbers (defined by amino acid sequence) cannot increase. Natural pathogen transmission is inefficient requiring large numbers for few successes, improbable for proteins alone. Pathogenic prions can form by genetic errors (11), cross species barriers (12), and are not cured by immune responses (13), so why are the TSE diseases rare? Prions are mammalian proteins capable of folding into conforms that aggregate, and by binding to normal conforms also induce their misfolding. Collectively they cause the disease pathology. Whatever initiates the autocatalytic misfolding process leads to the prion diseases. Many microorganisms modify host proteins as pathogenic mechanisms, why should the naturally transmitted TSE cases be different?

At the Rocky Mountain Laboratories (RML), NIH pathologist William Hadlow studied naturally acquired scrapie in sheep. He connected scrapie pathology with that of the human disease Kuru (14). At the RML he taught that prions alone could not account for all the TSE pathology. In natural scrapie the incubation period from exposure to disease is often years, time enough for a pathogen to alter the normal prion conform as its pathogenic mechanism, causing scrapie clinical signs and spongiform pathology. Since the prion mechanism is autocatalytic (15, 16), it is transferable by brain injections, causing disease without being a naturally transmissible infectious agent. Perhaps the difference in interpreting data between Hadlow and Prusiner is that each studied a different model. Prusiner chose the lab animal hamster scrapie model, maintained by brain injections, whereas Hadlow studied natural scrapie in sheep. Hadlow’s insights stimulated searching for pathogens overlooked in scrapie affected sheep.

### 4.1 Recovery and Propagation of Agents from Sheep with Scrapie and Comparisons with SMCA

For primary isolations eye tissues were chosen because eyes are affected in scrapie and eyes evolved as brain extensions. They are potential routes for neurotropic pathogens to enter or exit the host. A BCE cell line derived from eye corneal endothelial cells was used as this cell type does not seem to support prions. Without available diagnostic reagents, primary isolations were difficult. Studying potentially infected monolayers for signs of pathology failed to detect differences between normal monolayers, those incubated with eye tissue from sheep with scrapie or sham infected with tissues from a normal sheep eye, even after multiple serial passages and months in culture as studied by LM and TEM. In cell cultures some cells are routinely released from the monolayer and spaces left filled by dividing adjacent cells. Examining released cells from each monolayer began to reveal differences. At first morphology indicated cell death in infected monolayers was by apoptosis, but detailed TEM examination also revealed products of a pathogen (Fig. 1, a-u). Subsequent studies with SMCA showed that its infectious process is similar to agents recovered from scrapie affected sheep. Studies with SMCA indicate apoptosis is a mechanism to obtain host cell constituents for its use. In thin sections, the surface blebs of infected cells, a characteristic of apoptosis, reveal granular fibrillary material of the pathogen including VLP (Fig. 4, c-f). Fluorescein labeled antibodies, raised against SMCA grown cell free, were provided for these studies. When applied to SMCA infected cells they showed positive reactions but typically as large inclusions with little evidence of spiroplasma (Fig. 4, o-q).

**Figure 1.**
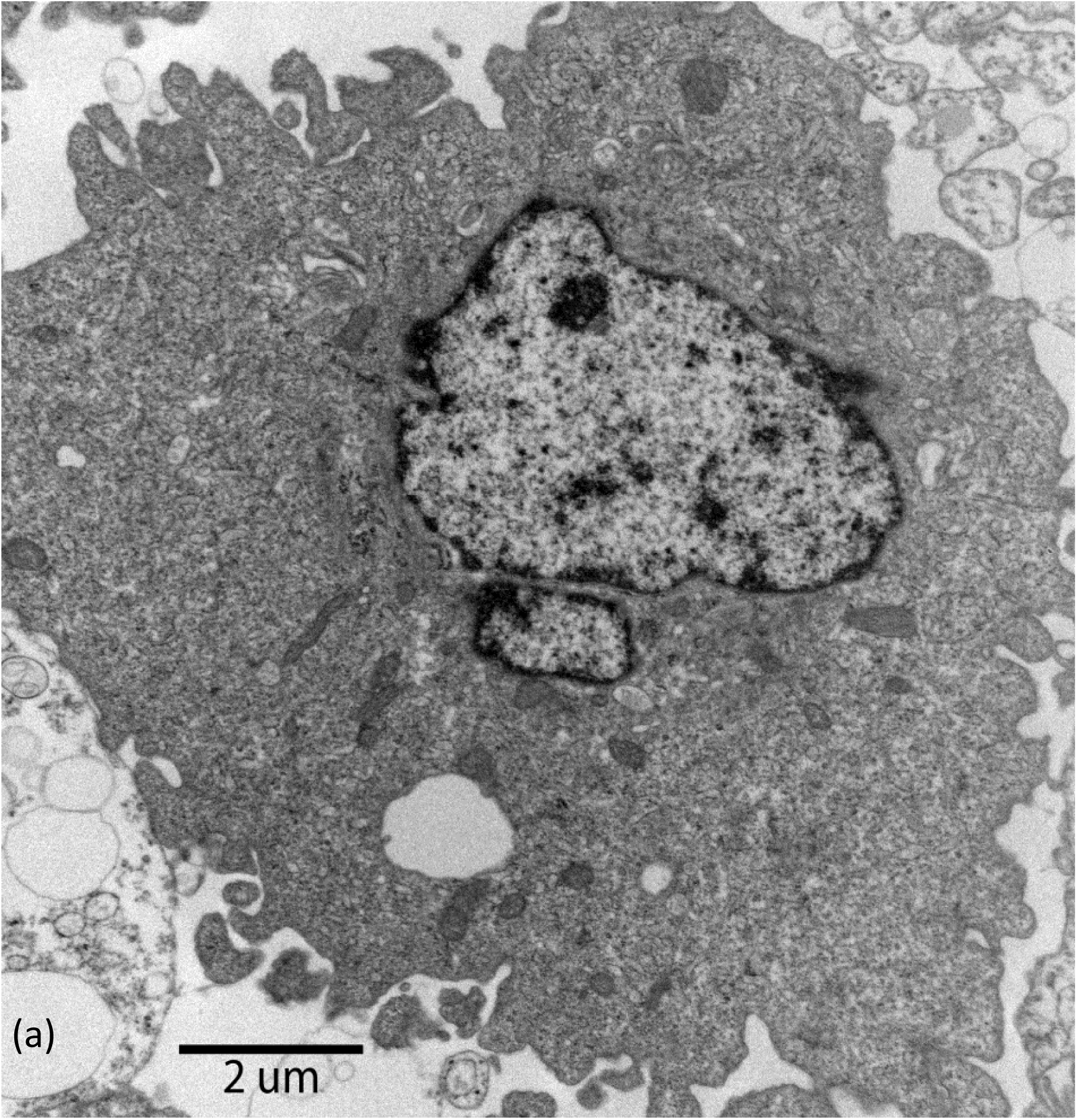
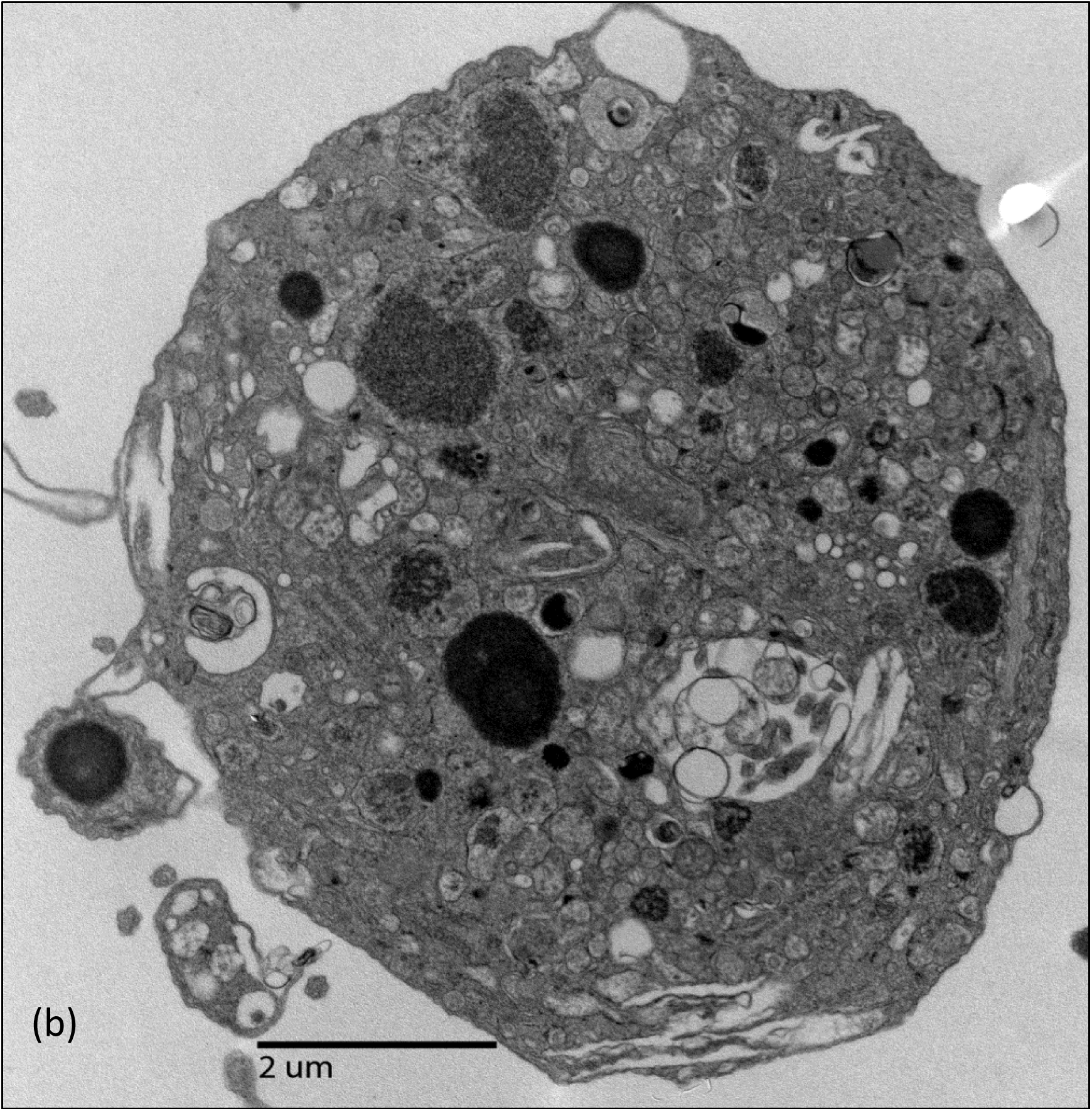
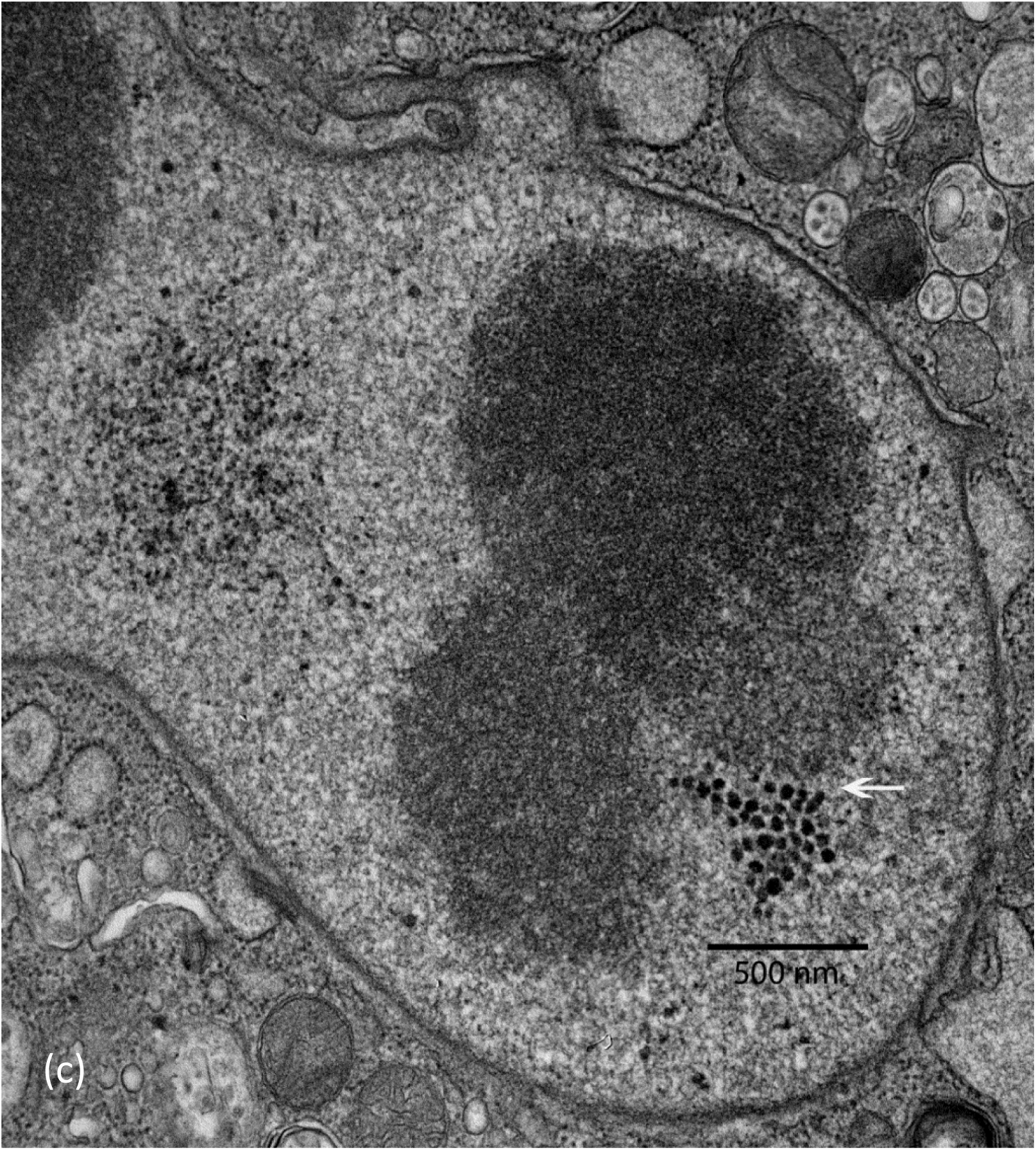
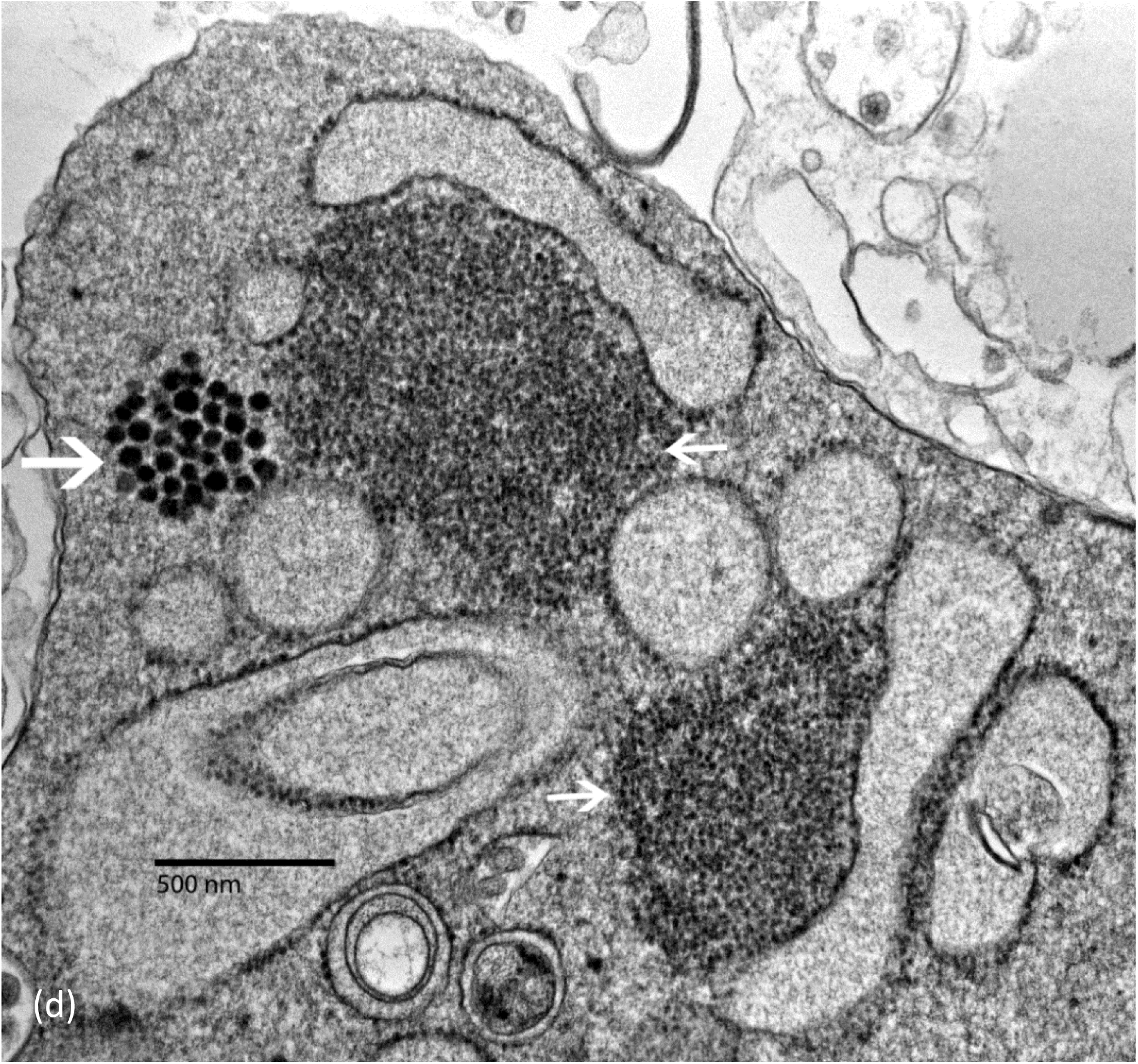
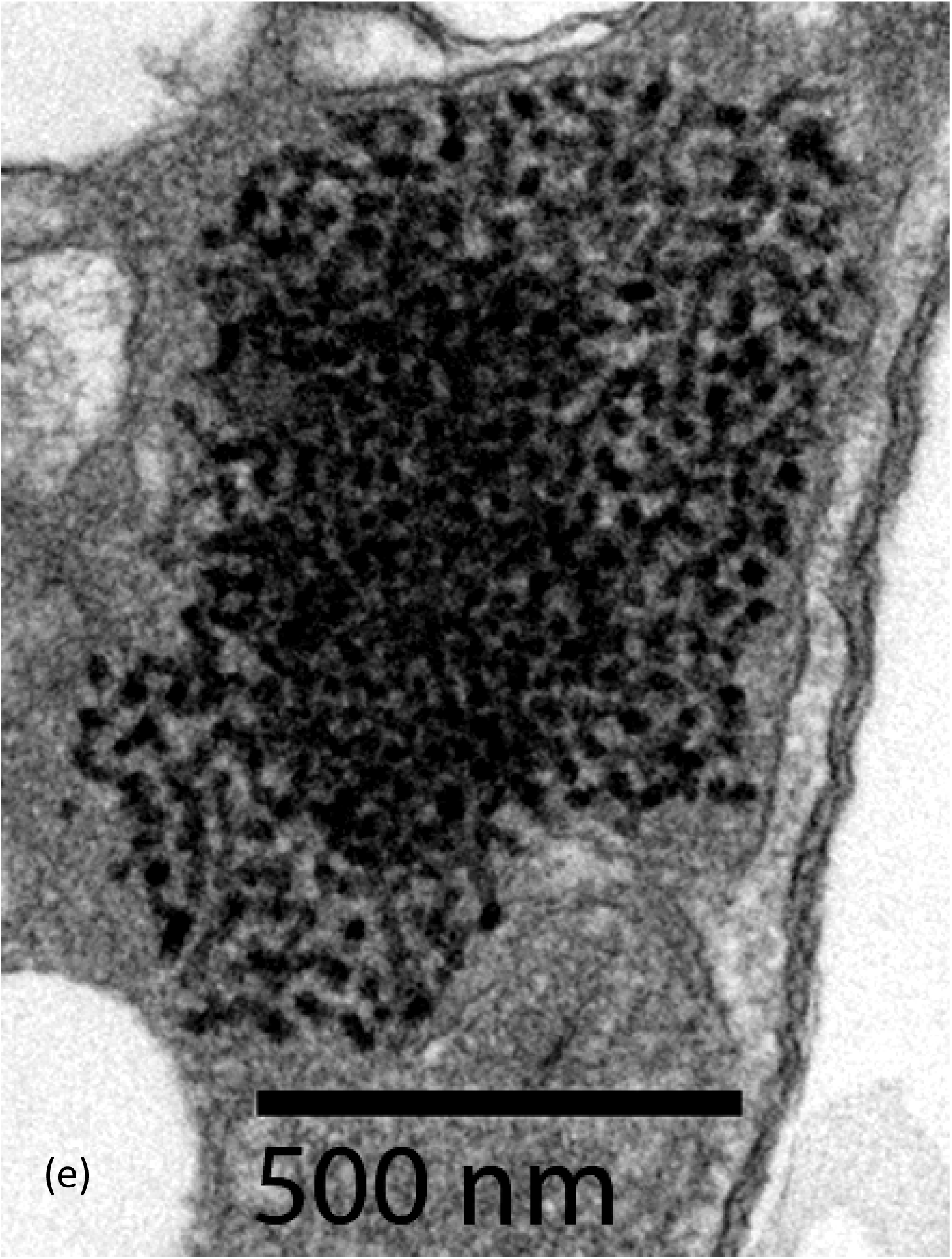
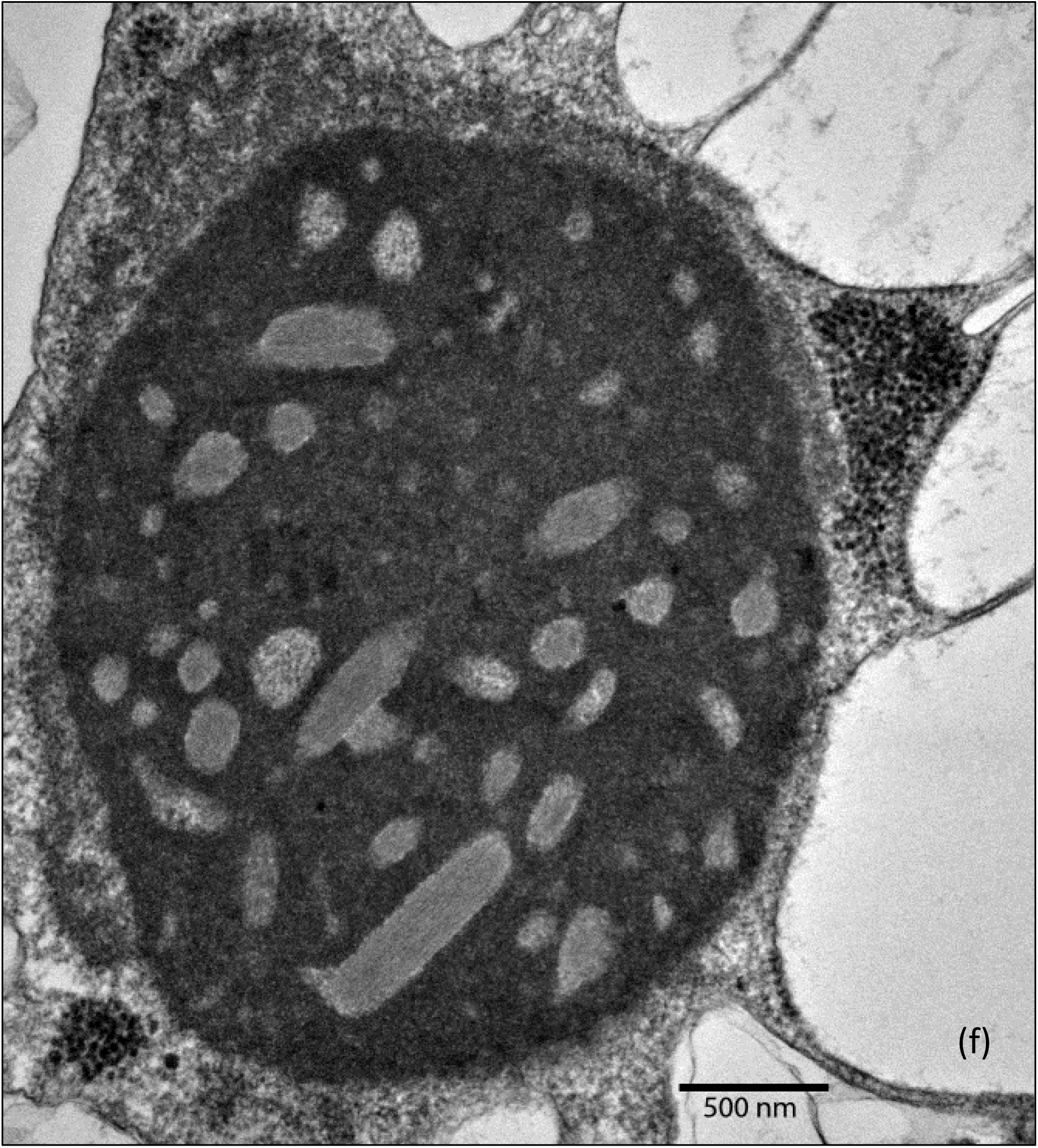
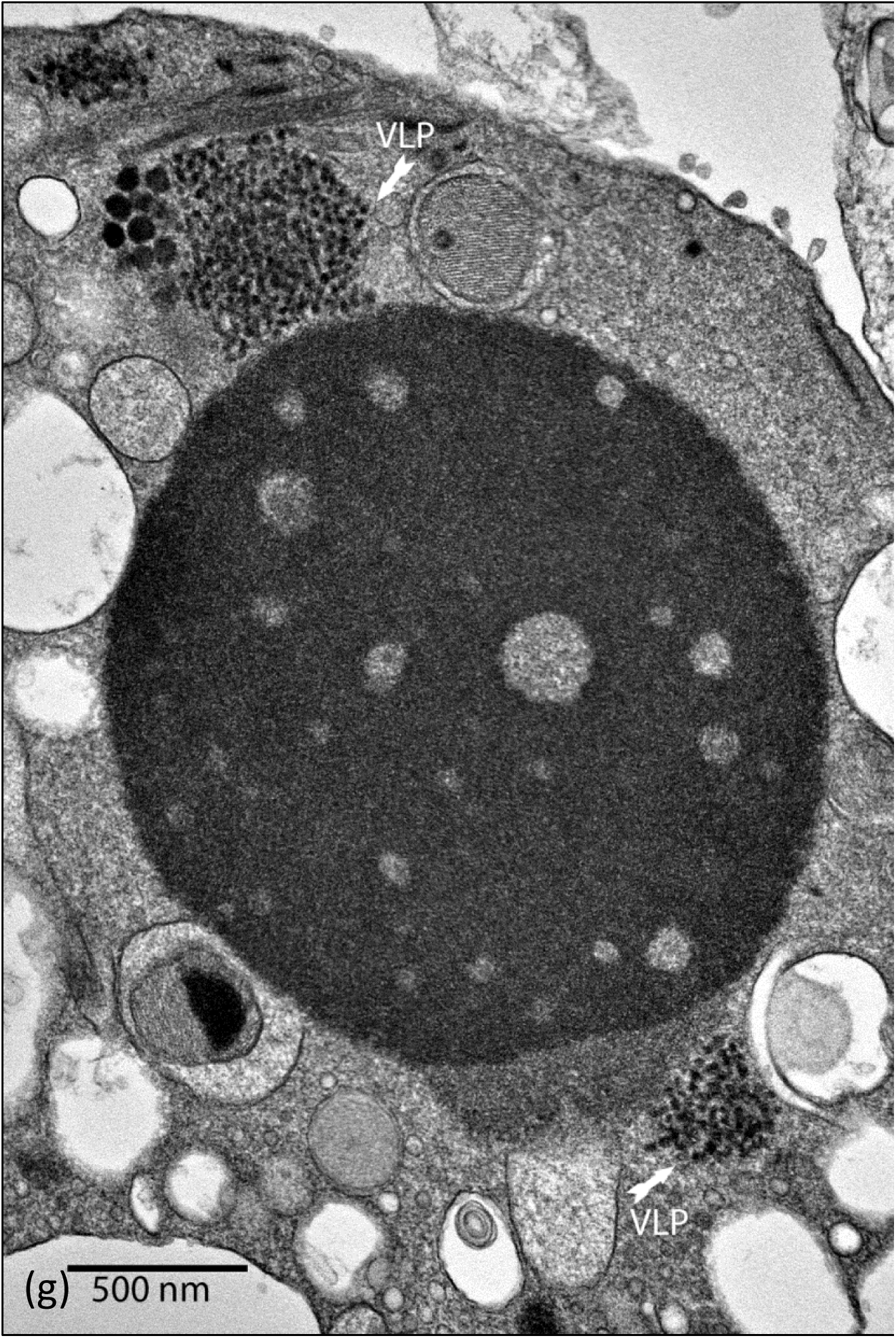
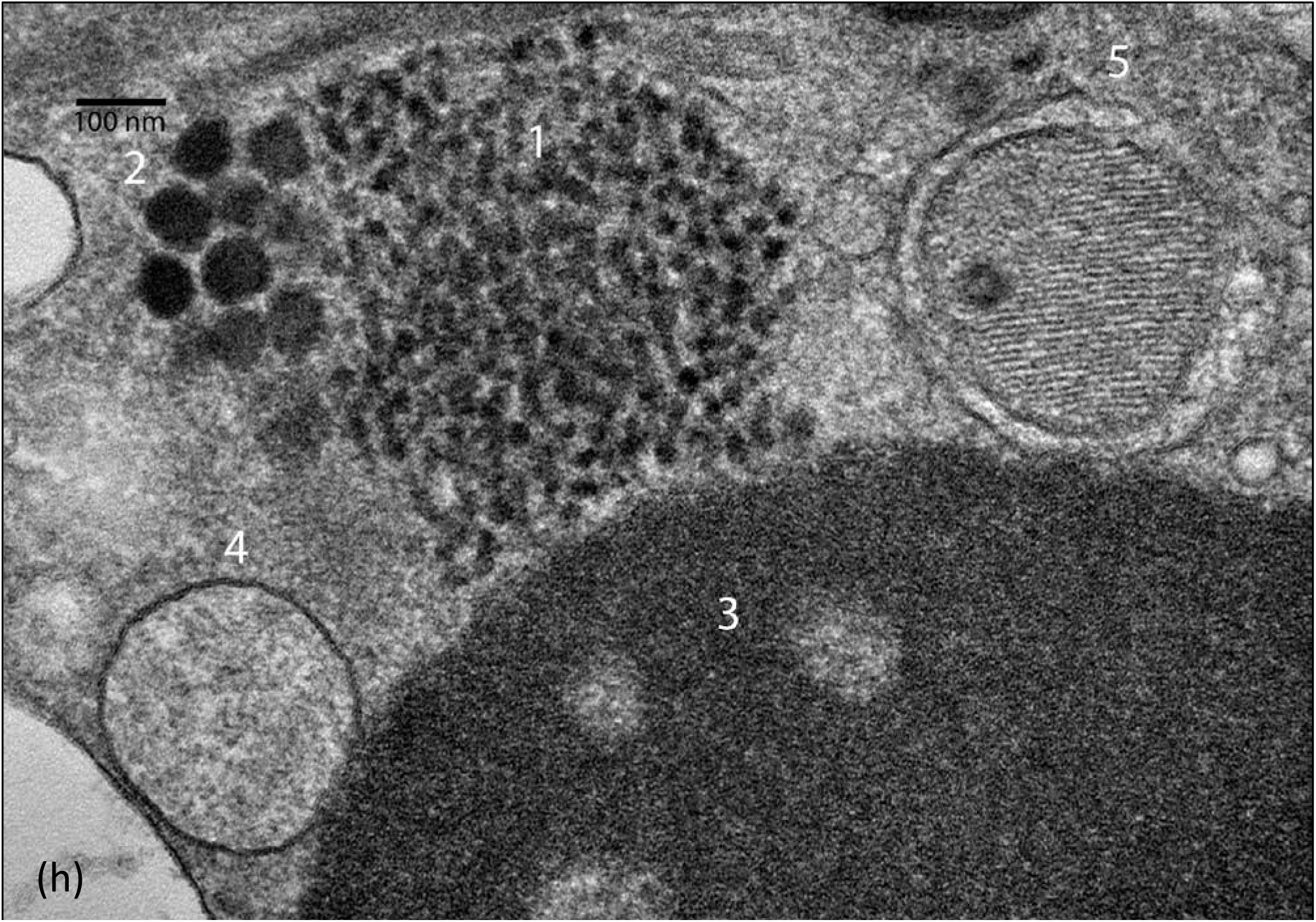
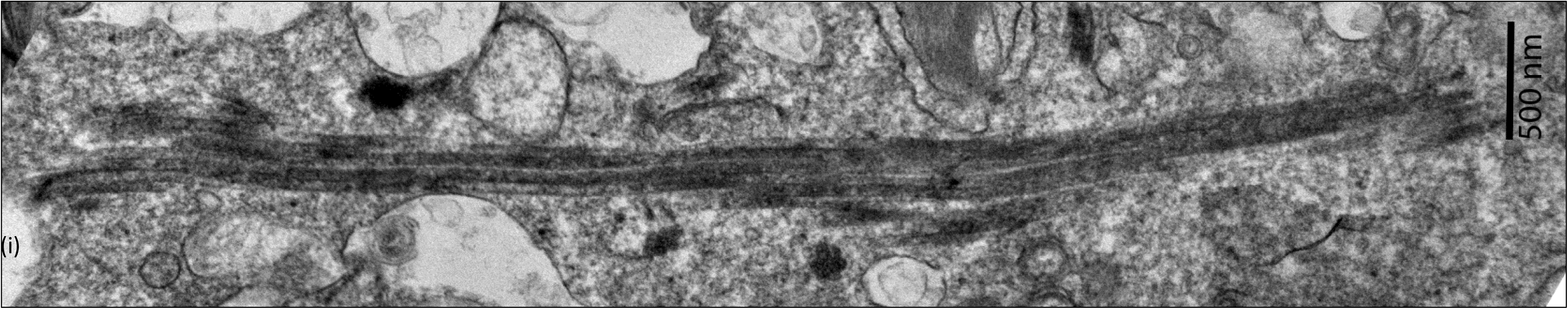
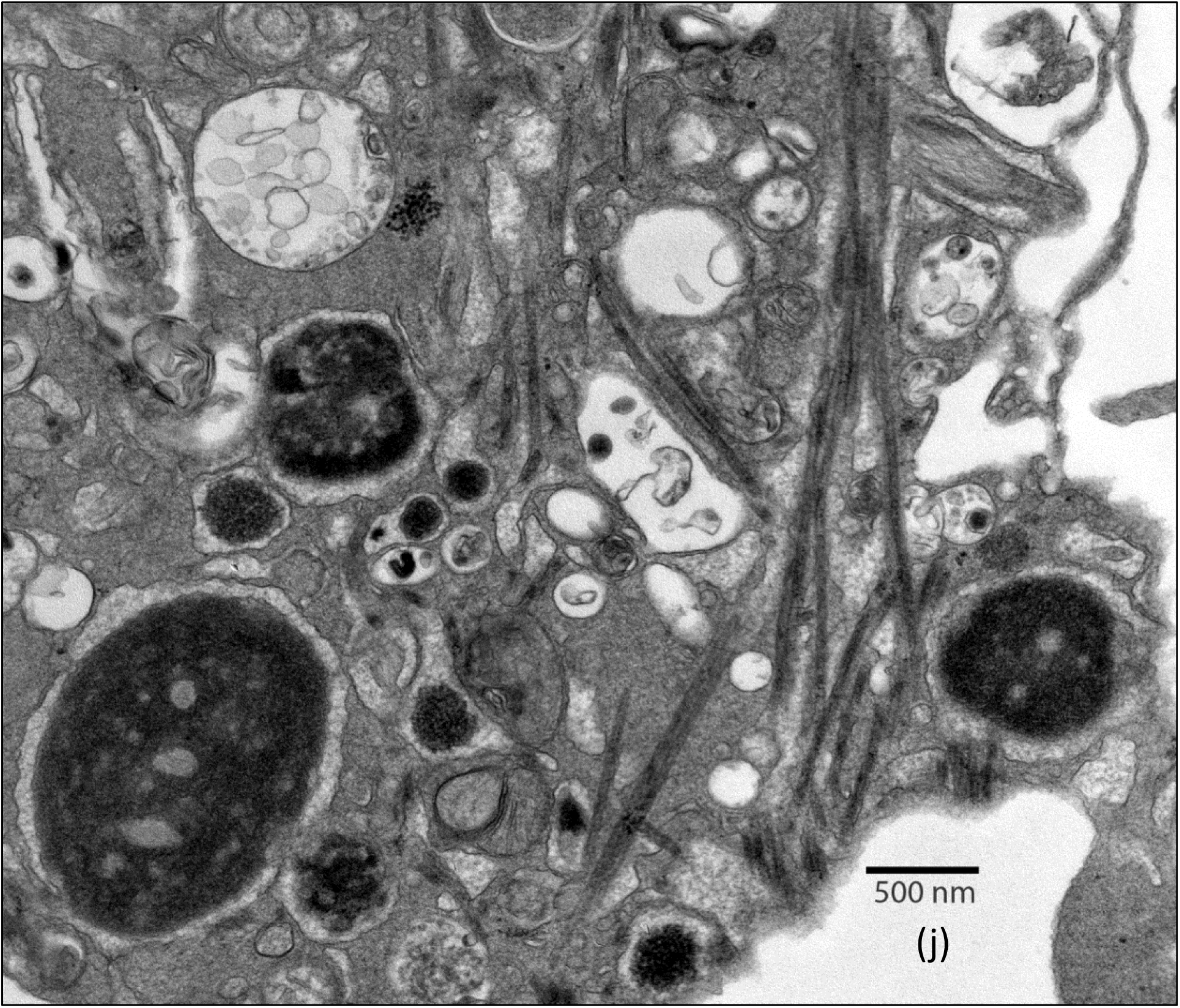
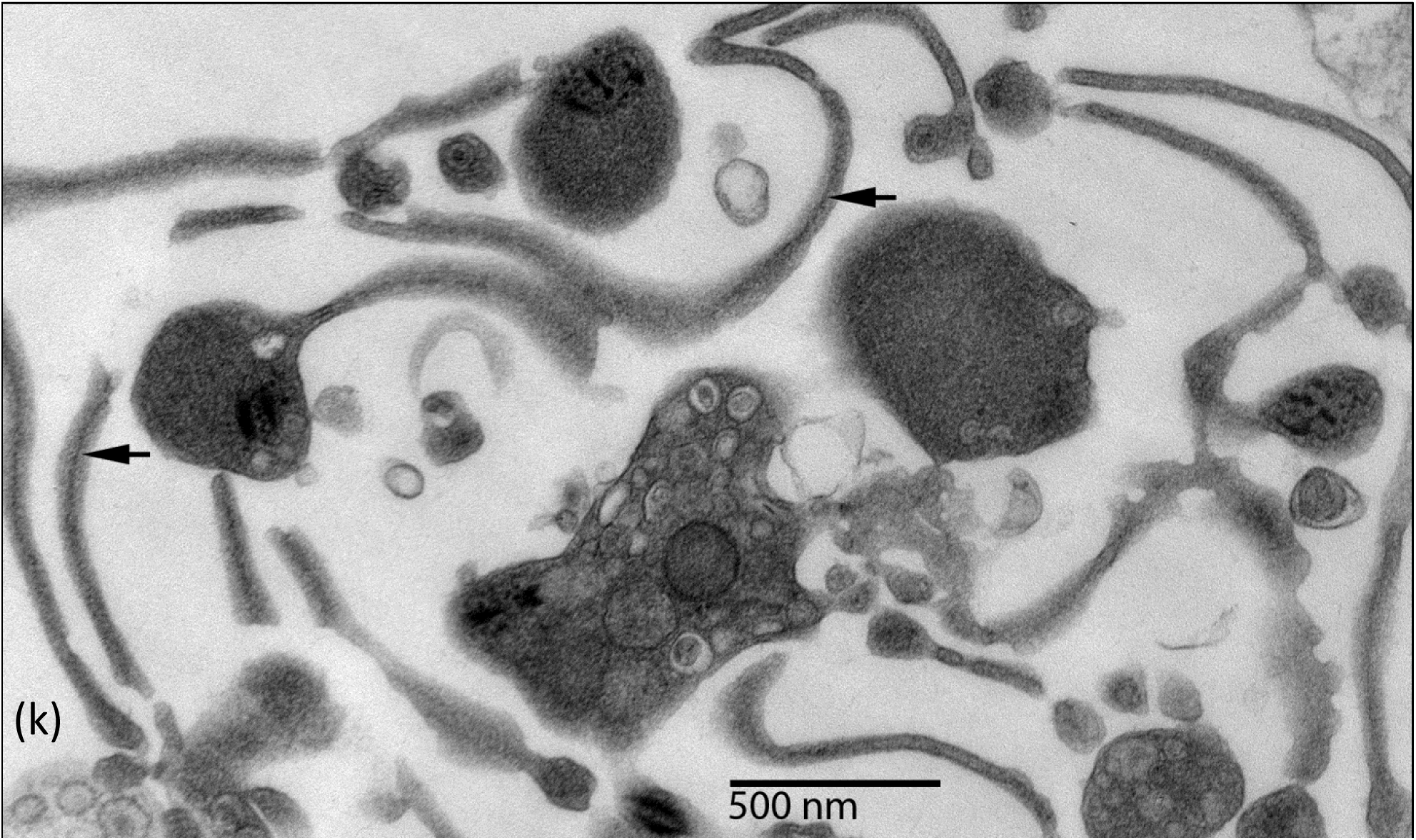
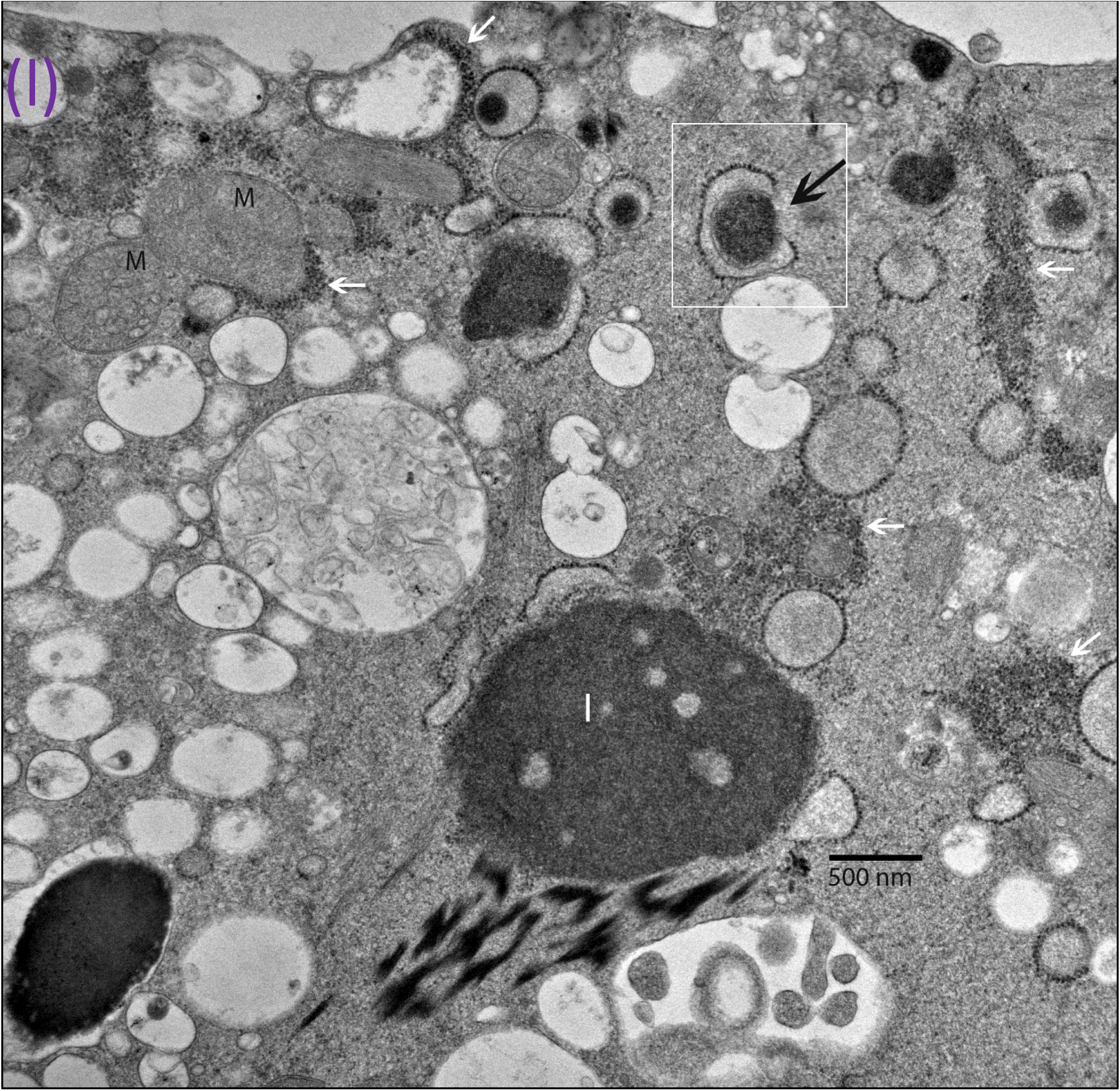
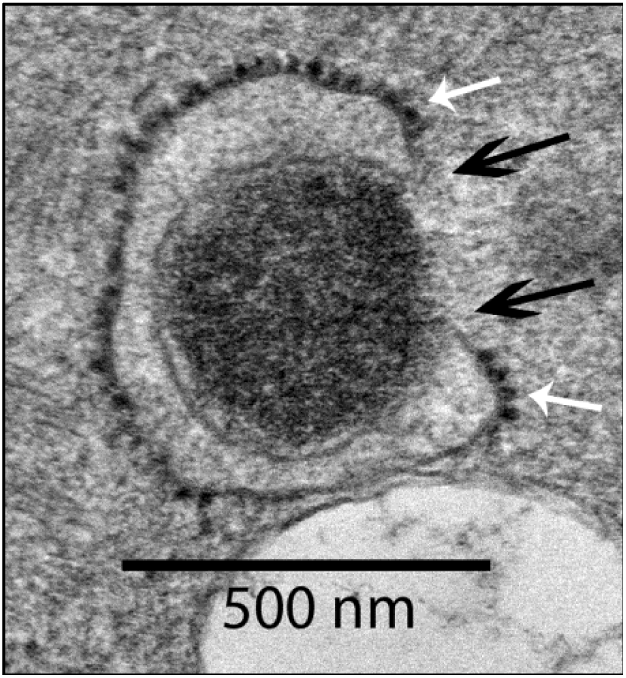
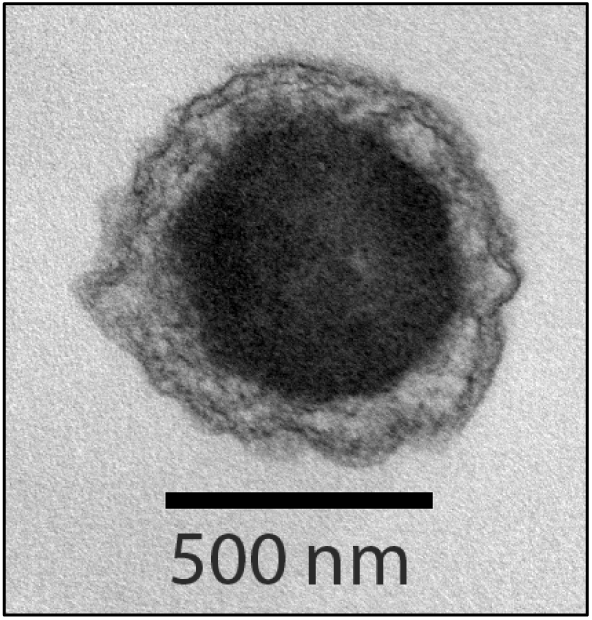
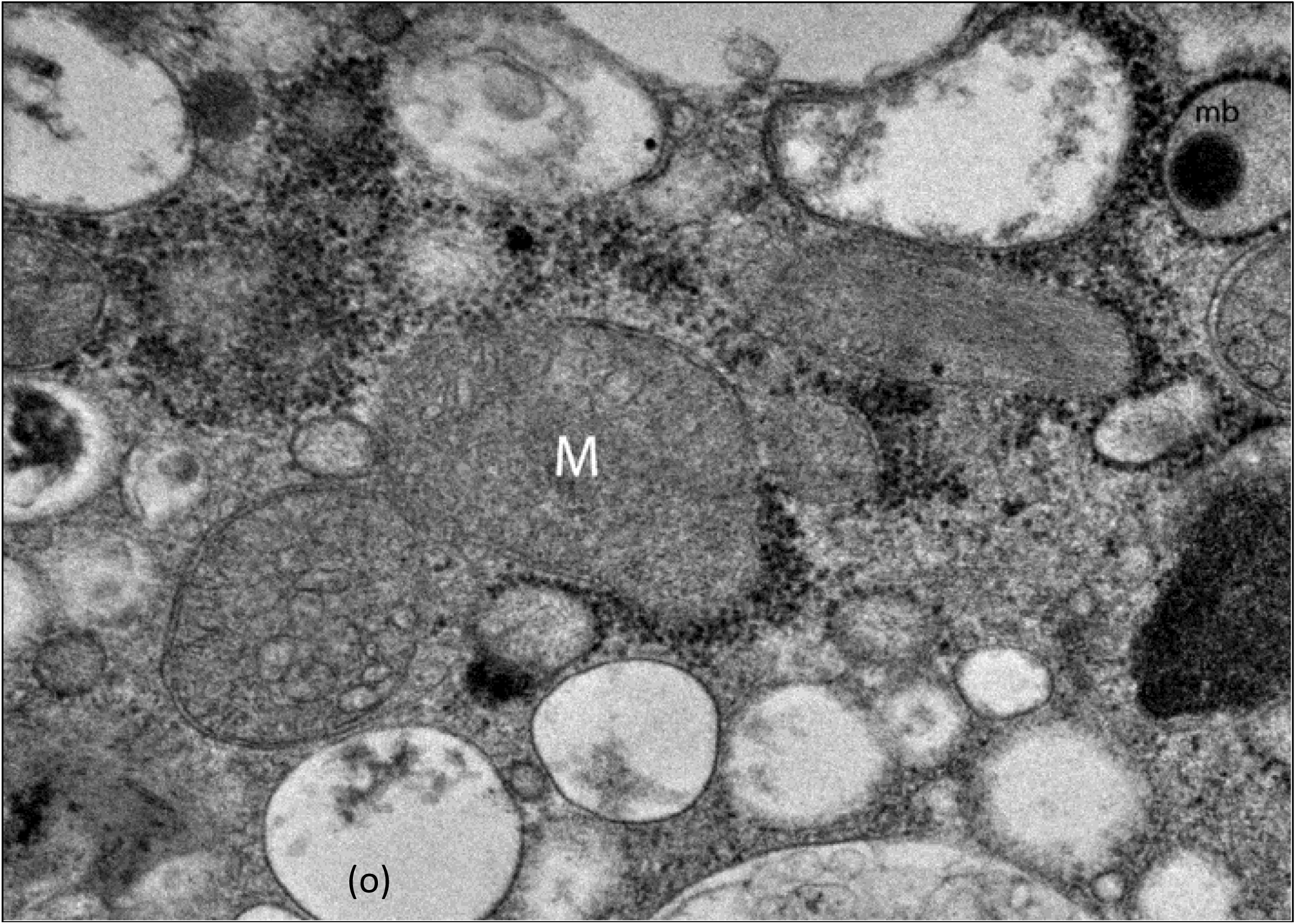
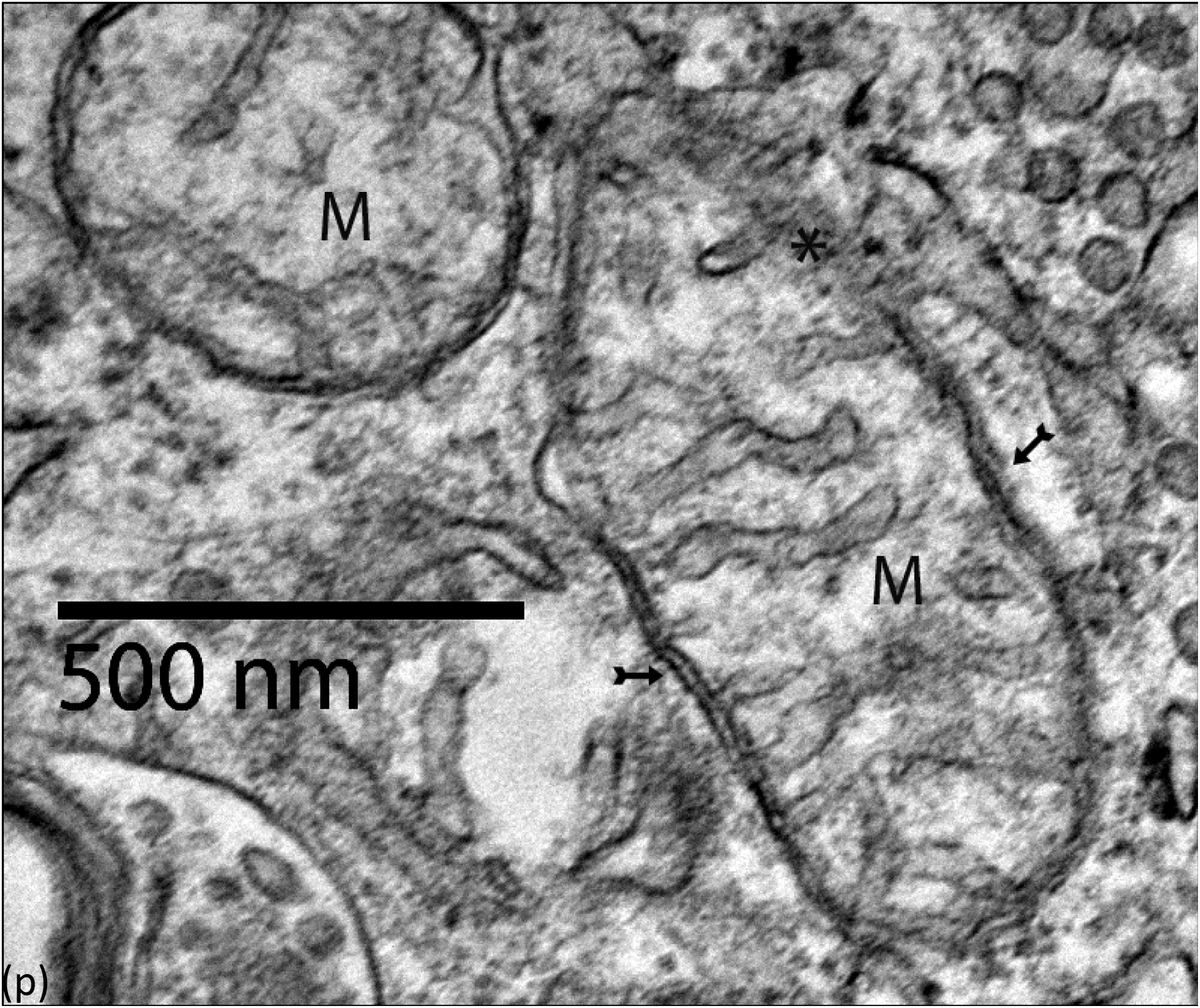
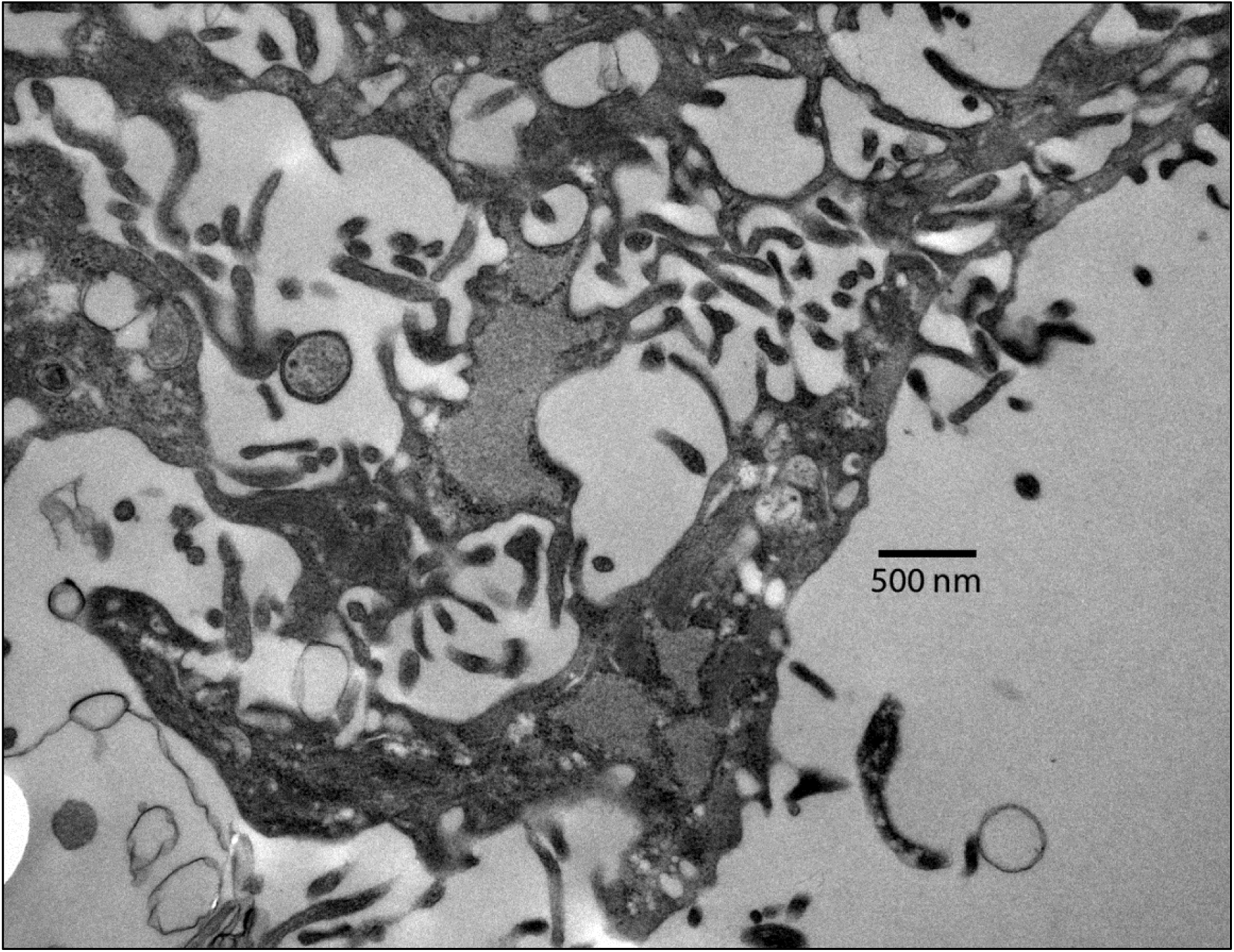
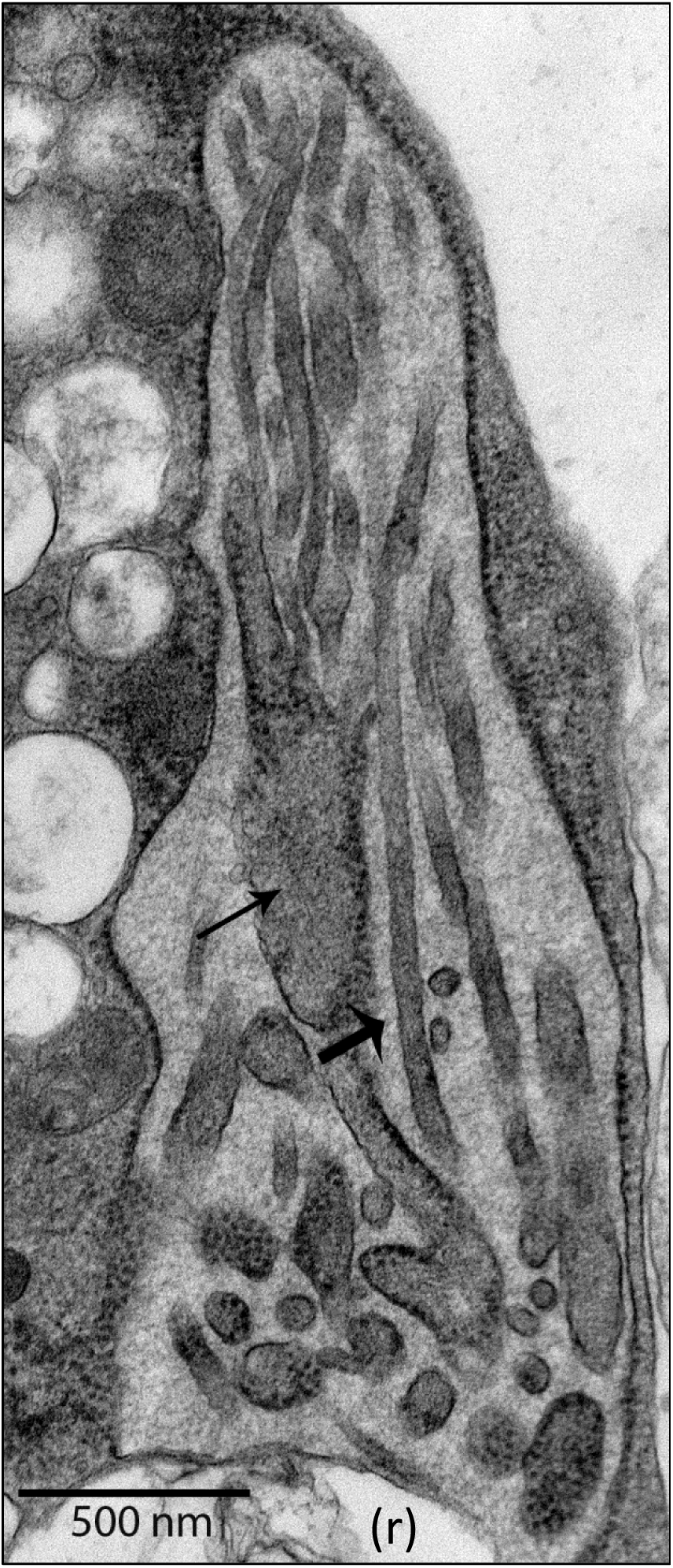
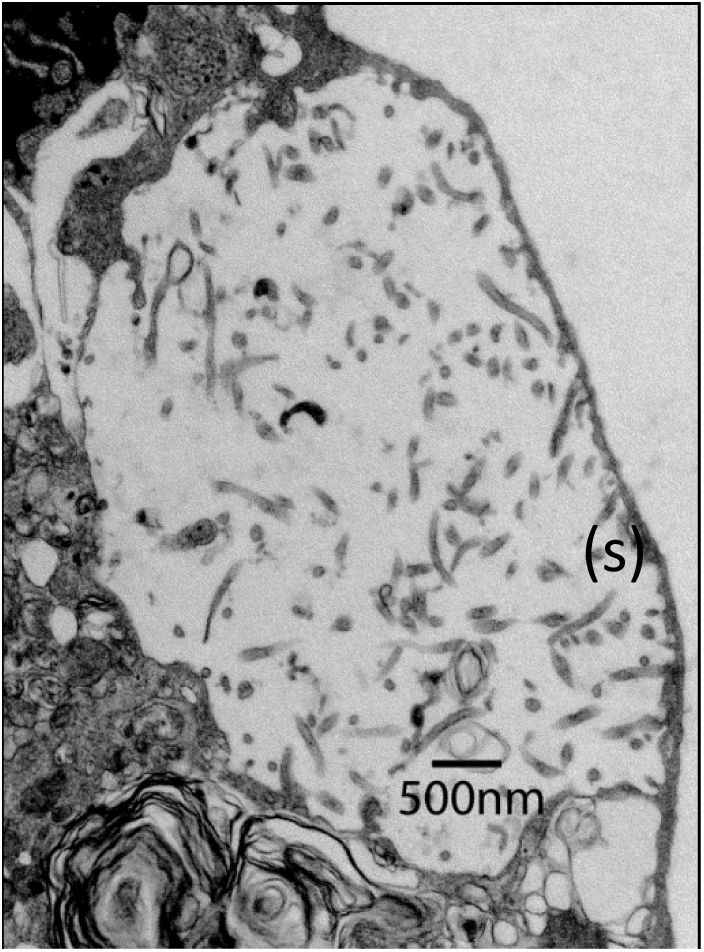
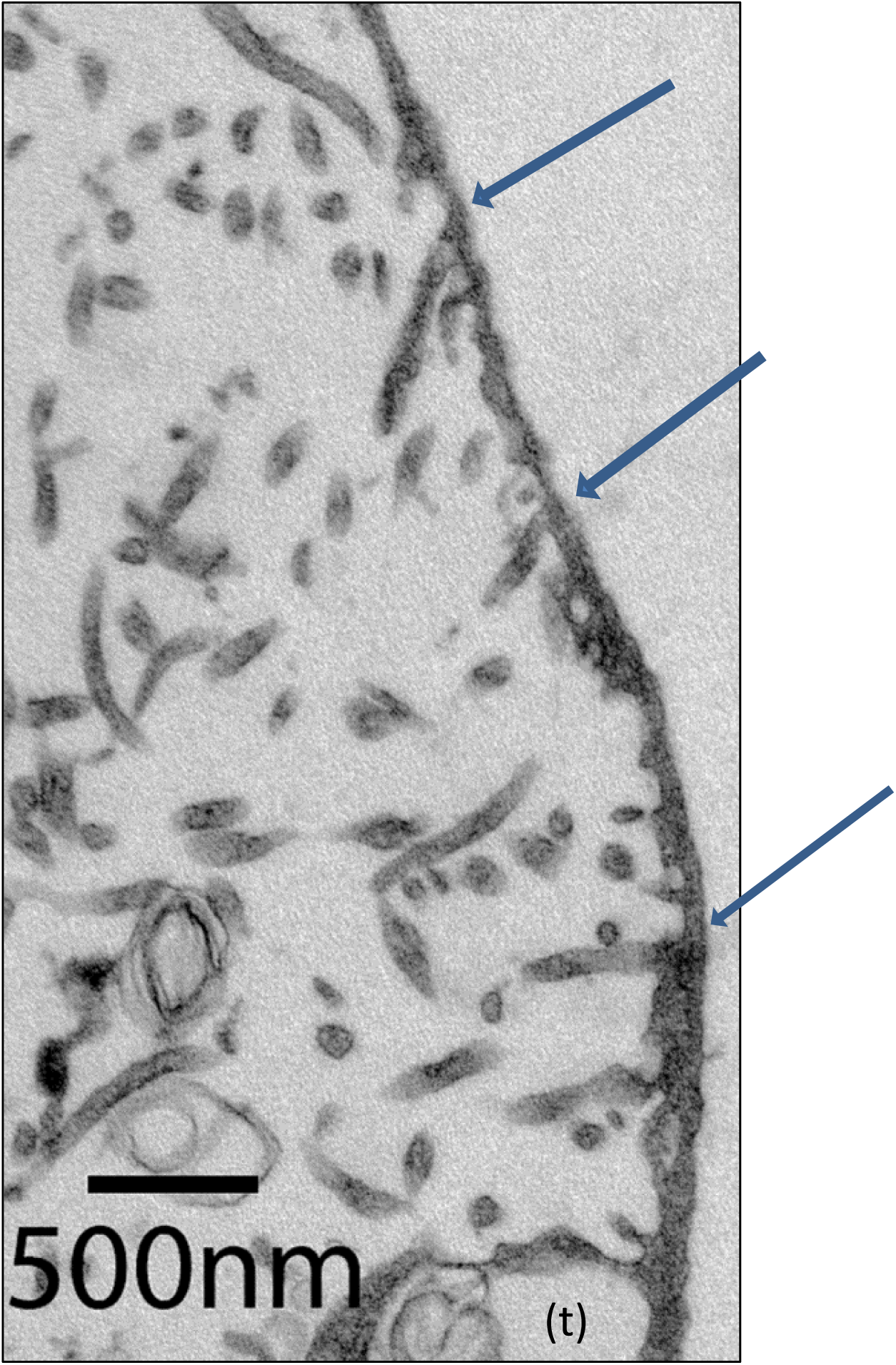
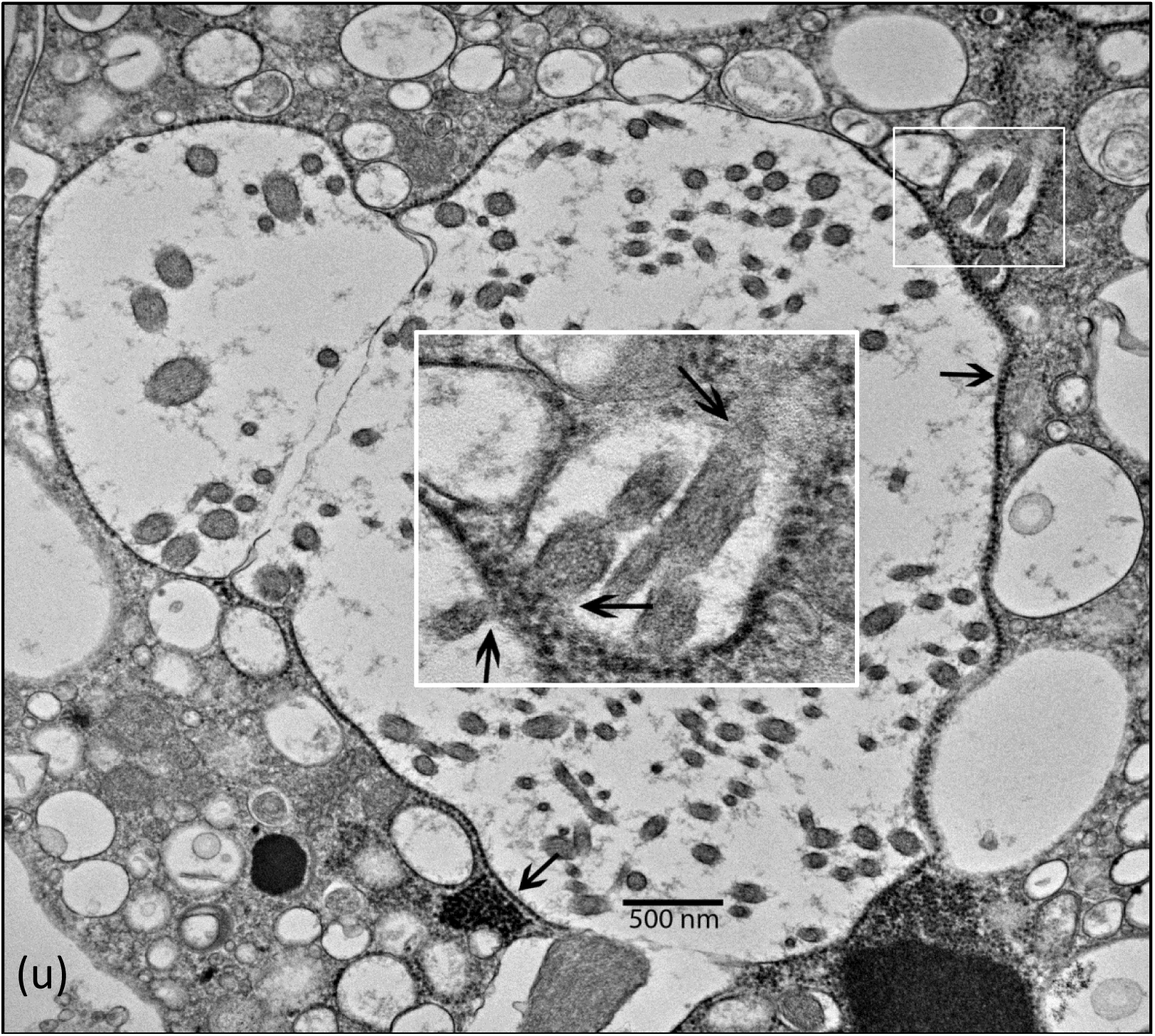
The Effects of Infecting Cells with Agents Recovered from Sheep with Scrapie. (a) Uninfected bovine corneal endothelial cell (BCE) (b) Infection of BCE by the agent from sheep with scrapie destroys the organelles of the host cell, replacing them with products of the pathogen. (c) and (d) After VLP form in the nucleus (arrow), the nuclear membrane breaks down releasing nuclear contents into the cytoplasm. The VLP are condensations along granular fibrillary strands as shown in high magnification (e); because their formation is by aggregations along the strands they can form different sizes of VLP as indicated by white arrows above. (e) The pathogen’s ‘VLP’ are clusters along granular fibrillary strands, likely DNA; and they are continuous with the strands rather than individual particles. (f) Most common inclusion detected. They are also continuous with VLP as detected in peripheral areas. (g) and (h) Individual cells will vary with the types of products produced. Most are continuous with VLP. White arrows (g) show small VLP. (h) #1 small VLP, #2 large VLP, #3 large inclusion continuous with VLP, #4 amorphous inclusion, #5 inclusion of parallel arrays. (i) And (J) tubular structures filled with granular fibrillary material. (k) tubular vesicular structures filled with granular fibrillary material were found at peripheral areas of infected cells. (l) Infecting cells with isolates from sheep with scrapie results in producing products of the pathogen and changes the entire cell. White arrows show VLP. The white I is an inclusion with continuity with VLP and other material. The white box and dark arrow show packaging of a membrane bound form. (m) Enlargement from Fig. (i). Assembly of a membrane bound structure. Arrows indicate assembly process. (n) When mature they resemble minimal bodies of obligate intracellular bacteria, though they are produced using virus-like mechanisms. (o) Enlargement of mitochondria degeneration from Fig. (l). They break down in association with VLP and granular fibrillary material of the pathogen as shown in (p). The letters M are mitochondria. Dark arrows seen at high magnification show chains of granules along the opened mitochondrial membrane. (q) The granullar fibrillary material is also detected within cytoplasmic extensions about the size of microvilli. They appear as extensions of the host cell membrane. (r) Similar long thin forms can also form within vacuoles. (s) Inclusions contain forms easily mistaken as spiroplasma, but at higher magnification (t) they form *de novo* in the infected cell cytoplasm and mature by budding through host membranes (arrows), analogous to viral processes. They are likely transitional stages between intracellular and extracellular functions. (u) Formation of spiroplasma -like forms by budding through a membrane of the host cell. These forms are not the same as those that propagate cell free.

Infecting cells with either scrapie isolates or SMCA reveals similar structures and cell pathologies, distinct from known pathogens. Hardly anything is known about them. First, we should determine if their structures and cell pathologies are relevant to the literature on CNS neurodegeneration. A catalogue of figures is first presented followed by discussing their relevance to the literature and theories of causes.

The process of infection replaces host cell constituents with those of the pathogen (Fig. 1, b). The most significant feature is the formation of virus-like particles (VLP), although sometimes they are of two different sizes (Fig. 1, c, d, and h). They appear first in the nucleus, then as the nuclear membrane breaks down they are apparent in the cytoplasm (Fig. 1, c-h). Within the cytoplasm VLP appear as clusters along granular fibrillary strands (Fig. 1, e) and are continuous with complex virus-like inclusions (Fig. 1, e-h). Note the near complete destruction of host-cell organelles and formation of large inclusions, some surrounded by a membrane (Fig. 1, h and i). Surprisingly, shapes resembling spiroplasma emerged over time (Fig. 1, q-u) but they develop analogous to viruses, receiving their limiting membrane by budding through host-cell membranes, sometimes accumulating within inclusions (Fig. 1, r, s, t and u). The spiral shapes (formed by host cell processes) may be a transition stage between the pathogen’s intracellular processes and those for cell free propagation. It is noteworthy that the morphology of the scrapie isolates and SMCA that propagate cell free (Fig. 2, a, and 5, a) are similar to each other yet distinct from those of described bacteria.

**Figure 2.**
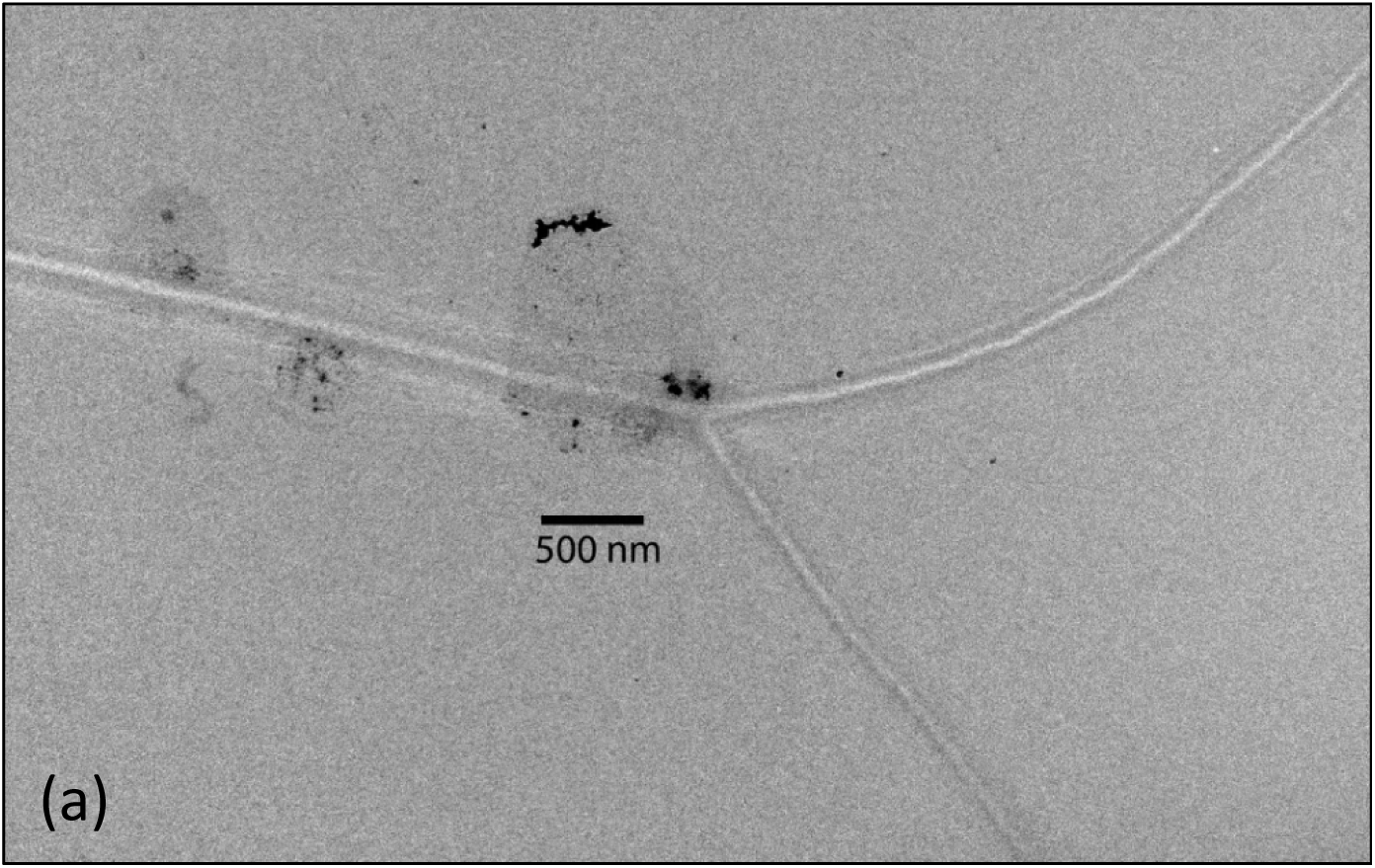
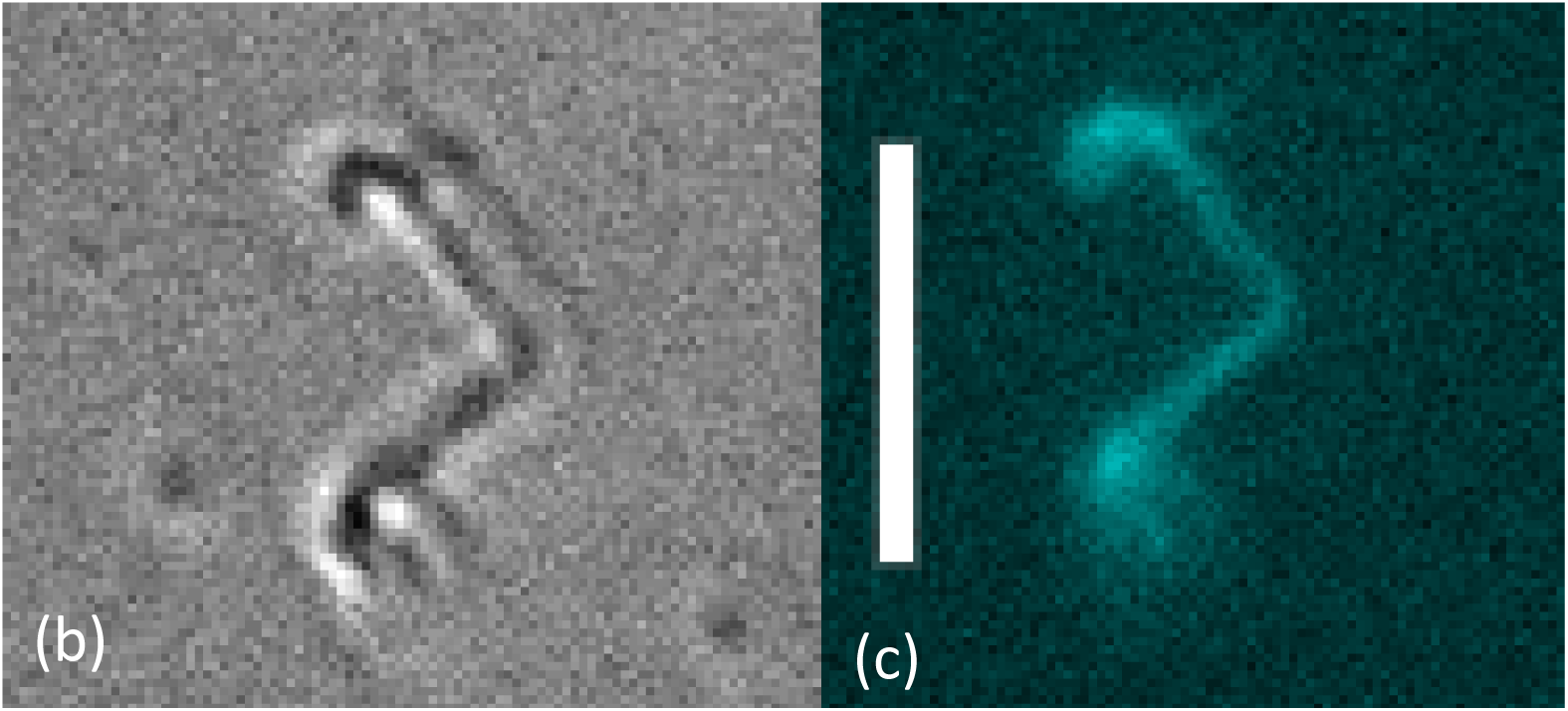
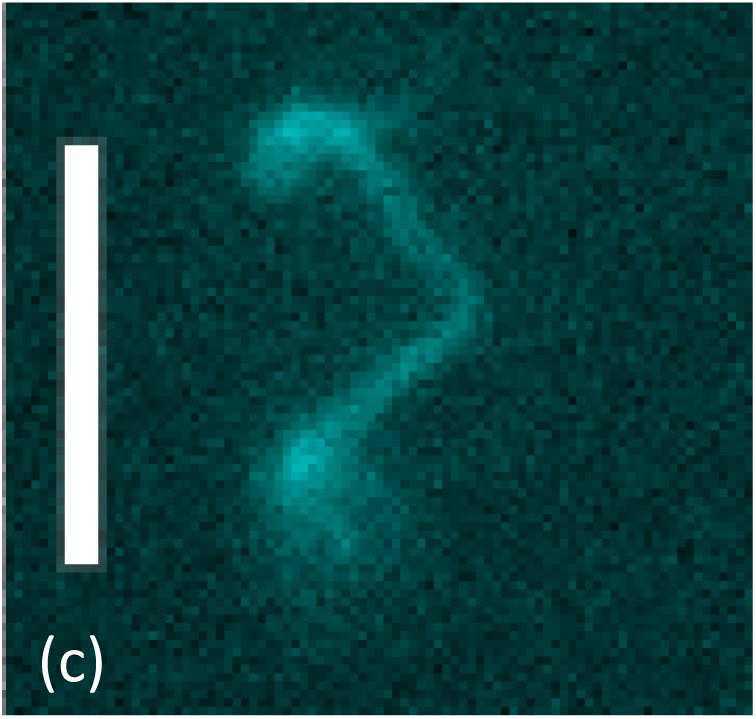
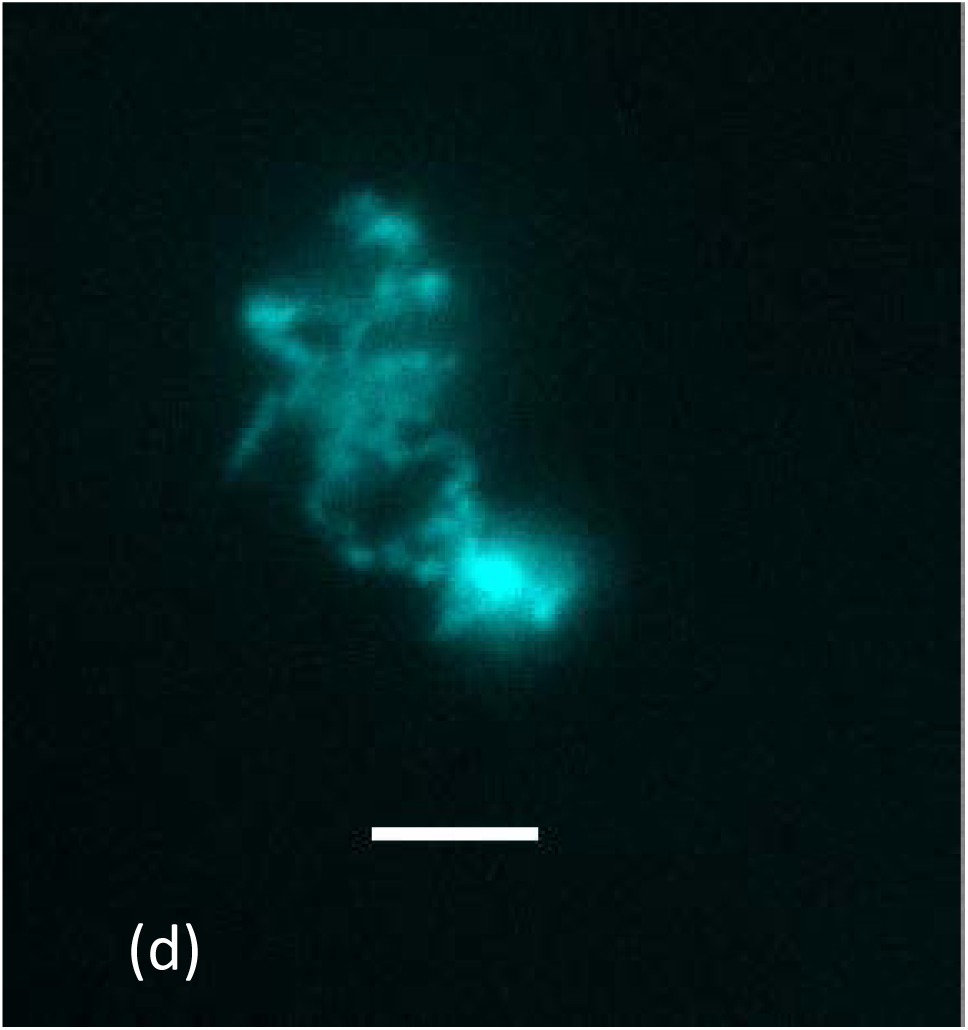
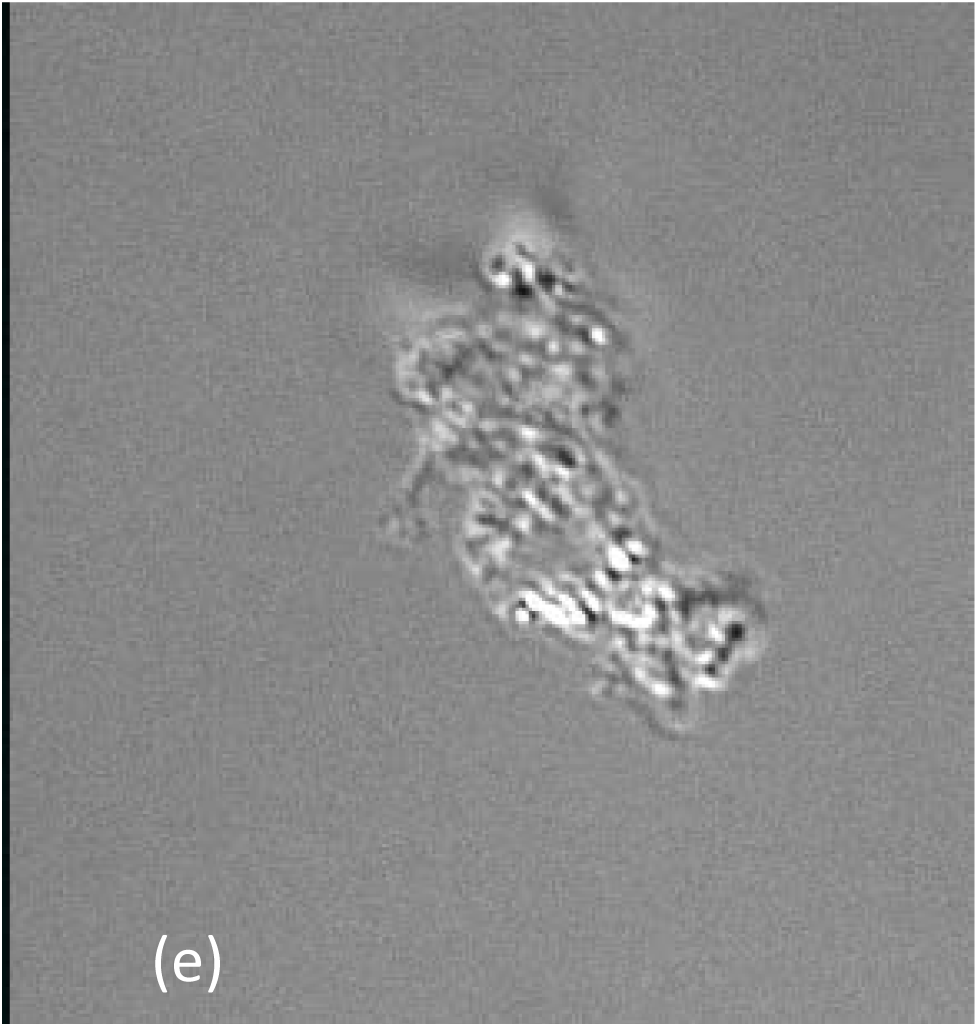
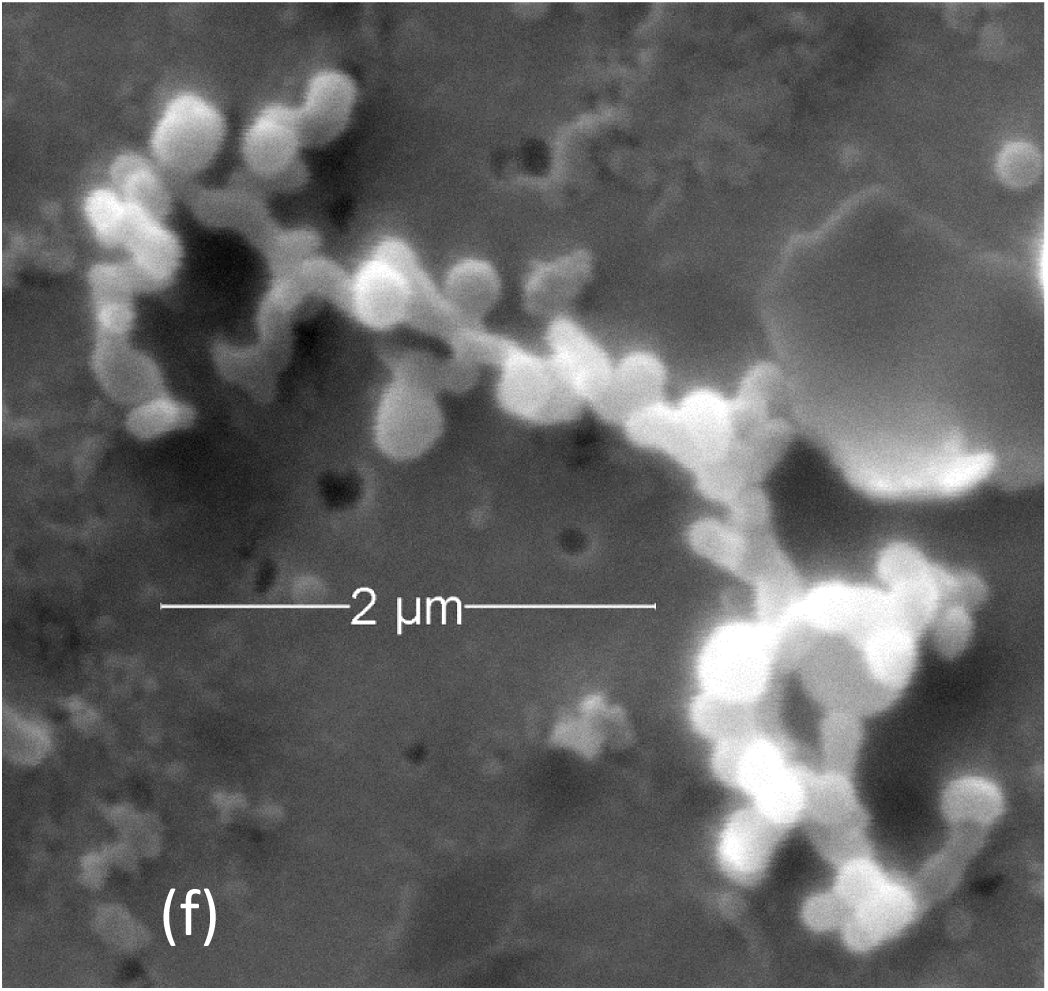
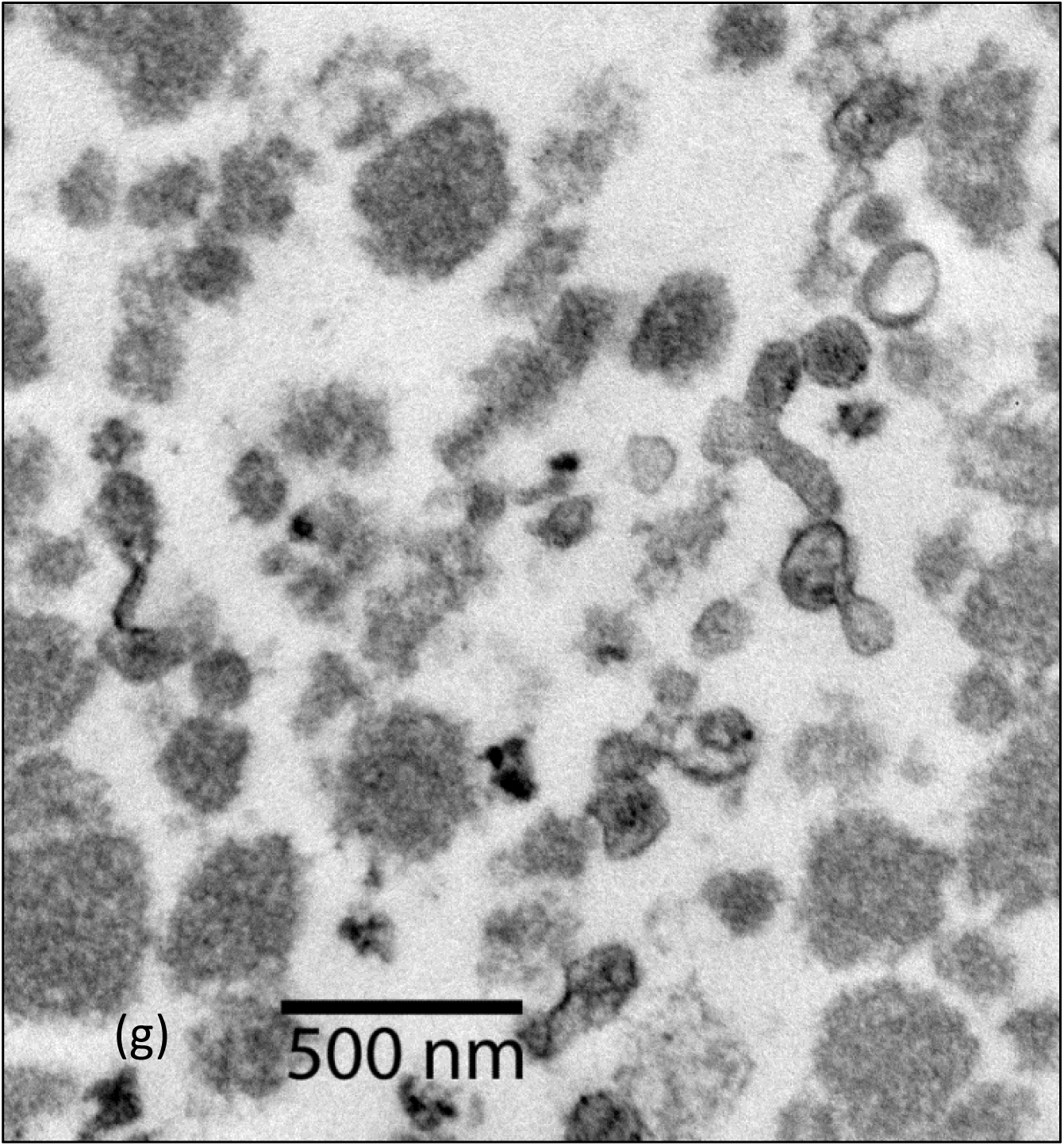
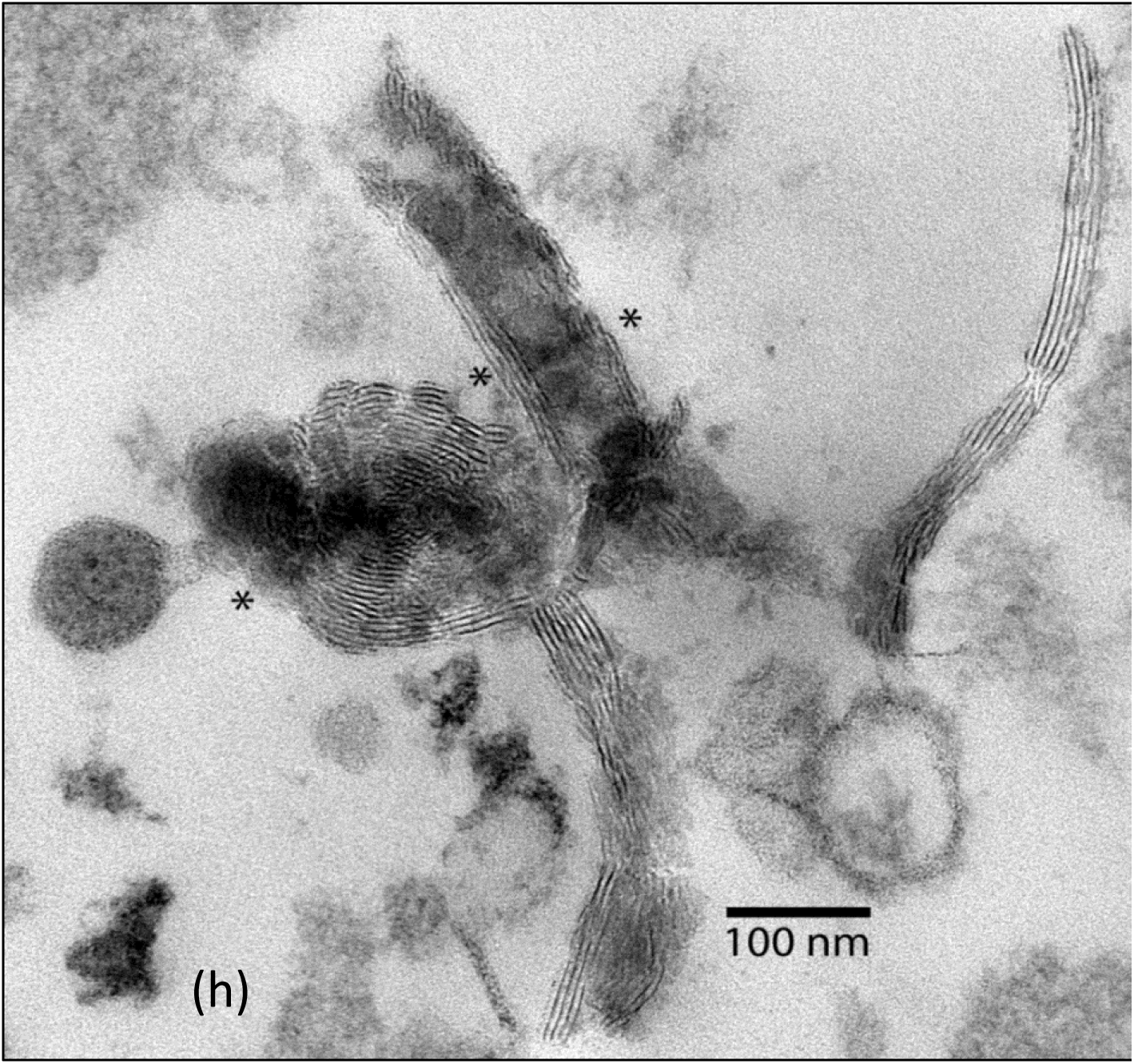
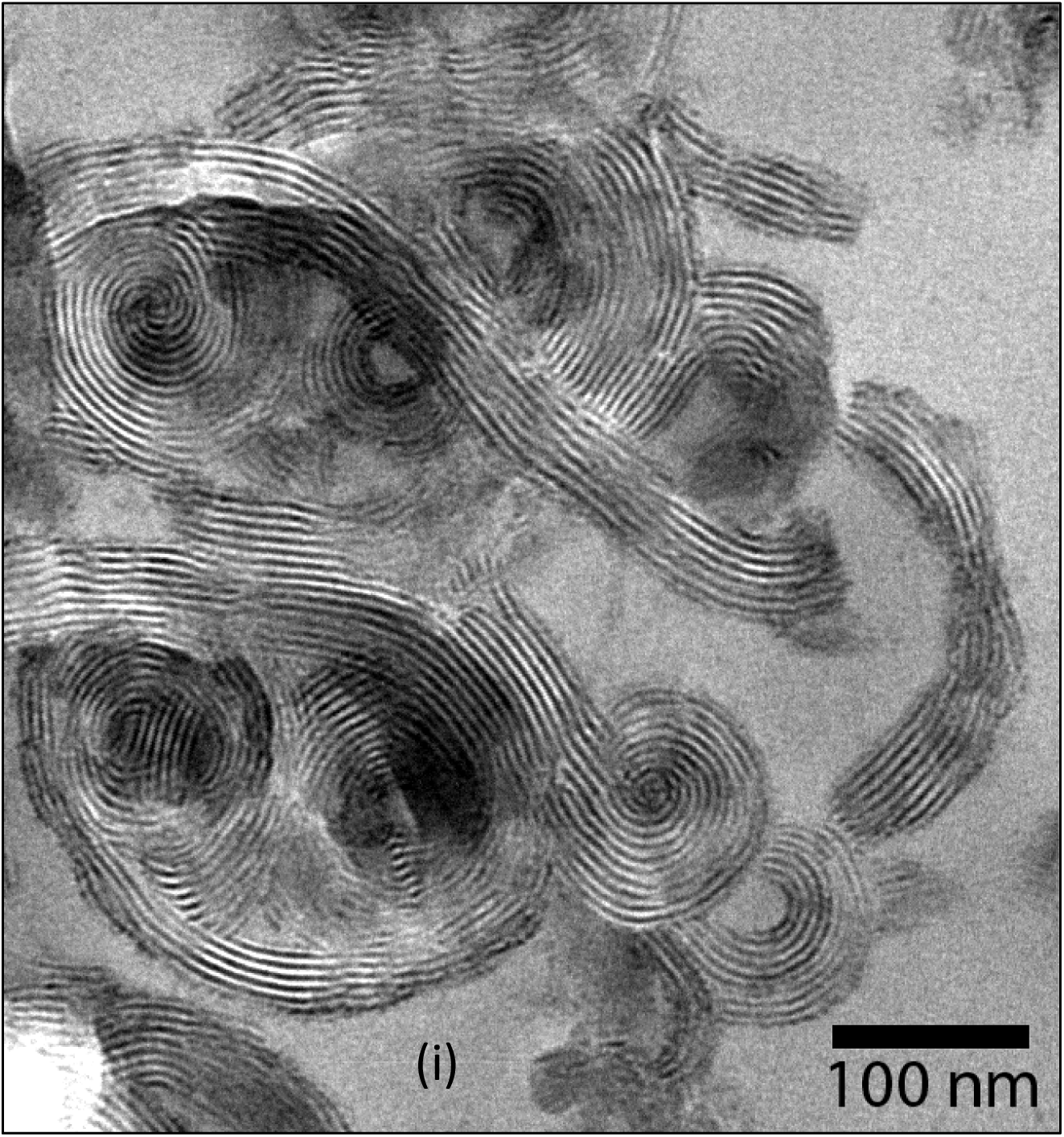
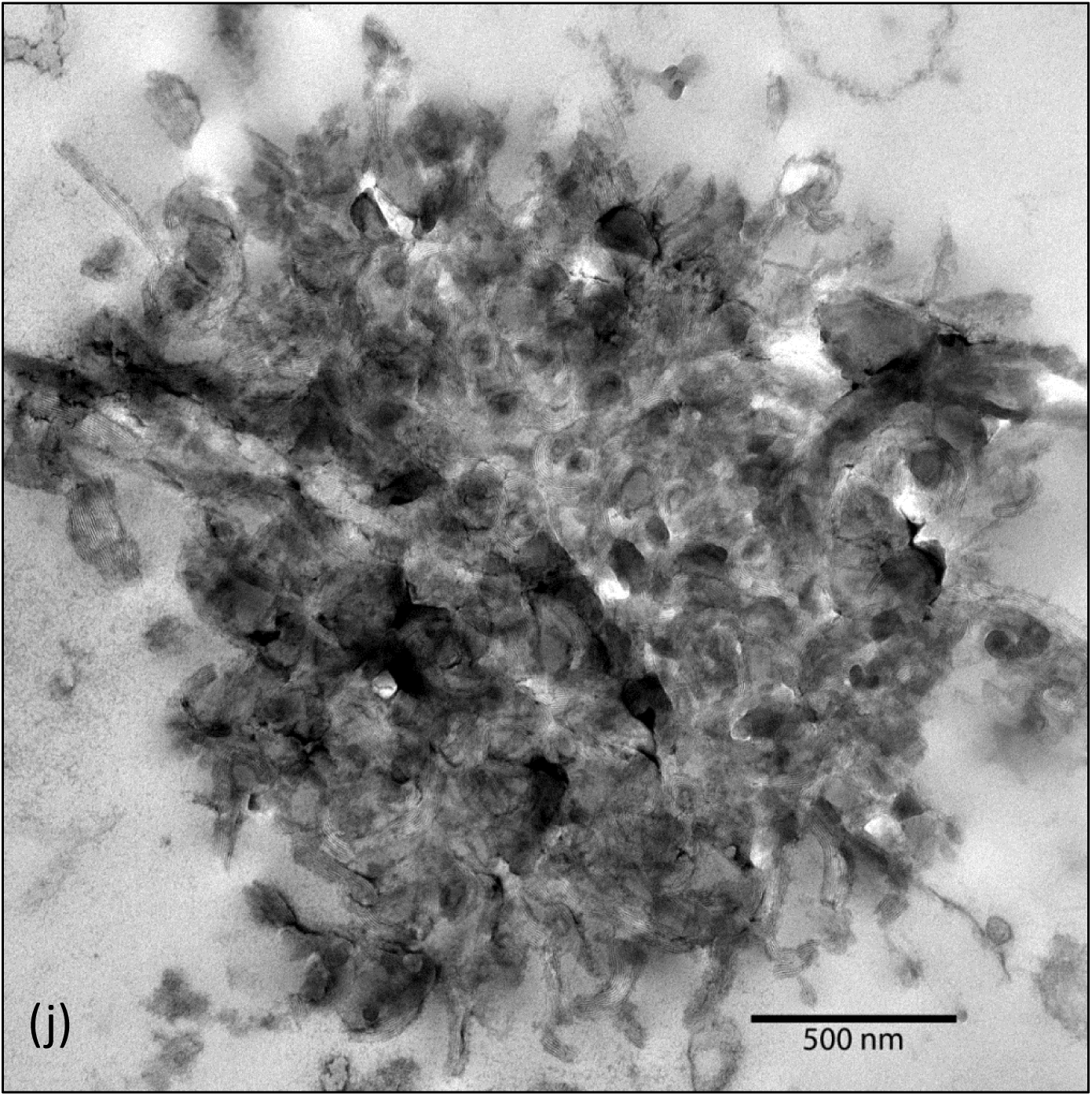
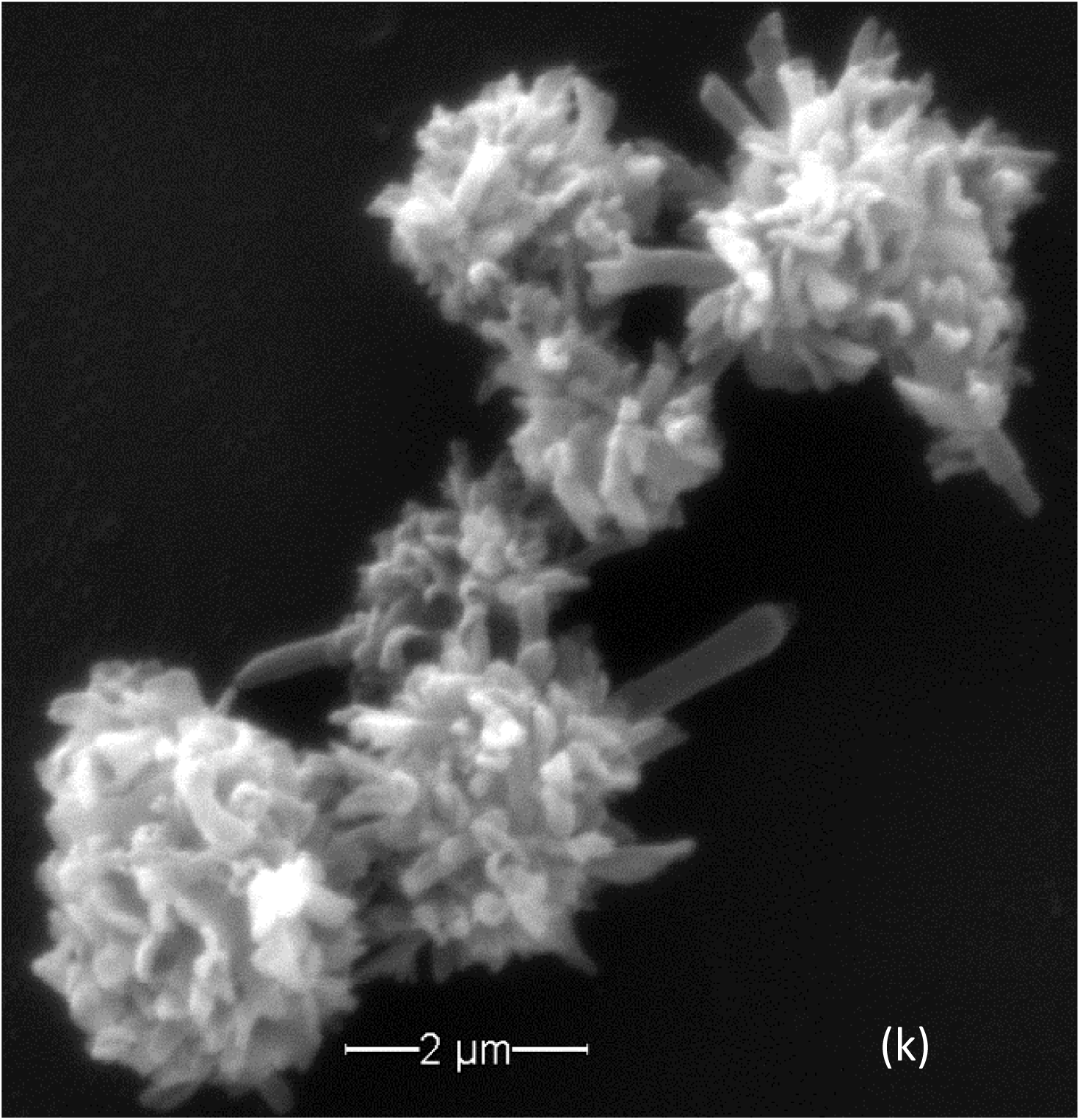
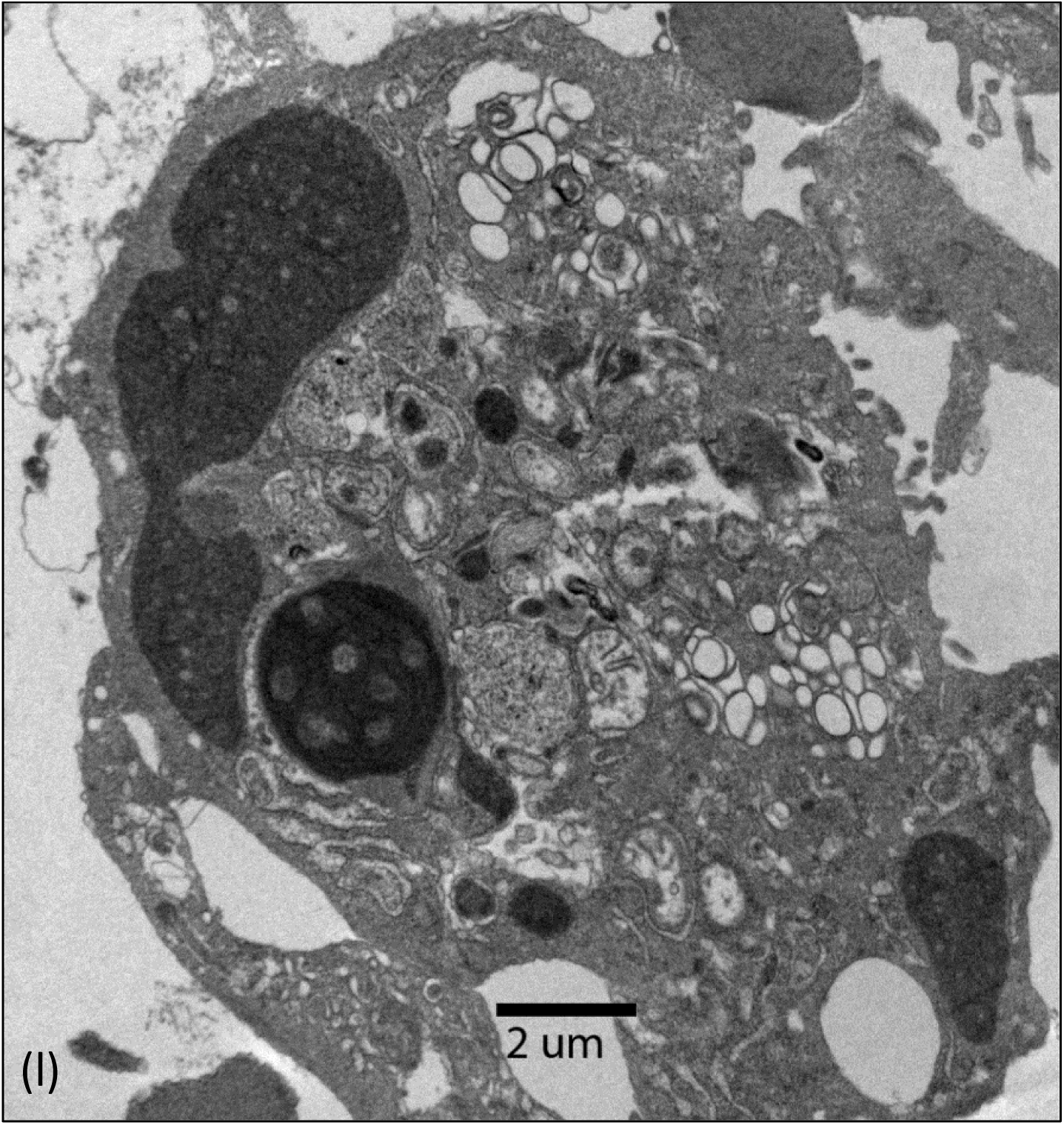
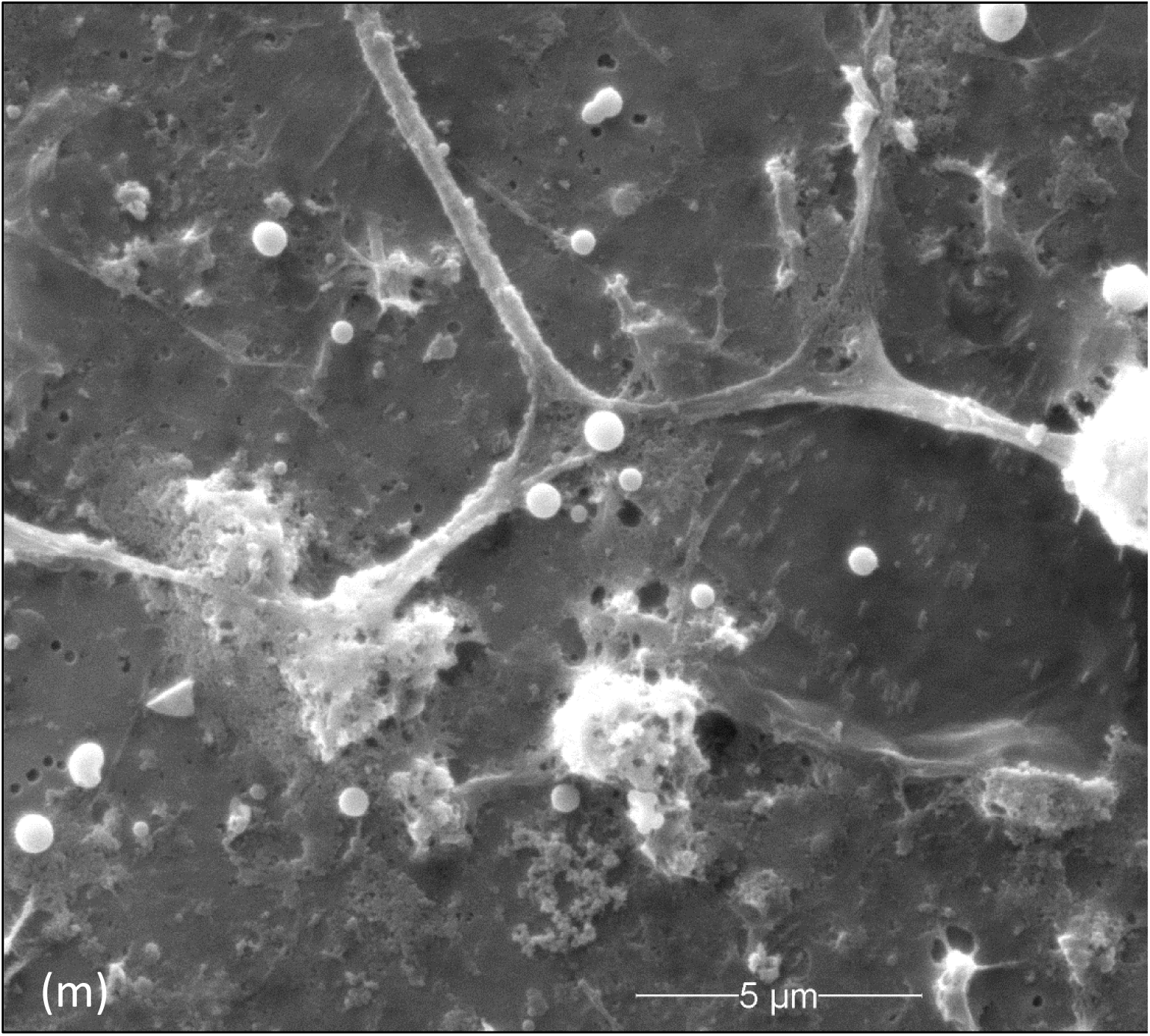
Microorganisms recovered from sheep with scrapie also propagate outside the host. (a)TEM negative stain of the scrapie isolate grown in broth. (b) Light microscopic examples of the scrapie agent grown in broth. When first passed they are mostly as individuals, but as they propagate they begin to aggregate. Blue is DAPI, a fluorescent DNA dye. Comparisons are by interference microscopy. Bar 5μ (d) and (e) By DAPI stain (d) and interference microscopy comparison (e) they are pleomorphic. and intertwine. In real time they change constantly. Bar 5μ The intertwined individuals can be detected by SEM (f). (f) Individual Cell-free forms intertwine as shown by SEM, and when cultures are dense, the microorganisms disintegrate, secreting their constituents (g) (g) Debris from disintegrated microorganisms are shown. This process enables their collective self-assembly into large highly organized compact structures. (h) Materials secreted by scrapie isolates, grown in broth, begin a self-assembly process forming large, highly-ordered structures. Asterics* mark areas of active transitions into ordered arrays. (i) High TEM magnification shows the highly organized products that form dense inclusions (j). These structures stain positive for nucleic acids and polysaccharides. By SEM (j) they reveal numerous rough edges and spines. For this reason they are called “microbial hedgehogs”. This cooperative process involving many individuals to produce this structure is unique among microorganisms. (k) Following months of continuous cell passages in broth, the cultures were held for weeks in broth until the large structures were formed. When then added to host cells they reestablishing the characteristic intracellular infections, completing an infectious cycle (l). (m) Biofilm artifact. The carbohydrate rich material released into solution (g) could be mistaken for a biofilm if attached to a surface, as shown by SEM. But the isolate from sheep with scrapie does not form biofilms.

#### Cell Free Growth and Product Development

When grown in broth, scrapie isolates and SMCA show minimal constituents. During early phases of growth, they are fragile. When processed gently prepared for TEM negative stain they are long, thin, branch and reveal blebs containing electron dense material (Fig. 2, a, and 5, a).

Cell free growth requires prokaryote mechanisms of protein synthesis. PCR tests for 16s rDNA showed positive results for scrapie isolates and SMCA (Fig. 3, a), consistent with their ability to grow cell free.

**Figure 3:**
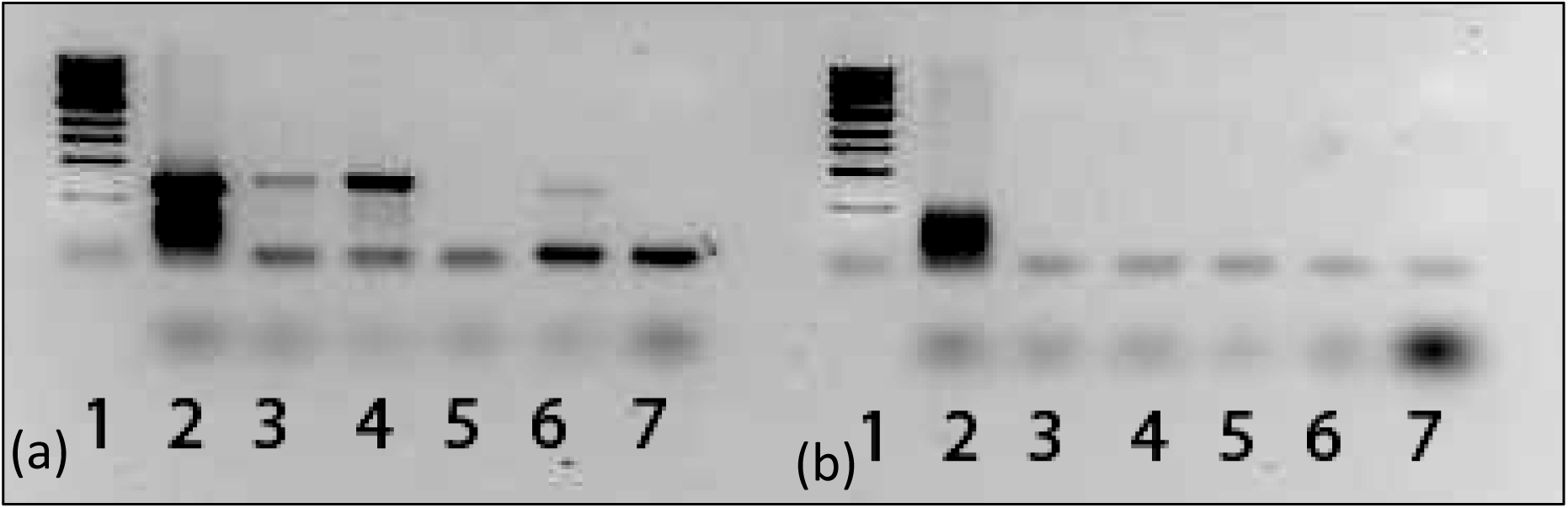
PCR of 16s rDNA and Hsp60. (a) 16s rDNA: lane 1, DNA ladder; lane 2, *S. citir;* lane 3, case #5061; lane 4, case #301; lane 5, case #296; lane 6, SMCA; lane 7, primers only. (b) Hsp60: lane 1, DNA ladder; lane 2, *S. citir;* lane 3, case #5061; lane 4, case #301; lane 5, case #296; lane 6, SMCA; lane 7, primers only.

Only the cell free form the scrapie isolate was studied further. When grown to high concentrations, individuals intertwine as clusters (Fig. 2, b-f). Subsequently, these fragile forms of the scrapie isolate disintegrate, releasing their constituents into the medium (Fig. 2, g). Oddly they seem to have self-destructed; they look dead. However, these microorganisms are new to science and rather than draw conclusions it is best to just follow the data. Their debris contains a carbohydrate type polymer that stains with ruthenium red and mimics a biofilm when attached to a surface (Fig. 2, m), though because this material is produced by freely growing organisms and not subsequent to surface attachment they are not biofilm producers. Remarkably, what appears as collective wreckage from many microorganisms actually self-assembles into large highly organized structures (Fig. 2, h-k), an energy requiring process. However, they lack an obvious way to produce the required energy. This process of assembly appears as dehydration driven by the energy of hydrogen bond formation between water molecules. Dehydration is a preservation process, which would be useful to protect the pathogen’s genome. The end products are compact structures comprised of smooth alternating layers with rough surface edges (Fig. 2, i-k). Due to their bristly surfaces these products were nicknamed microbial ‘hedgehogs’ by Sokolova. To produce just one ‘hedgehog’ requires sacrifices of many, an interesting example of microbial social behavior to preserve the species genome. Since these forms are fragile during the growth phase and form ‘hedgehogs’ only at high population densities, this process seems suitable for small protected areas. It is possible that they grow within the eye aqueous of infected animals, and are then shed into the environment or within vectors that die in fields if they cannot find a suitable host. The condensed ‘hedgehog’ forms are suggested as structures to protect the pathogen’s genome during long periods between hosts. This would explain the seeding of fields grazed by sheep with scrapie that remain infectious to other sheep for years (17, 18). These large structures, assembled from debris, are infectious to cells. When ‘hedgehogs’ are added to the host BCE, they infected those cells producing the same characteristic infection as the original isolates from sheep with scrapie, completing an infectious cycle (Fig. 2, k, l), at least under laboratory conditions. If the intracellular form of the scrapie isolate predominates its life cycle, and cell free growth is limited, these microorganisms would not require their own Hsp60 gene. PCR tests indicate they lack the spiroplasma Hsp60 gene (Fig. 3, b).

Ultrastructural methods revealed the existence of the fibrillary VLP mechanism of forming inclusions and other structures of the two different microorganisms studied. This distinctive mechanism enables the infected cell to produce varied products of the pathogen, most of unknown significance. Ultrastructure also revealed formation of transient spiral forms and showed the unique properties of extracellular growth, including assembly of highly organized structures produced by the scrapie isolate named ‘microbial hedgehogs’. It shows that the patterns of cell pathology of scrapie isolates and SMCA are similar to those from CJD and MS biopsy materials (19, their fig. 2 and 20, their fig. 5), reproduced here as Fig. I and II when permission can be obtained. But ultrastructure has noteworthy limitations.

As a caution, the diversity of inclusions shown establishes the genetic flexibility of the granular fibrillary VLP mechanism. These intracellular processes can form or mimic almost any inclusion or body described as associated with different neurodegenerative diseases, without necessarily bearing relevance to the specific agent of cause. Only some of the inclusion types are shown. As examples of genetic flexibility of the fibrillary VLP mechanism three inclusions, formed by the same SMCA agent infecting the same cell line, are shown (Fig. 6, a-c). The most common type of inclusion is Figure 6, a. Figure 6 b shows formation of minimal type bodies (white arrow) and simultaneous formation of structures resembling pseudoviral hollow-cored vesicles (21), or as in Figure 6 c, this agent also forms large circular arrays. What controls the products formed is a mystery. The structures shown must mean something, but all we know is they exist. This granular fibrillary mechanism is new to science and both agent examples are associated with neurodegenerative diseases. Figures 1h, 1m, and 6 b are clear examples of forming products using viral mechanisms of assembly as are the formations of the large inclusions freely within the host cell cytoplasm. Figures 2, a-f and 5, a) show these same agents can multiply in broth, a property of prokaryotes and not of viruses. Collectively the data indicate they are hybrid microorganisms. But ultrastructure has limitations. Applying more varied approaches to study these organisms is essential to understand them. It is also improbable that microorganisms with small genomes could encode all the intracellular process and extracellular functions shown. Perhaps when intracellular they activate ancient genes of pathogens embedded in mammalian genomes as suggested by Bandea for prions (22).

### 4.2 The Relevance of Scrapie Isolates and SMCA to the Literature on Infectious CNS Neurodegeneration

The data shown provides insight into diverse theories of infectious CNS neurodegeneration and implicates types of pathogens that could cause CNS neurodegenerations. Biopsy tissues from neurodegenerative cases are used as standards to interpret lab data and evaluate theories of infectious cause. Patient biopsy samples from cases of CJD (19) and MS (20) were chosen as to represent two dissimilar but significant neurodegenerative diseases. Remarkably, when viewed side-by-side, reproduced as Fig. I and II, they show similar structures and cell pathologies. Cells infected with either SMCA or isolates from sheep with scrapie display features similar to each other and to CJD and MS biopsy data implicating a common pathogen ancestry.

Half a century ago CNS neurodegenerative diseases were labeled ‘slow virus’ diseases, with scrapie affecting sheep (the prototype) and CJD (a human equivalent). However failure to recover viruses led to four other theories of infectious cause. Manuelidis et al. proposed ‘VLP’ as causal agents (23, 24). Her VLP theory arose from the ‘slow virus’ period following detection of VLP in biopsy material form a CJD case (19, fig. 2) and finding VLP in CNS biopsy material from a sheep with natural scrapie (25). Manuelidis et al. established that infectivity resides in cell fractions separate from prions (24, 26, and 27). Though scientists with alternative views expressed doubt about her work (28), her VLP observations are consistent with CJD biopsy data. In her model prions cause pathology but are not the infectious causes. Payne and Sibley’s data and high magnifications of VLP (19) reveal VLP as comprised of rather uniform aggregations and continuous with long strands of granular fibrillary material. Data by Narang and Field show VLP can exist in at least two sizes (20). Studies of scrapie isolates and SMCA indicate that VLP are products of a pathogen’s intracellular processes (Fig. 1, c-h, Fig. 4, b, g, h), and not themselves the causal agents.

**Figure 4.**
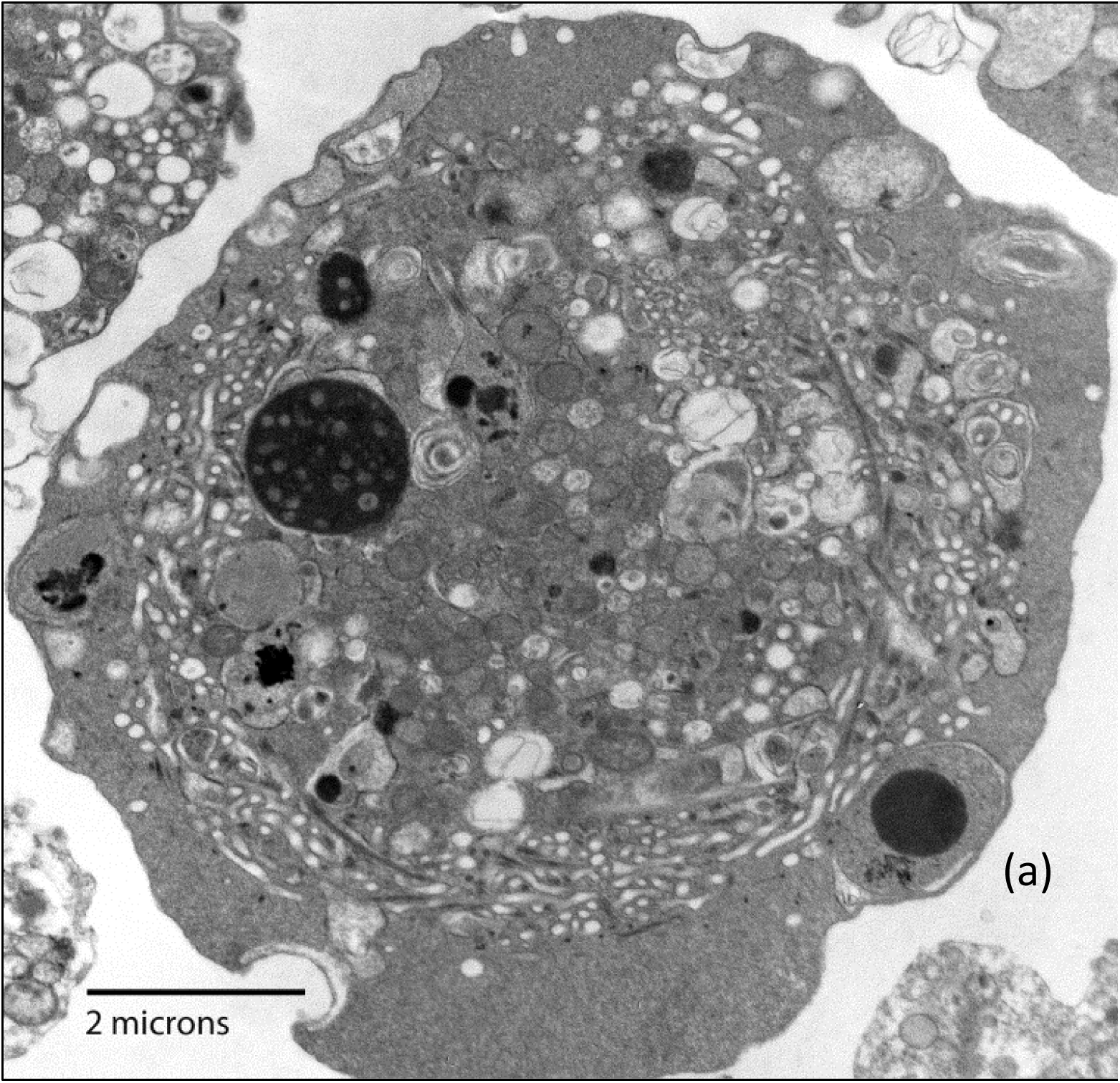
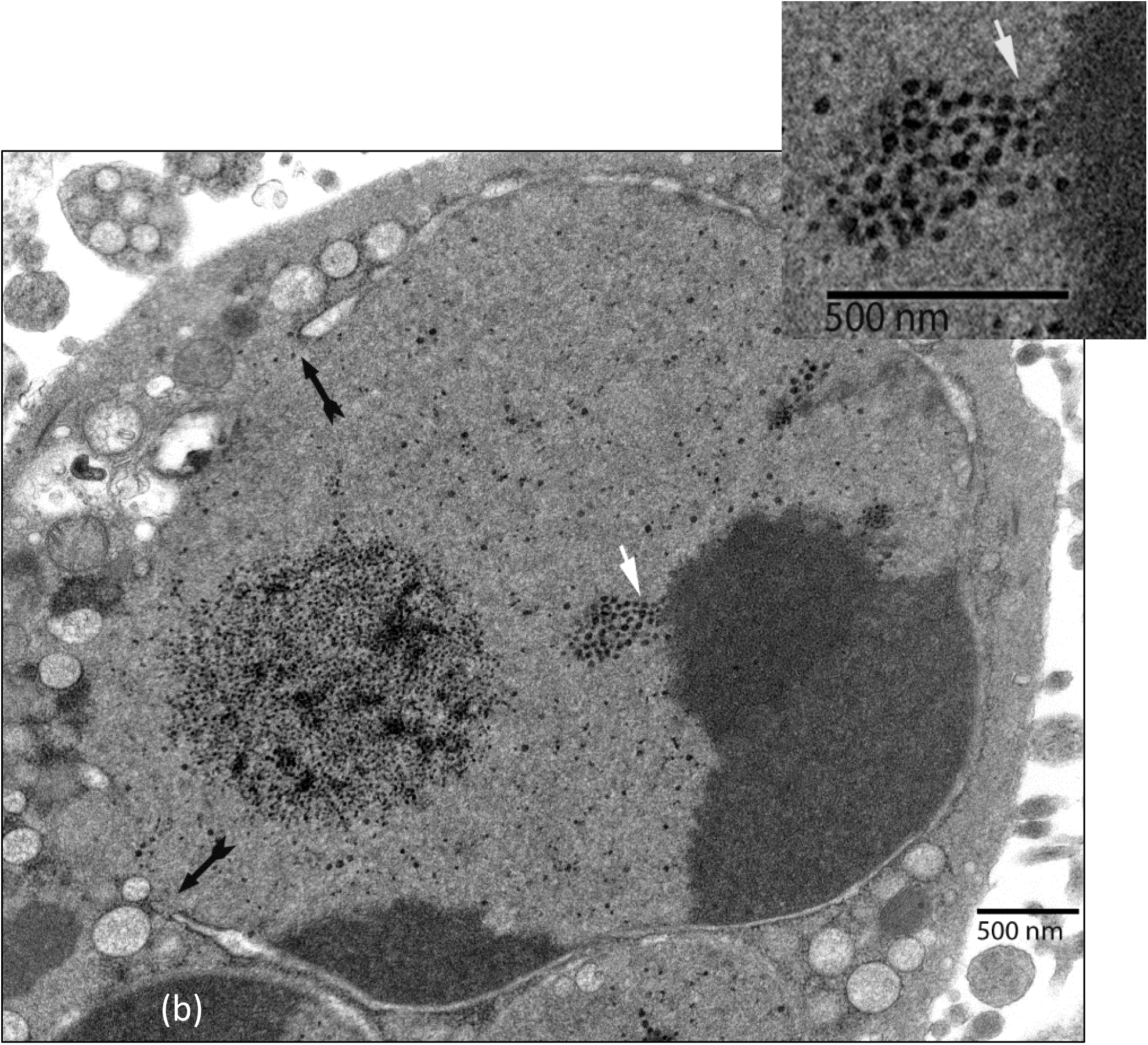
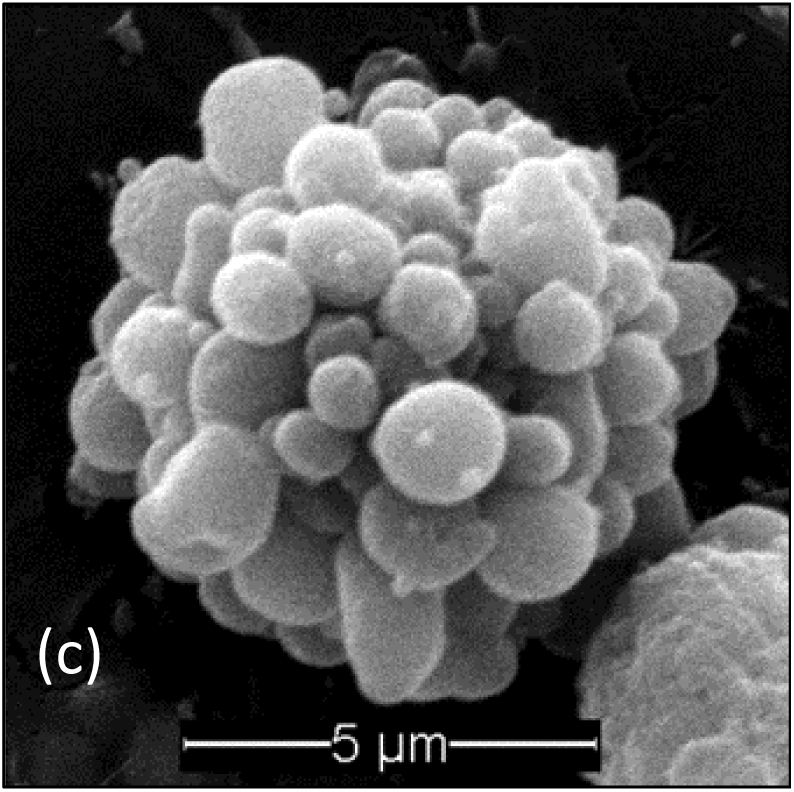
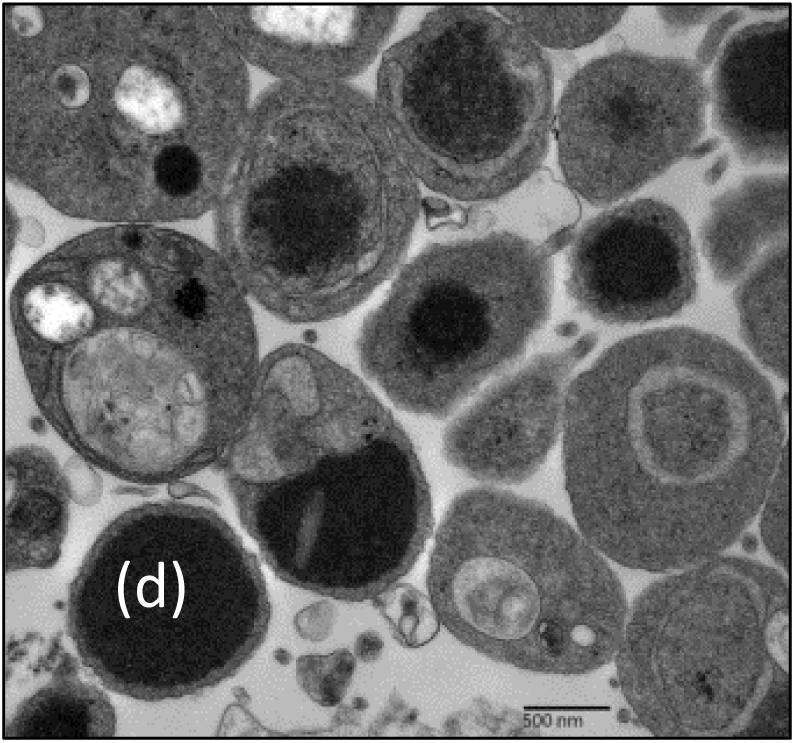
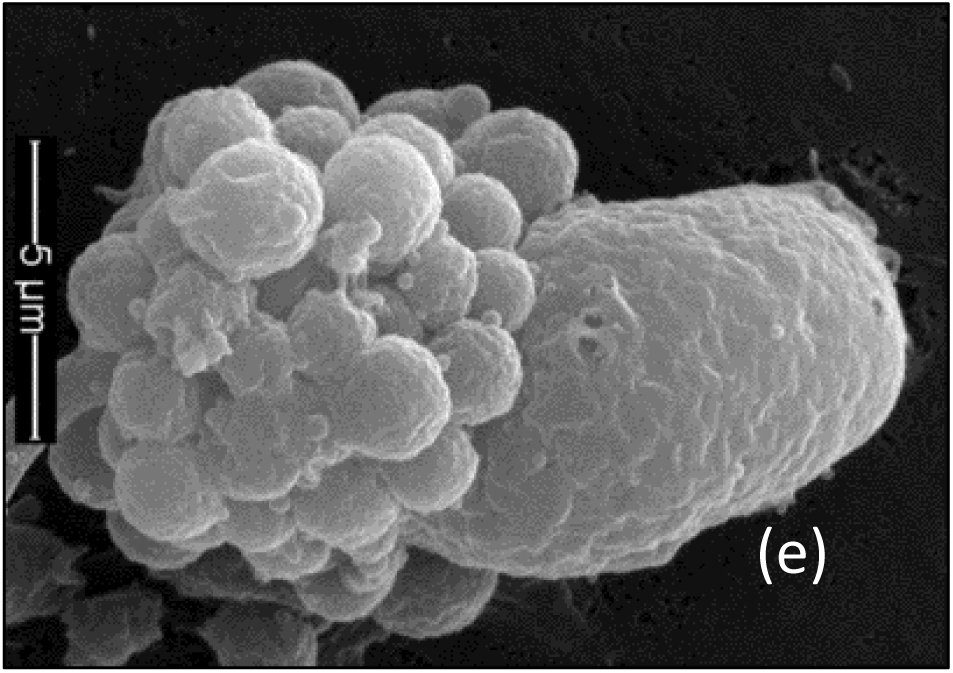
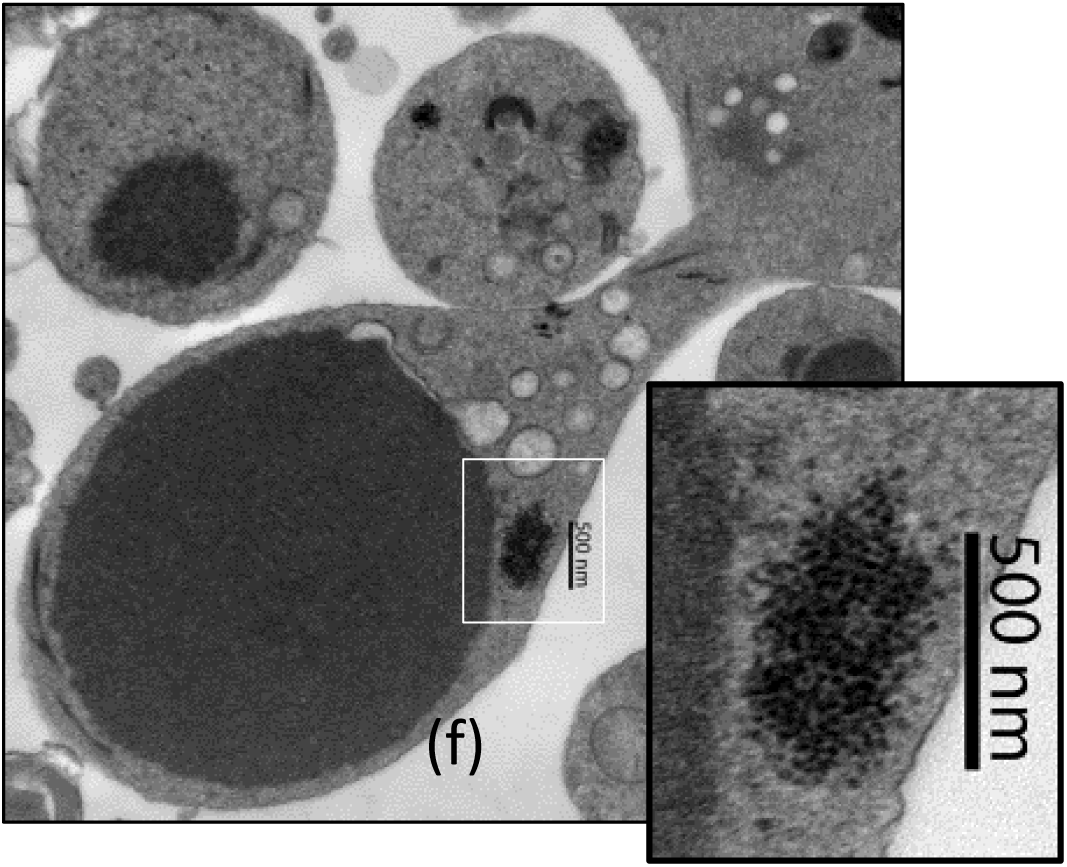
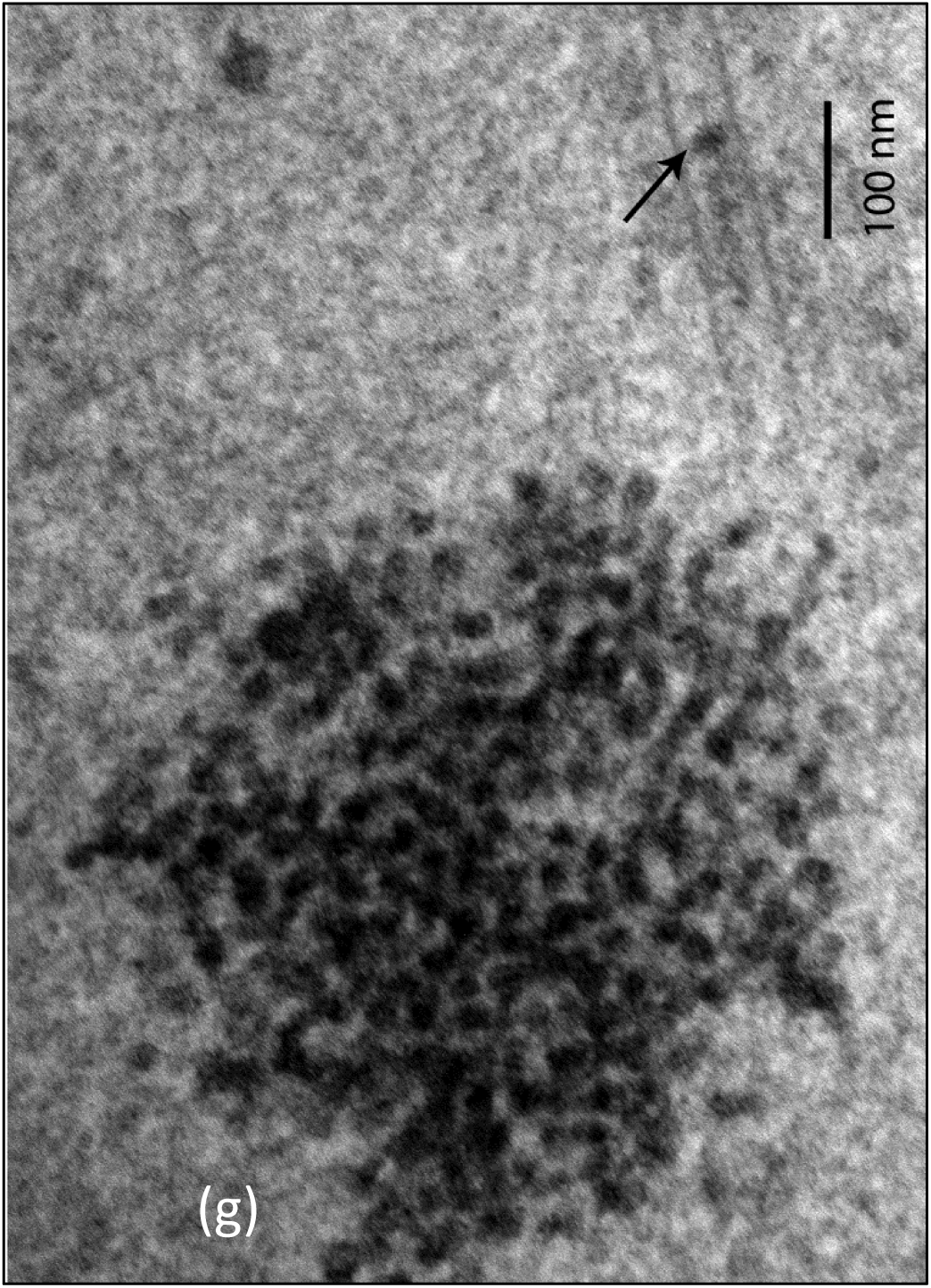
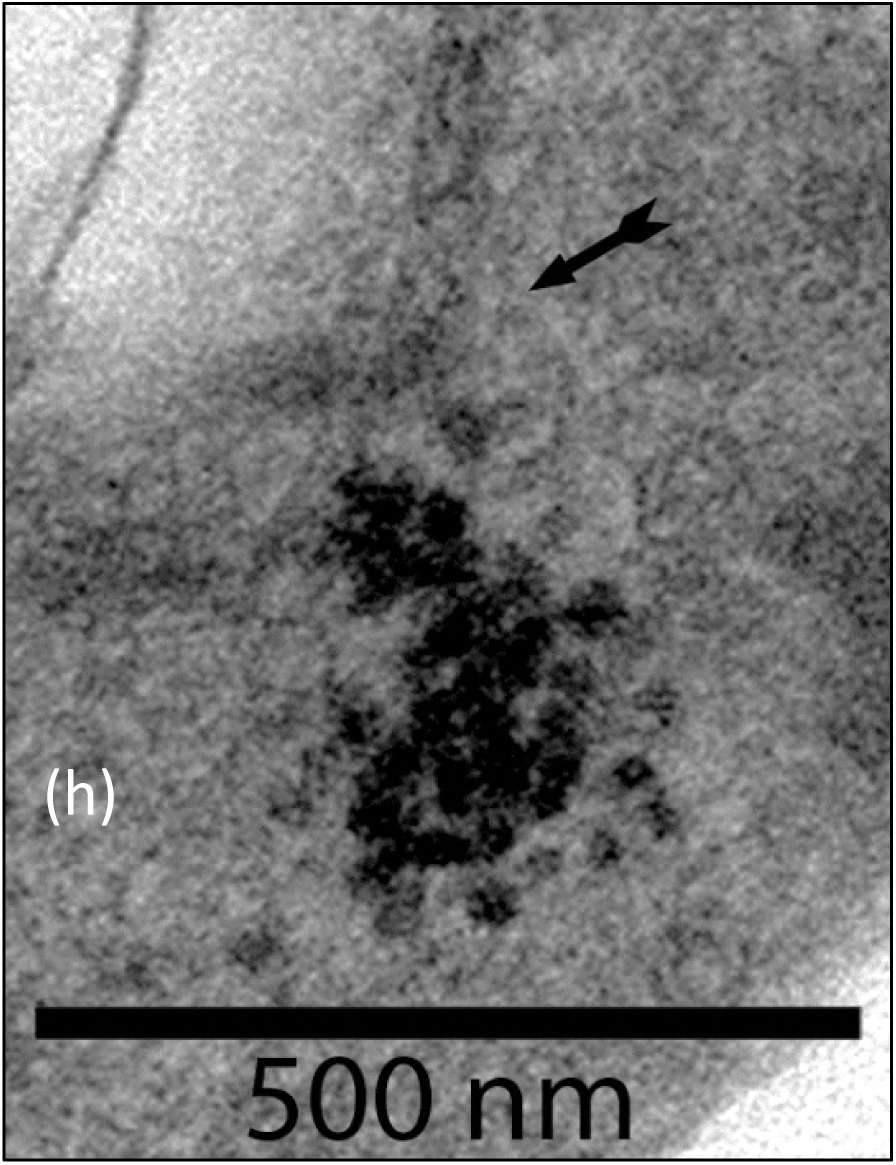
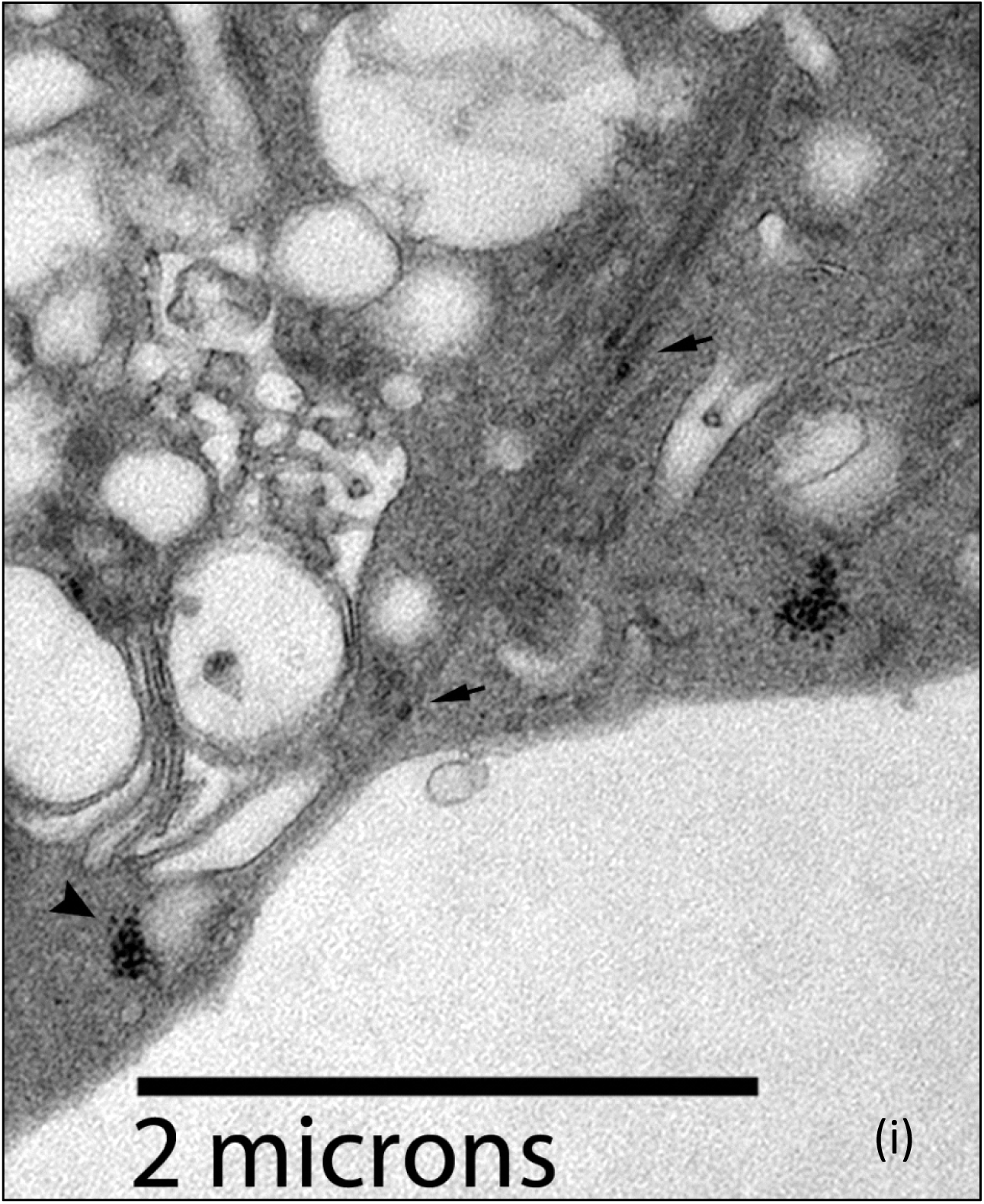
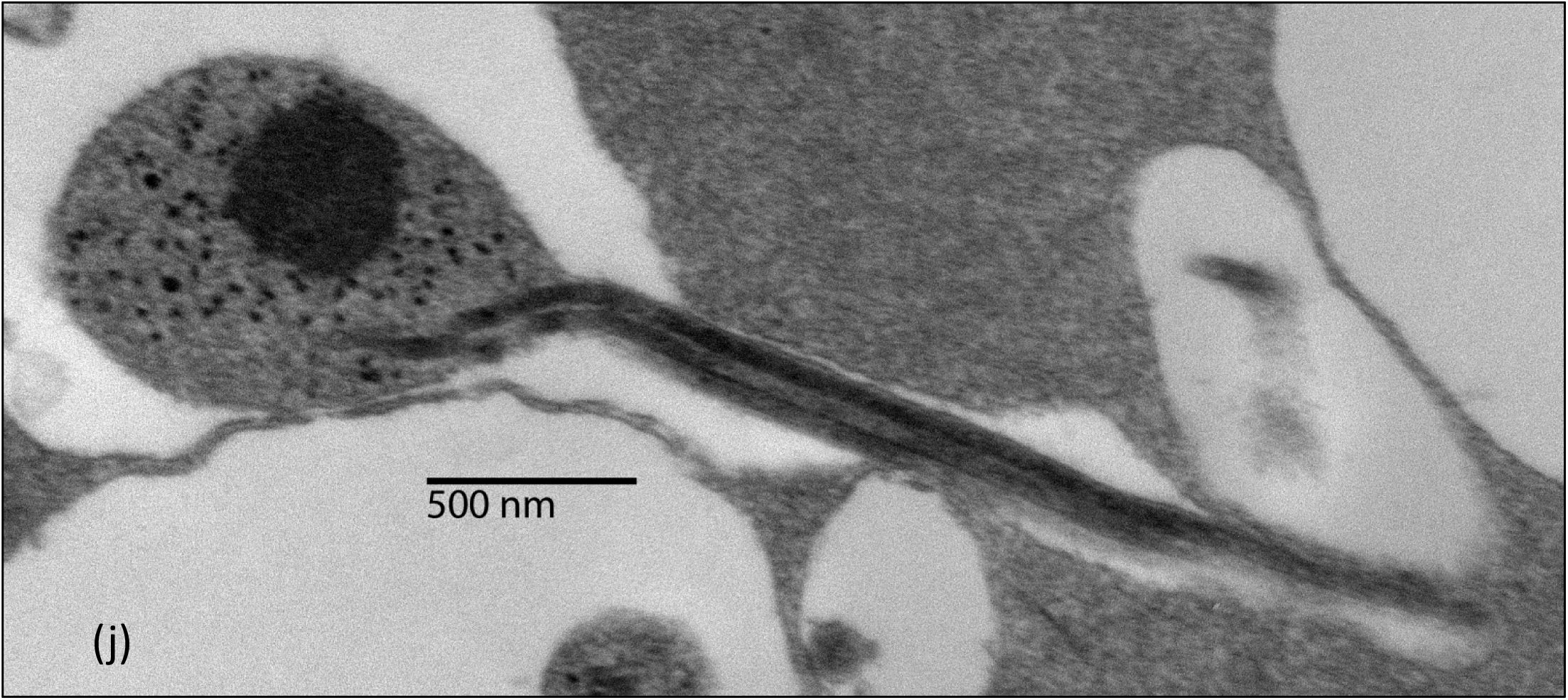
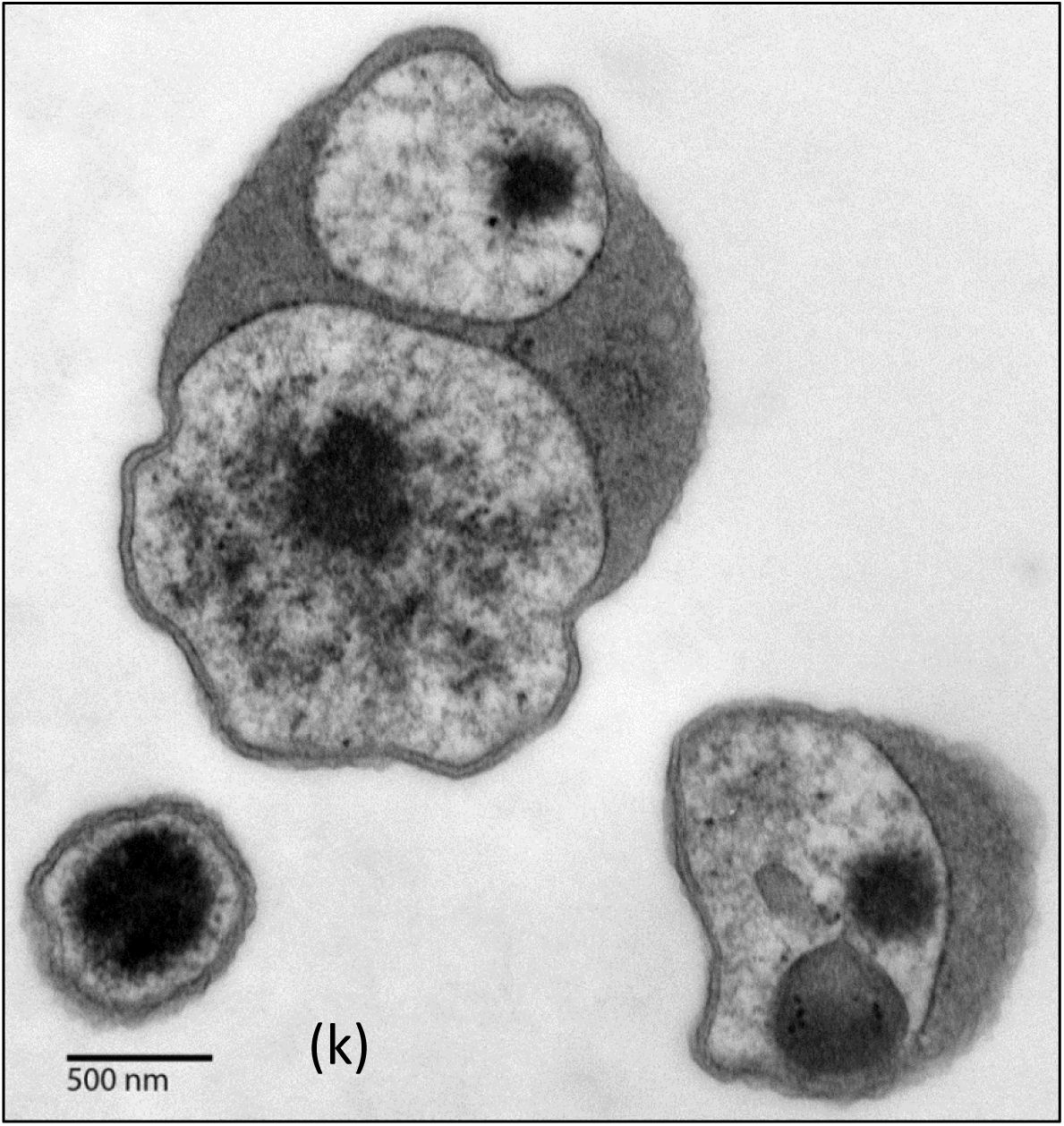
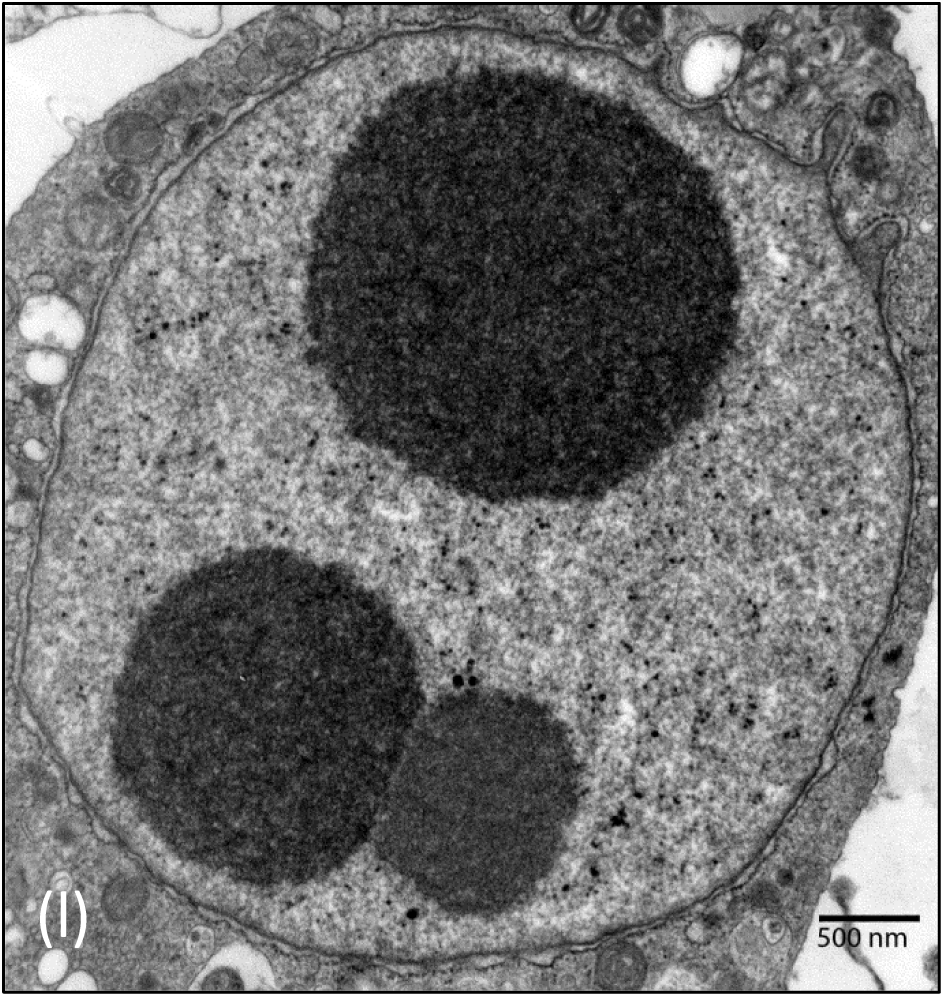
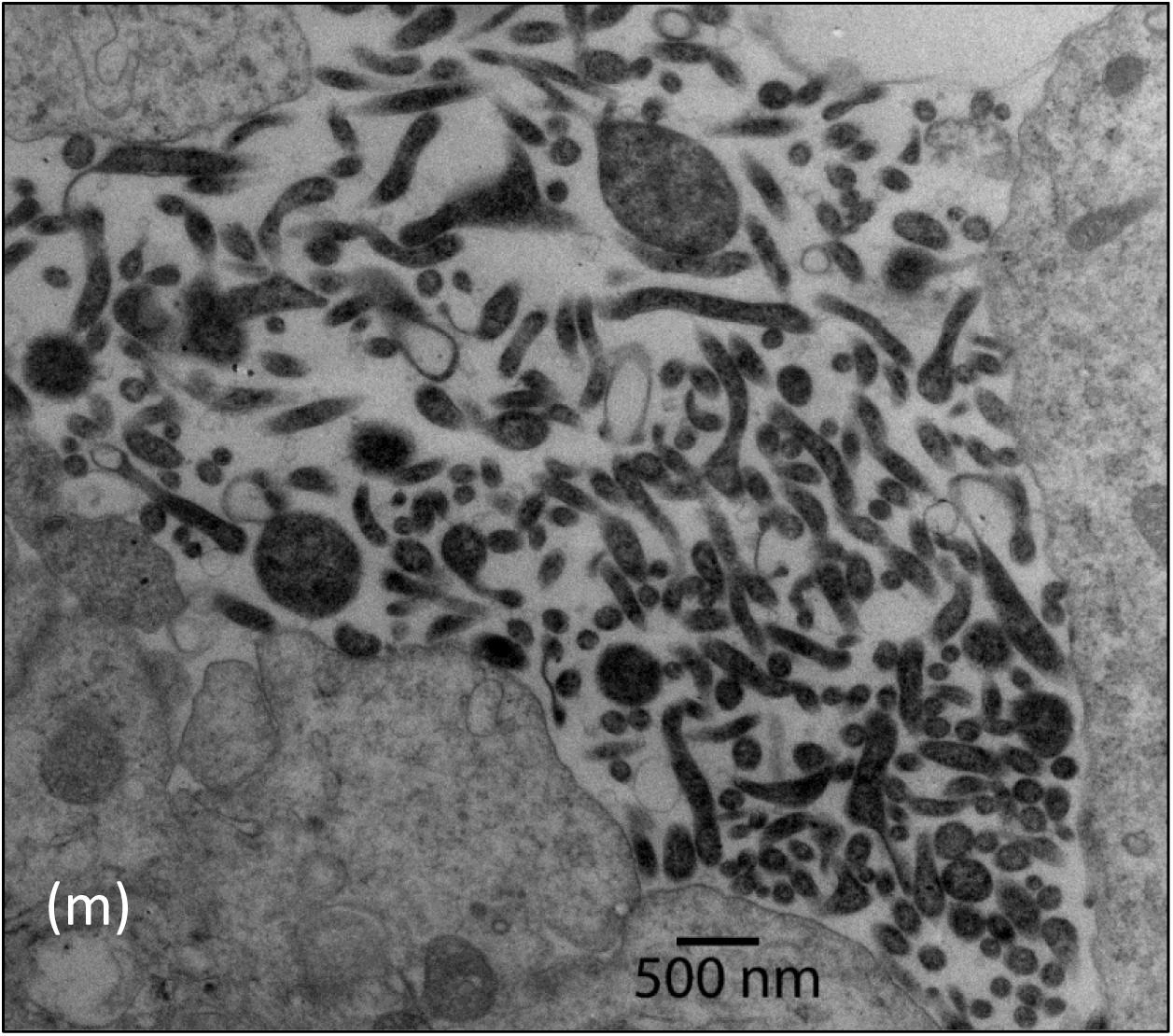
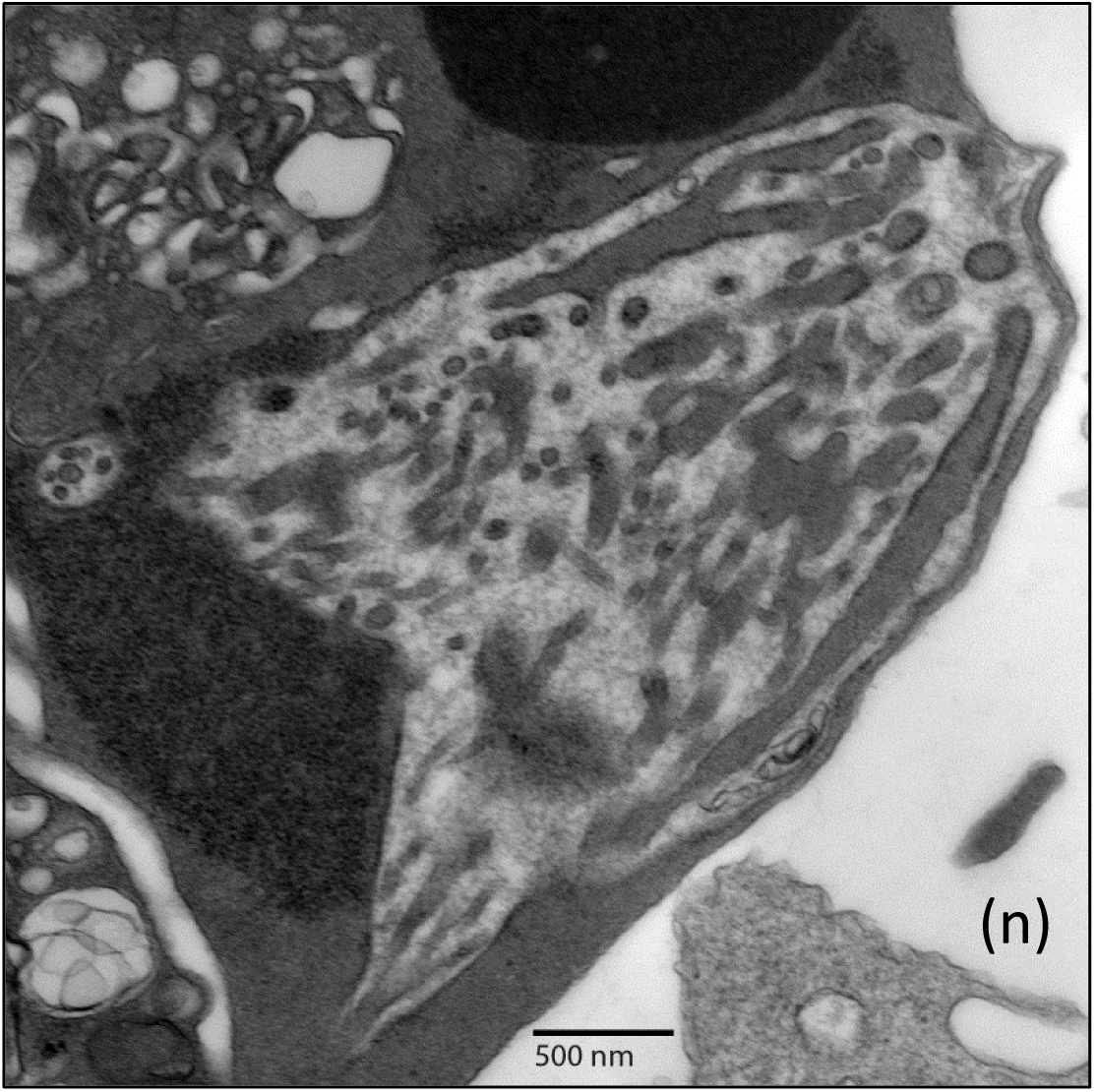
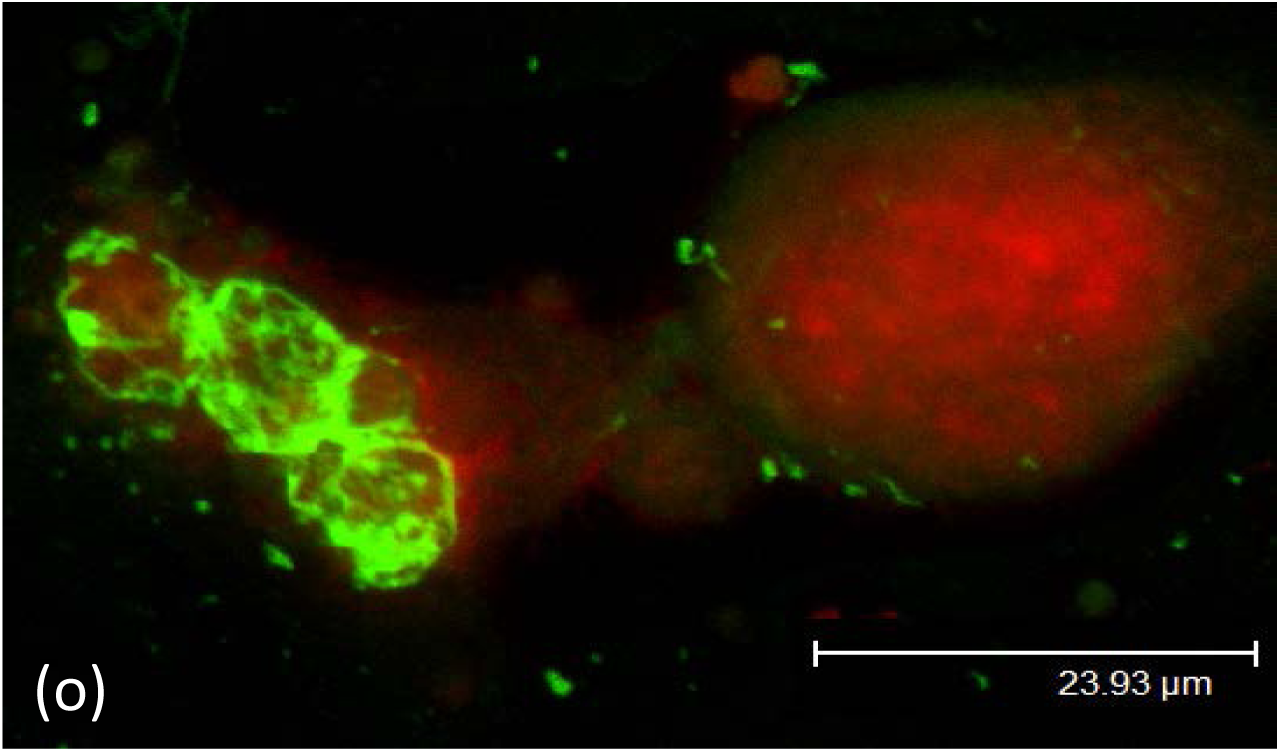
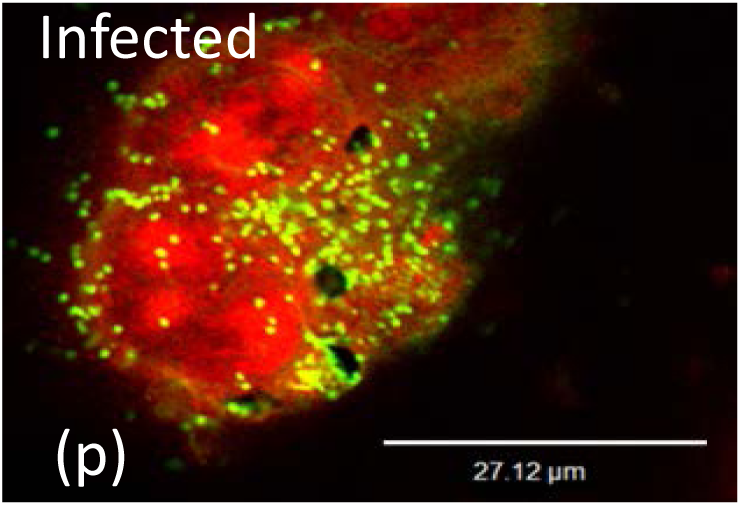
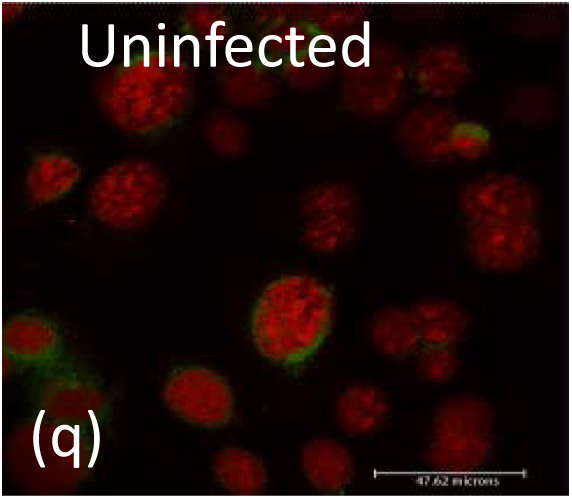
The Effects of SMCA infection on host cells. (a) SMCA-infected host cell; note the complete destruction of host organelles. (b) VLP form in the nucleus (white arrows) and inset at high magnification. The nuclear membrane opens (black arrows), forming continuity between the nuclear contents and those of the cytoplasm. One feature of apoptosis is formation of blebs as shown here with SMCA-infected cells. (c) and (e) SEM. (d) and (f) TEM thin sections through similar areas. At high magnification they reveal VLP (g) and (h) Though difficult to detect, VLP within the infected cell cytoplasm apparently can unravel and enter into tubules (arrows). (i) The association of VLP with tubules can be lengthly; perhaps this material can transverse via tubules (small arrows) and reform VLP elsewhere (arrowhead). (j) This tubulovesicular structure shows continuity between vesicles with VLP and tubules containing granular fibrillary material. Sometimes minimal type bodies seem to form by condensation while they also can form by packing strands of VLP. Lewy bodies are inclusions with a ‘fried egg’ appearance and associated with the α-synucleinopathies, discovered first in patients with Parkinson‘s. This example is from a cell infected with SMCA, and those cells tested positive for aggregations of α-synuclein by FA (7). They appear to form in the nucleus under certain culture conditions, and are likely of the type associated with MSA. (m) and (n) Spiroplasma-like forms can be detected especially after long incubations. As shown in (n) they appear to be assembled. Attempts to culture these forms cell free failed. These forms may be transitions between intracellular and extracellular processes of propagation. The form of SMCA that grows in cell free broth is shown in figure 5 (a). FITC-labeled antibodies, raised against SMCA grown in broth, were used to probe BCE cell cultures infected with SMCA. Red background stain, propodeum iodide. The antibodies decorate SMCA antigens in a manner consistent with assembly of structures; shapes of spiroplasma are rare. The antibodies used were provided by the Dept. Vet. Science, LSU Agcenter.

Tubulovesicular structures (TVS) were proposed as the ultrastructural hallmark for prion diseases (29, 30). Narang, Asher and Gajdusek describe the morphological and immunochemical relationship between tubulofilaments in scrapie-infected brains and scrapie-associated fibrils (31). They also provide evidence that DNA is present in these structures from scrapie tissues (32). The TVS and tubulofilaments are also in cells infected either by SMCA (Fig. 4 i, j) or isolates from scrapie sheep (Fig. 1, i-k). Cores of TVS and tubule-filaments are likewise filled with electron dense granular fibrillary material as originally described by Narang et al. (31, 32). These structures occasionally reveal VLP at the beginning and ends of the tubulofilaments (Fig. 4, g-i) implying a transport mechanism.

The spiroplasma theory of infectious cause arose from SMCA research. In 1964 Clark reported recovering an agent from ticks causing neurodegeneration in lab animals (33). Because cataracts were prominent in rodent models, Clark’s named his isolate the suckling mouse cataract agent. For a decade SMCA was studied as a ‘slow virus’ (33, 34). Then Schwartz and Elizan detected small mycoplasma-like forms associated with cells of SMCA-infected embrynated eggs. They could not resolve if these forms were separate or continuations of host cell membranes. Yet they still favored a virus model (34). From SMCA-infected eggs Tully et al. concentrated spiral shaped forms and reported they grew in broth (35, 36). Without testing the effect of SMCA on host cells, SMCA was classified as *S. mirum*, the type strain of a new spiroplasma species infecting mammals (37). However, their report of growing SMCA as a spiroplasma in broth is irreproducible in the senior author’s hands. What grows, using the same SMCA strain and broth formulation, are too fragile to survive their concentration methods. Furthermore, the forms that do propagate (Fig. 5 a) are similar to those from scrapie affected sheep (Fig. 2 a), and are different from those of any described genus.

**Figure 5.**
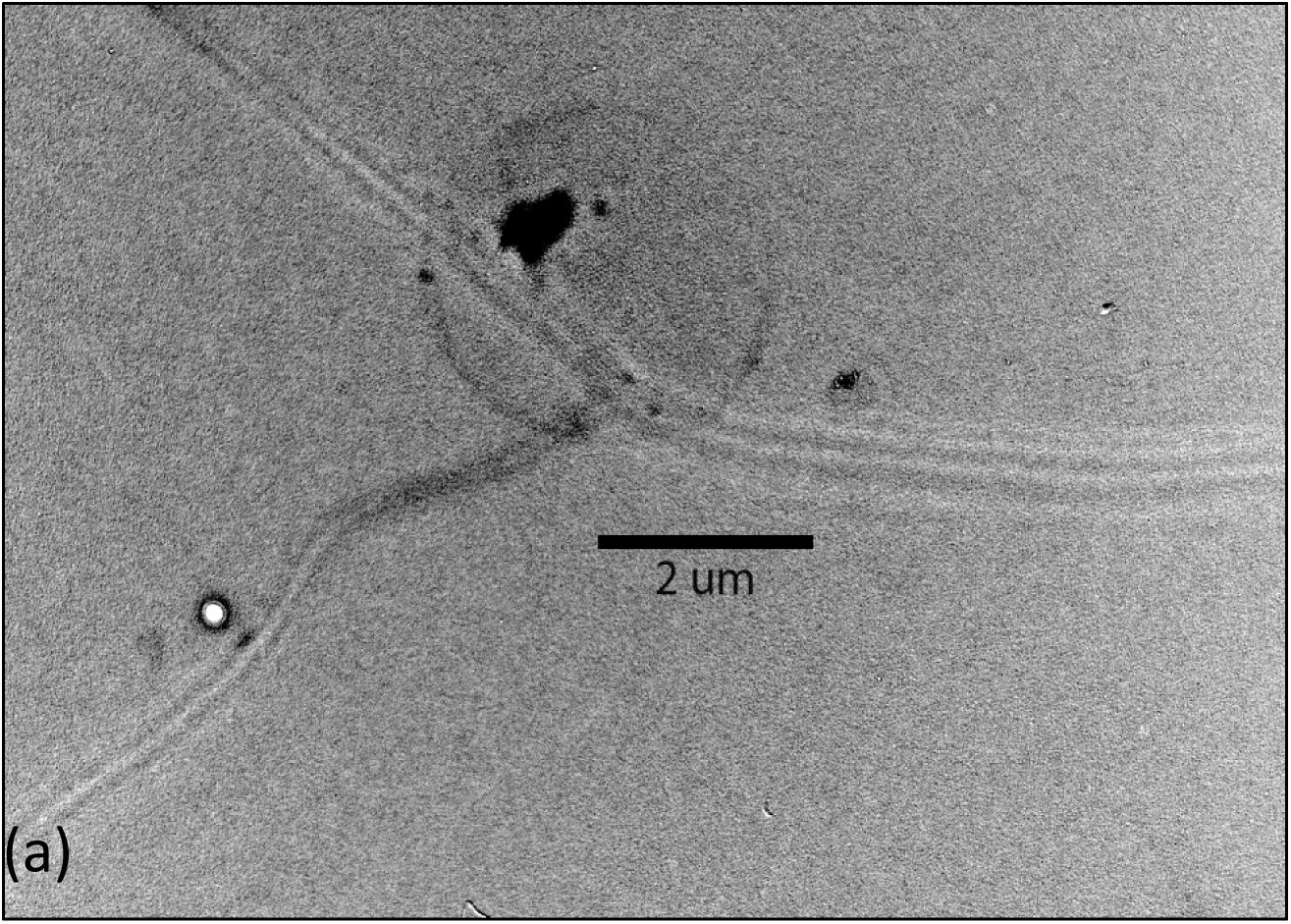
SMCA propagates in medium free of host cells. This form is ultrastructural indistinguishable by ultrastructure from that of agents recovered from sheep with scrapie. Negative stain from a culture of SMCA grown in broth. This growth form of SMCA has not been previously described.

**Figure 6.**
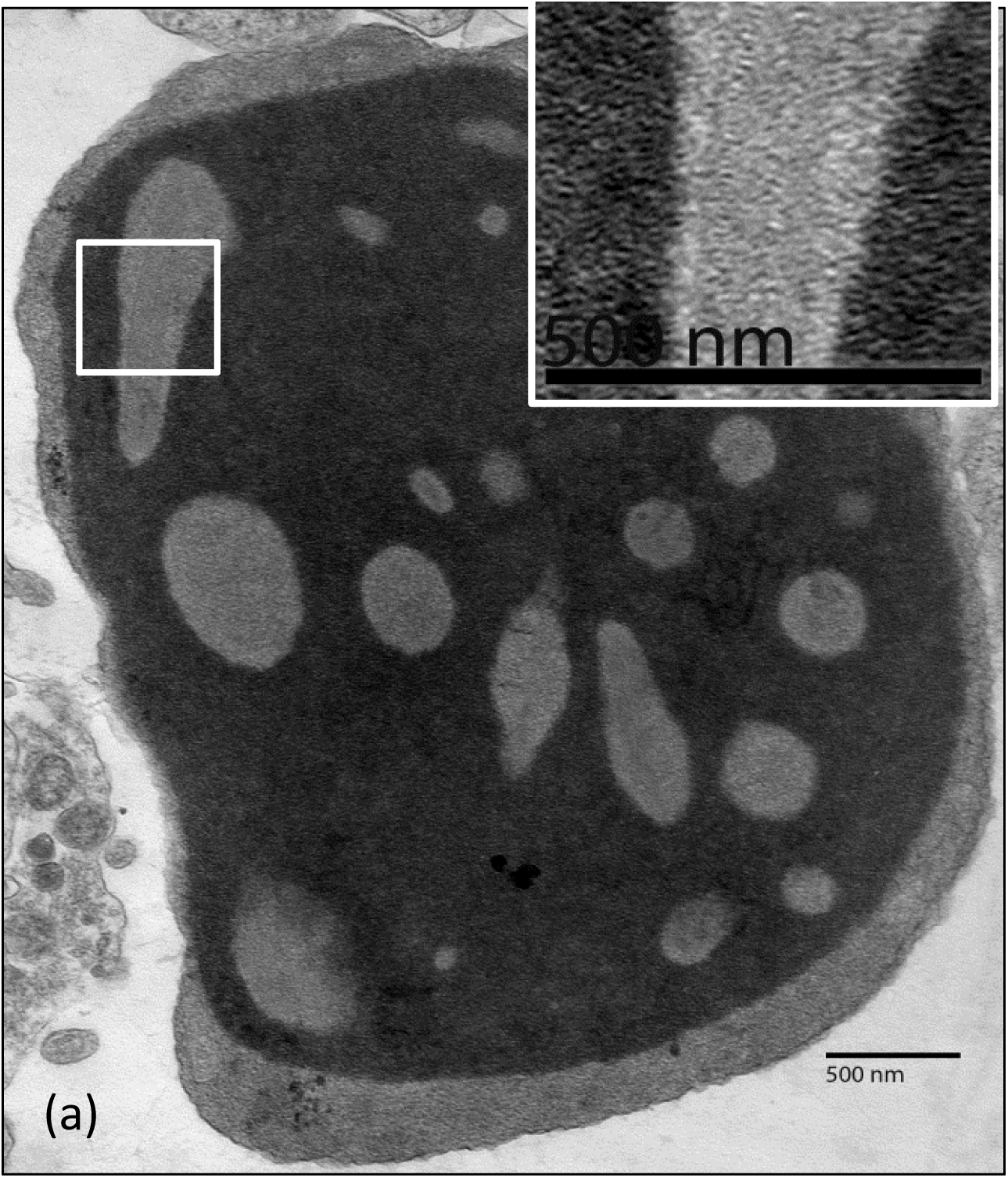
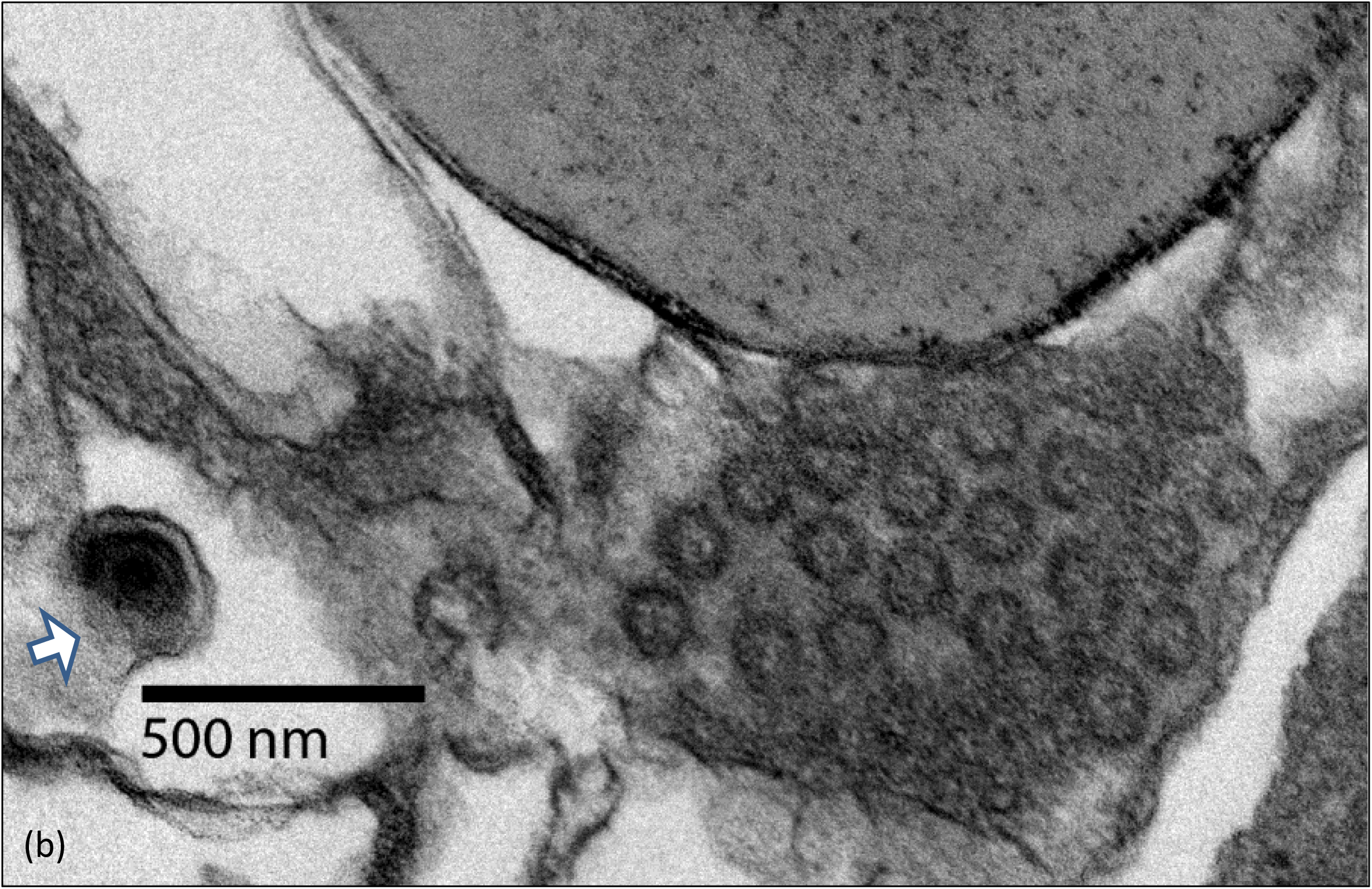
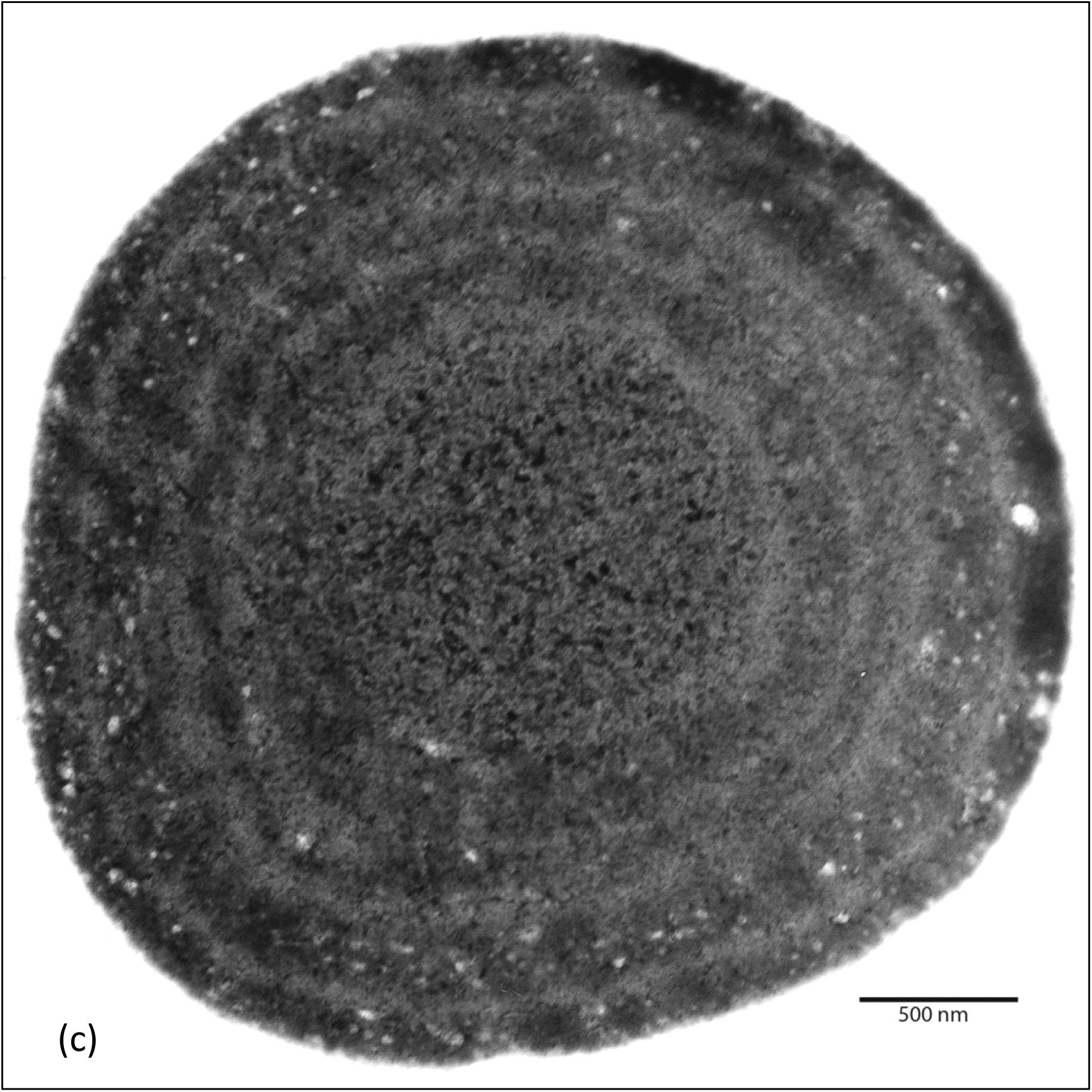
The granular fibrillary VLP mechanism is so flexible inclusions of almost any description can form. The significances are unknown. Microorganisms with these functions are new to science, and they are associated with neurodegenerative infections. Common type of inclusion, similar to those of the scrapie agent Structures appearing as ‘pseudoviral hollow-cored vesicles’ were found (f) along side a budding membrane bound form (arrow) indicating the remarkable variability of products produced. Large circular arrays can also form within SMCA-infected Cells

Following classifying SMCA as a spiroplasma, Bastian reported detecting spiral shapes in biopsy material from a CJD case (38, 39) and publishes that spiroplasma cause TSE (40, 41, 42). But Liberski and Jeffrey, who also show spiral shapes in TSE cases, conclude “Thus spiroplasma-like inclusions have no connection to real “spiroplasmas”, the cell-wall free prokaryotes taxonomically belonging to the class *Mollicutes*. The notion that spiroplasma have no role to play in TSEs is also supported by the lack of any antibodies directed against several spiroplasma species in CJD patients” (43).

In 2007 Bastian et al. reported recovering spiroplasma from TSE cases and grew them until plentiful, though none were confirmed (44). The authors specified that injecting *S.murim*, strain SMCA, into brains of neonatal deer triggered CWD clinical signs (documented by video) and caused spongiform pathology. But as Bastian and Feng communicate “it is noteworthy that *S. mirum* has induced formation of alpha-synuclein (the amyloid associated with Parkinson’s disease in mammalian tissue cultures [48])…” (45). However, CWD is a prion disease and is not caused by α-synuclein amyloid alone, leaving the CWD report perplexing. Other investigators show that SMCA does not cause TSE. Hamir et al. injected SMCA into brains of raccoons, animals susceptible to TSE without causing clinical signs or spongiform encephalopathy (46). Goats are highly susceptible to TSE. French et al. establish that SMCA does not cause TSE when injected into brains of goats (47). This leaves little evidence for reports that SMCA causes TSE.

Bastian’s US patent 7,888,039 B2: 2011, “Assay for Transmissible Spongiform Encephalopathies” is a reliable resource to test his spiroplasma data. When bacteria cause systemic infections they usually elicit immune responses of diagnostic significance. If spiroplasmas cause TSE then spiroplasma-specific antibodies should be in sera from TSE cases. Bastian reports his patent detects spiroplasma specific antibodies in sera from TSE cases, at high specificity and sensitivity (8, 48, and 49). His test uses the spiroplasma Hsp60 antigen described as cloned from *S. mirum*, strain SMCA (48), but by sequence, (GenBank CP006720 [*S. mirum* ATCC 29335]) and PCR (Fig), SMCA lacks this gene. The patent incorporated Hsp60 sequence shares 100% identity with *S. citri*. Furthermore, the microorganisms recovered from TSE affected sheep are also PCR negative for spiroplasma Hsp60 (Fig. 3, b). These prion+ sheep with scrapie would test TSE free by Bastian’s spiroplasma assay.

Bastian et al. describe spiroplasma biofilms as “breakthrough research” (48, 8, 9, and 45). But biofilms are produced after organisms bind to surfaces, and are not produced when freely growing in solution. By ignoring culture controls (48), it is hard to differentiate a biofilm from an artifact (Fig. 2, m). The “microbial hedgehogs”, formed by a scrapie isolate, produce and use biofilm-like materials when freely growing in broth for purposes distinct from biofilms (Fig. 2, h-k).

The four theories above, ‘slow viruses’, VLP, TVS and spiroplasma show resemblances to structures produced by SMCA and isolates from sheep with TSE. The biopsy examples shown prominently reveal VLP and pathology of nuclei opening as continuous with the cytoplasm and mitochondrial destruction. Spiral forms comparable to those of spiroplasma are detected but these forms do not propagate in spiroplasma broth (Fig. 2, a and 5,a). The growth forms in broth are bizarre, far different from those known as produced by microorganisms. The end products of scrapie isolate cell free growth are also unique to the literature (Fig. 2, i-k). Prokaryotes typically possess a 16s rDNA gene. Bastian and Foster report detecting16s rDNA of spiroplasma in tissues from TSE cases (50). Our tests show that SMCA and scrapie isolates have this 16s rDNA gene (Fig 3, a). Sokolova established this gene is most homologous to the 16s rDNA gene of spiroplasma from deer flies (51). However, incorporating genes from other organisms is common in microorganisms, animals and humans without defining the species. The structures and functions of SMCA and scrapie isolates shown differ from known microorganisms. They lack the spiroplasma Hsp60 gene (Fig. 3, b) described as present in all members of the genus spiroplasma (US patent 7,888,039 B2: 2011).

#### The Prion Theory of “Proteinaceous Infectious Particles”

The origin of prions as “proteinaceous infectious particles” arose from studies of scrapie, an infectious neurodegenerative disease affecting sheep. Failure to recover pathogens such as viruses, bacteria, fungi or protozoa led to testing the agent’s physical properties. Alper et al.’s impeccable radiation data are cited as the origin of prion theory (52). Following radiation of brain tissues from affected sheep she showed that scrapie clinical signs and spongiform pathology were transferable by injections into brains of normal sheep, indicating the cause propagated without nucleic acids, a heretical concept.

Griffiths suggested Alper’s agent was a protein that could self-replicate via two stable conformations, one normal and one infectious. The infectious conform would cause disease but would also multiply by binding to the normal conform causing its conversion into the infectious conformation (53). Prusiner named Griffith’s proposed infectious protein a prion defined as a “proteinaceous infectious particle” and set forth to purify prions using a hamster scrapie model, as documented in Taube’s investigative report (54). What he purified was not the expected foreign agent, but the β-sheet conform of a normal hamster protein which was also named a prion. This normal prion conform is encoded in mammalian genomes. Its miss-folded conform causes disease in diverse species (55). In 1997 Prusiner received The Nobel Prize “for his discovery of Prions -a new biological principle of infection” (56). The question is, did Prusiner discover infectious proteins, or did he uncover a new biological principle that the conforms of some proteins express their own functions and can replicate? Differences in these data interpretations can be traced to Prusiner choosing the Syrian golden hamster scrapie model rather than natural scrapie (57). Data from lab animal models always reflect the model but not necessarily naturally caused diseases.

The hamster scrapie surrogate was developed by injecting extracts from brains of TSE affected goats into the brains of Chandler’s mice and then into rat brains, and then into hamster brains, where the incubation time was reduced (58). Due to receptor and sensing differences, surviving through multiple species barriers is improbable for a pathogen. Evidence by David-Ferreira (59) implies the scrapie pathogen lost control of its genetic expression and died prior to reaching hamsters. Perhaps the conserved prion β-sheet conform crossed species barriers by serial brain injections. If so, hamster scrapie simulates the prion pathogenic mechanism but not natural transmission. For example, being contagious is one mechanism of natural scrapie transmission. Morales et al. (60) report the “lack of prion transmission by sexual or parental routes in experimentally infected hamsters”. Alexeeva et al. failed to detect 16s rDNA in the hamster model (61) yet this gene was reported brain tissues of TSE cases by Bastian and Foster (50) and is found in isolates from sheep with scrapie (Fig. 3 a). This gene and the isolates from sheep with scrapie are clues to differentiating hamster scrapie from naturally acquired scrapie and other TSE diseases.

Today the prion label refers to normal proteins with two stable conformations, each expressing different functions. The normal conform is α-helices rich, but when triggered to fold into its ancillary conform, the α-helices decrease while the β-sheets increase, become predominant, drastically changing the prion’s function (62, 63, 64). Once the normal prion conform is caused to fold into its pathogenic conform, even by binding to steel wires (64), it will self-amplify by converting normal prions into the pathogenic conform, an autocatalytic process, whether injected into brains or vials containing normal prion conforms (15). The prion mechanism is also found in other diseases including MSA, and in simple organisms as fungi and a bacterial species (65, 66, 67, 68, and 69). These findings establish prion mechanisms as a newly recognized biological principle explaining how a protein (or at least a protein’s conform) can increase its numbers without the need for nucleic acids. Most experiments designed to support prions as infectious agents end by brain injections (70) which cannot differentiate an infectious agent from the prion autocatalytic pathogenic mechanism.

Since the β-sheet conform alters a normal mammalian protein it is a toxin. Consumption or other extensive exposure to toxins can cause neurodegeneration by triggering a prion mechanism, though toxins themselves are not infectious causes. The CJD variant is an example of this (71). In addition, excessive exposure to neurotoxins such as paraquat and some agricultural insecticides can cause Parkinson’s disease. This is most likely caused by the toxins triggering the misfolding of prion type proteins such as α-synuclein, which results in the pathogenic cascade leading to disease (72, 73, and 74). The concept of infectious proteins is not essential to interpret prion data; perhaps they are not the naturally transmitted infectious causes.

Basing a theory on negative data, such as the failure to recover pathogens from scrapie affected sheep, is risky. Many microorganisms modify host proteins as pathogenic mechanisms. Just one example of a pathogen that triggers the prion mechanism should end the theory of infectious proteins, but the fundamental importance of the prion biological principle remains as a major accomplishment. According to Prusiner both Scrapie and MSA are prion diseases (75). Establishing that SMCA induces α-synuclein aggregations and results in MSA type Lewy bodies should be further researched, along with determining if isolates form scrapie affected sheep cause prion aggregations. Determining if each agent can cause either aggregations of α-synuclein (SMCA) or prion amyloid (isolates from sheep with scrapie) using relevant cell cultures is the easiest beginning for additional research, as preliminary results are encouraging (7, 8 and 9).

#### Links to MS and the α-synucleinopathies

For several decades prior to prion theory diverse neurodegenerative diseases were thought to be caused by similar pathogens. For example, during the 1960s scrapie afflicting sheep and MS affecting humans were studied as caused by related agents. As E. J. Field explained “The link between Multiple Sclerosis (MS) and scrapie rests at present on two sets of experiments, one in Iceland (Palsson et al., 1965) and one in Newcastle (Field, 1965). In both cases scrapie emerged in animals inoculated with multiple sclerosis material (sheep in Iceland; mice in Newcastle) and in both cases incubation period shortened and signs became more typical with passage—a phenomenon to be expected when the “species barrier” is crossed” (76). Though corroborative biopsy data from the 1970s strengthened Field’s view (19, 20), the connection shattered when prions were accepted as the infectious cause of scrapie, but not MS. Yet the pre-prion concept that similar pathogens could cause dissimilar diseases remains viable as the prion mechanism is more clearly understood. With recovering of agents from sheep with scrapie sheep that share similarities with TSE and MS biopsy studies Field’s observations deserve further consideration.

Similarities between the MS biopsy from Narang and Field (20) and the data presented include the fibrillary VLP network with VLP of two different sizes (Fig 1 d, e, h), opening of nuclear membranes leading to continuity between nuclear and cytoplasmic contents (Fig 1 c, d) and similar patterns of mitochondria destruction (Fig 1 o, p). Because the process of forming “microbial hedgehogs” would be difficult to evolve, they likely are important for the pathogen’s transmission. These extensive correlations are not causations but reasons to test further. If the correlations hold true then Tanaka et al’s. finding an inclusion in an MS leukocyte found only when outside the patient (77) is a reason to test if leukocytes during episodes (and CNS tissues) would reveal 16s rDNA but not Hsp60, paralleling SMCA and scrapie isolates. According to Brown (78) MS distribution correlates with tick-borne diseases. If correct the pathogen’s genetic markers should be within blood during episodes in addition to disease relevant areas of the CNS. Until cures are realized all possibilities of infectious causes should be tested. Because MS does not use a prion type pathogenic mechanism, and the above conjectures hold, then MS is projected as the first neurodegenerative disease to be cured, by killing its infectious cause.

Though there are several known causes of Parkinson’s including genetic errors and exposure to toxins, in most cases the etiology is unknown. Braak proposed pathogens traveling along neurons as likely cause of idiopathic Parkinson’s disease (79). The fibrillary strands continuous with VLP, produced by infecting cells with SMCA, appear to unravel enabling entry into tubes (Fig 4 g, h, i), findings consistent with Braak’s insight. Identifying inclusions comparable to Lewy bodies (Fig 7) led to test for α-synuclein aggregations, a defining feature the α-synucleinopathies (80).

**Figure 7.**
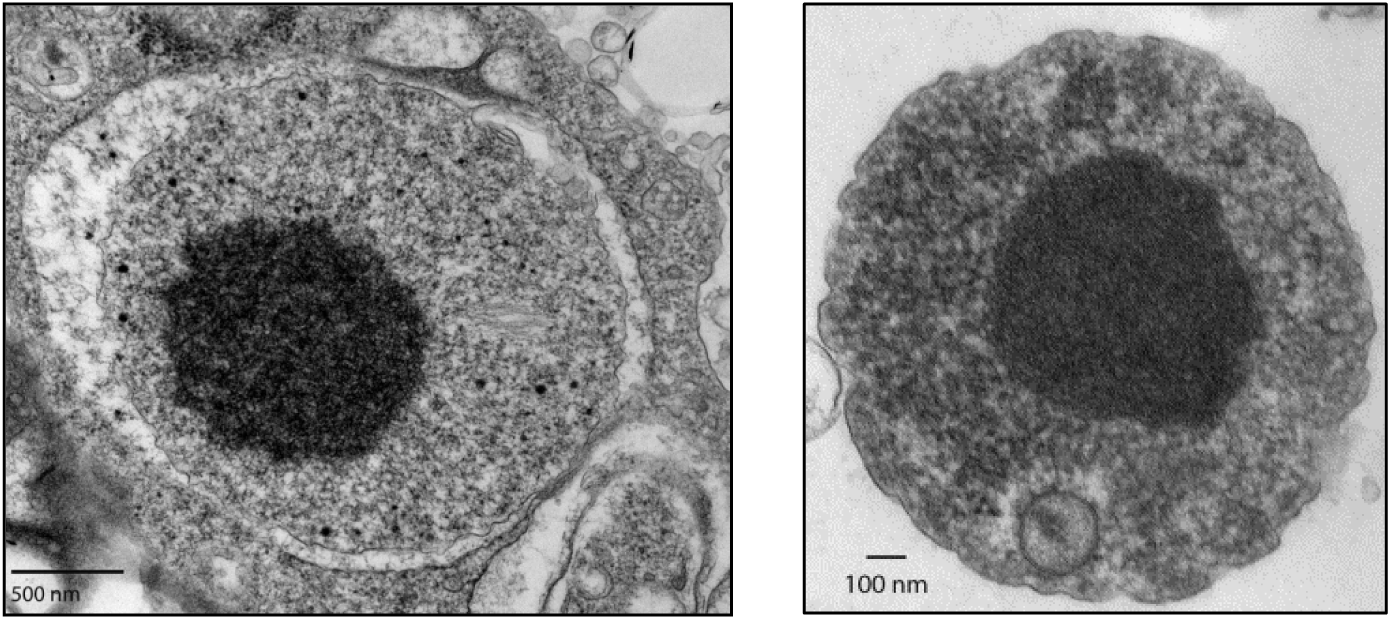
Not all inclusions that look like Lewy bodies are Lewy bodies. The inclusions above were found in BCE infected by scrapie isolates. They test negative for α-synuclein aggregations. They were found in cells infected with an microorganism recovered from sheep with scrapie. Could they be associated with prion amyloid?

Reports that aggregations α-synuclein form in SMCA infected BCE and N2a cells are encouraging (7, 8 and 9). The double limiting membrane surrounding the Lewy body type inclusion (Figure 4 l), detection of tight junctions and EM observations indicate these inclusions form in infected cell nuclei, consistent with MSA (81, 82), supporting Braak’s theory that pathogens could cause α-synucleinopathies which also include Lewy body dementia and Parkinson’s.

Remarkably, the multiplication rate of N2a cultures, separately receiving culture fluids from BCE, infected with either scrapie isolates or SMCA, but not BCE alone, rapidly accelerated, yielding a 50% increase in packed cell volume over five to seven days. This result supports prior studies by Kotani et al. using SMCA (83) and Oleszak et al. and Manuelidis et al. with CJD extracts (84, 85), indicating they induce cell proliferation, a phenomenon always of interest.

A surprising find was a Lewy body type inclusion in cells infected with a scrapie isolate. Since it is delineated by a single membrane (Fig 7), rather than the double membrane of similar bodies in the same cell line, it is not from the cell line. Could it play a role in triggering the prion miss-folding process, analogous to SMCA-infected cells that accumulate α-synuclein amyloid?

To aid further studies, four BCE cell lines each persistently infected with an isolate from different prion positive sheep with scrapie, were received by the Biodefense and Emerging Infections Research Resources Repository on May 22, 2013, for preservation and release to qualified investigators. 10801 University Boulevard, Manassas, VA 20110-4137. www.beiresources.org. They were deposited by LSU and are identified as BCE infected with scrapie, and each identified by the scrapie sheep case number provided by the Caine Veterinary Teaching Center, University of Idaho, and Caldwell Idaho 83607. The case numbers are #296, #301, #5061, and #6000. From three of these isolates cell-free forms can easily be recovered by gradual transitioning infected cell culture fluids into SP-4 medium.

## Abbreviations

CJD: Creutzfelt-Jacob Disease
CNS: central nervous system
MS: multiple sclerosis
MSA: multiple systems atrophy
SMCA: suckling mouse cataract agent
TSE: transmissible spongiform encephalopathy
TVS: tubulovesicular structures
VLP: virus like particles

## Acknowledgements

Eyes from scrapie sheep were purchased using an in-house LSU AgCenter grant to F. O. Bastian and W. J. Todd to determine if spiroplasma could cause TSE. Funding for the deposited scrapie isolations and cell-free growth data were from Margret Pahl Stewart Foundation and LSU colleague Gregg Henderson. Funding in kind includes trading lab, finances and tenure for bench space in the lab of Ron Thune, Head, PBS, SVM and access to SVM facilities where this work was completed. Olga Borksenious provided EM assistance during the beginning of this work. Helpfulness of C. Mitch Boudreaux and Sue Hagius of the LSU AgCenter and Angelina Demming, Greg McCormick and Thomas Gillis of the National Hansen’s Disease Program at the LSU SVM are greatly appreciated. The antibodies to SMCA were provided from the Department of Veterinary Science LSU AgCenter. Dominique Toumadje (MID-BRR Project Coordinator) is appreciated for helping secure the scrapie isolates. This manuscript was patiently edited by Susan Todd, Executive Director of 504HealthNet, New Orleans.

## Summary and Conclusions

This research uncovers a group of microorganisms that expand our concepts of microbial diversity to include those expressing two separate sets of structures and functions. When intracellular they use a virus-like process based on flexible granular fibrillary strands which often condense and appear as virus-like particles (VLP). The litheness of this mechanism allows formation of many structures and inclusion bodies similar to those found in biopsy material from cases of CNS neurodegeneration, but they are of unverified utility. When independent of host cells the two microorganisms studied propagate as fragile prokaryotes not previously described. These microorganisms recovered from scrapie were studied further, producing surprising data. When cultured to high concentrations these fragile microorganisms disintegrate. The released constituents self-assemble, forming large highly organized structures named ‘microbial hedgehogs’, which are hypothesized as structures to protect the pathogens genome during long periods between hosts; they are infectious to mammalian cells. All microorganisms studied are available for testing, and more research needs to be done. It is unknown if microorganisms with the functions described cause diseases in nature. Welcome to the magical microbial world where their reality exceeds our expectations.

## Author’s responsibilities

William J. Todd planned this study, recovered and established persistently infected cell cultures and cell free cultures of agents from scrapie affected sheep, and wrote the manuscript. Lidiya Dubytska designed and ran the PCR tests revealing genes key to understanding features of SMCA and scrapie isolates. Peter J. Mottram’s microscopy using a scrapie agent and Xiaochu Wu’s microscopy with FA labeled SMCA revealed unique features of these agents. Yuliya Y. Sokolova shared the EM specimen preparations and screening of EM images.

